# Improved polygenic prediction by Bayesian multiple regression on summary statistics

**DOI:** 10.1101/522961

**Authors:** Luke R. Lloyd-Jones, Jian Zeng, Julia Sidorenko, Loïc Yengo, Gerhard Moser, Kathryn E. Kemper, Huanwei Wang, Zhili Zheng, Reedik Magi, Tonu Esko, Andres Metspalu, Naomi R. Wray, Michael E. Goddard, Jian Yang, Peter M. Visscher

## Abstract

The capacity to accurately predict an individual’s phenotype from their DNA sequence is one of the great promises of genomics and precision medicine. Recently, Bayesian methods for generating polygenic predictors have been successfully applied in human genomics but require the individual level data, which are often limited in their access due to privacy or logistical concerns, and are computationally very intensive. This has motivated methodological frameworks that utilise publicly available genome-wide association studies (GWAS) summary data, which now for some traits include results from greater than a million individuals. In this study, we extend the established summary statistics methodological framework to include a class of point-normal mixture prior Bayesian regression models, which have been shown to generate optimal genetic predictions and can perform heritability estimation, variant mapping and estimate the distribution of the genetic effects. In a wide range of simulations and cross-validation using 10 real quantitative traits and 1.1 million variants on 350,000 individuals from the UK Biobank (UKB), we establish that our summary based method, SBayesR, performs similarly to methods that use the individual level data and outperforms other state-of-the-art summary statistics methods in terms of prediction accuracy and heritability estimation at a fraction of the computational resources. We generate polygenic predictors for body mass index and height in two independent data sets and show that by exploiting summary statistics on 1.1 million variants from the largest GWAS meta-analysis (*n ≈* 700, 000) that the SBayesR prediction *R*^2^ improved on average across traits by 6.8% relative to that estimated from an individual-level data BayesR analysis of data from the UKB (*n ≈* 450, 000). Compared with commonly used state-of-the-art summary-based methods, SBayesR improved the prediction *R*^2^ by 4.1% relative to LDpred and by 28.7% relative to clumping and *p*-value thresholding. SBayesR gave comparable prediction accuracy to the recent RSS method, which has a similar model, but at a computational time that is two orders of magnitude smaller. The methodology is implemented in a very efficient and user-friendly software tool titled GCTB.

## Introduction

The capacity to accurately predict an individual’s phenotype from their DNA sequence is one of the great promises of genomics and precision medicine^1–5^, recognising that the accuracy of a genetic risk predictor is dependent on the genetic contribution to variation in the trait. It is anticipated that genetic risk prediction will be useful for informing early disease intervention and aiding diagnosis by identifying individuals with an increased genetic risk of disease^5–7^. Accurate genetic predictors for complex traits and disorders are currently limited, due mainly to an incomplete understanding of complex genetic variation, small training sample sizes and suboptimal modelling^4,8,9^. Through large consortia and biobank initiatives, sample sizes for genome-wide association studies (GWASs) are reaching a critical point, now for some traits greater than a million individuals, at which, and under optimal modelling conditions, the predictors generated 3 could approach their maximum (from theory) prediction accuracy for some traits^10–13^.

One common approach for generating polygenic predictions uses GWASs effect size estimates derived from simple linear regression applied to each single-nucleotide polymorphism (SNP) independently across the genome, and uses a linear combination of the estimated effects and allele counts at genetic markers, chosen via marker pruning coupled with *p*-value thresholding^14–17^. Although simple to implement and useful, this method has been shown to provide suboptimal predictions with the best estimate of each marker’s effect requiring the effects to be treated as random^18–20^. In this work, we will restrict the term polygenic risk score to those predictors generated from using simple linear regression and use the term estimated genetic value (EGV) for the general concept of generating a polygenic predictor from SNP data. Linear mixed model (LMM) methodologies have been successfully applied in human genetics^21–25^ and are derived under the multiple regression model. These methods jointly analyse all SNPs, which accounts for linkage disequilibrium (LD) between markers capturing the maximum amount of variation at a genetic locus especially if multiple causal variants colocalise. Bayesian extensions of the standard LMM, which assumes a single normal distribution on the genetic effects, have been made to include alternative prior distributions for the genetic effects that deviate from the assumptions of the infinitesimal model, and were pioneered in plant and animal breeding^26–30^. Recent implementations of Bayesian multiple regression methodology require access to the individual level data^29,31^ and currently do not scale well computationally to sample sizes of greater than half a million individuals and millions of genetic variants.

The inability to access individual level genetic and phenotypic data has motivated methodological frameworks that only require publicly available summary data^9^. Summary statistics methodology now covers the gamut of statistical genetics analyses including: effect size distribution estimation^32,33^, joint SNP association analysis and fine mapping^34,35^, allele frequency and association statistic imputation^36–38^, heritability and genetic correlation estimation^39–43^ and polygenic prediction^44–46^. These methods require GWAS summary data, which typically include the estimated univariate effect, standard error, sample size and allele frequency, and an estimate of LD among genetic markers, which are easily accessed via public databases.

In this work, we extend the established summary statistics methodological framework through the utilisation of a likelihood that connects the multiple regression coefficients with the summary statistics from GWAS (similar to Zhu and Stephens^42^). We perform Bayesian posterior inference through the combination of this likelihood and a finite mixture of normal distributions prior on the markers effects, which encompasses the models proposed in Habier *et al.*^27^, Erbe *et al.*^28^ and Moser *et al.*^31^. Here, we focus on optimising prediction accuracy but the methodology is capable of simultaneously estimating SNP-based heritability 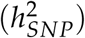, marker mapping and estimating the distribution of marker effects. We maximise computational efficiency by taking advantage of LD matrix sparsity and, importantly, once the GWAS effect size estimates have been generated the computational time of our method is independent of sample size making the method applicable to an arbitrary number of individuals.

We establish that our summary-based method, SBayesR, outperforms other state-of-the-art summary statistics methods in terms of prediction accuracy and 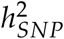 estimation in a wide range of simulations using real genotype data from 350,000 unrelated individuals of European ancestry from the UK Biobank (UKB). The state-of-the-art summary statistics methods used for comparison include those that seek to estimate posterior mean effect sizes from GWAS summary statistics by assuming a prior for the genetics effects and LD information from a reference panel stored for each chromosome in a block diagonal form or constructed from an LD matrix shrinkage estimator. Specifically, we compare with LDpred^44^, which assumes a point-normal mixture prior for the genetics effects and a block-diagonal LD matrix, summary best linear unbiased prediction (SBLUP)^45^, which assumes a normal distribution for the genetics effects and a block-diagonal LD matrix, Regression with Summary Statistics (RSS)^42^, which has a class of priors for the genetic effects to select from but we compare against the mixture of two normal distributions prior^29^ and is optimised for the use of a shrunk LD matrix^36^. We further compare with clumping and then *p*-value thresholding (P+T) implemented in the PLINK 2 software^47^ and the individual data implementation of the BayesR model^31^, which assumes a finite mixture or normal distributions (including a point mass at zero) prior on the genetic effects and has been optimised for time and memory efficiency. For 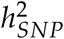 estimation comparison we use the widely used summary data LD score regression (LDSC) method^39^, which relies on the expected relationship between, under a polygenic model, per variant chi-squared summary statistics and LD scores from a reference, RSS, which can estimate 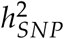 given the posterior mean of the genetics effects and the individual data Haseman-Elston regression (HEreg) method^48^, which relies on identity by state relatedness measures derived from a genetic relatedness matrix and the cross product of the phenotypes for pairwise individuals and is efficient on large data sets.

We show that SBayesR performs similarly in terms of prediction accuracy to individual data methods and outperforms other state-of-the-art summary methods in five-fold cross-validation with 1.1 million HapMap 3 (HM3) variants and 10 real quantitative traits from the UKB. We further perform large-scale analyses for height and body mass index using 1.1 million HM3 variants and the full UKB European ancestry (both related and unrelated individuals) data set and predict into two independent samples from the Health and Retirement Study (HRS) and the Estonian Biobank (ESTB). In these across biobank analyses, we show that by exploiting summary statistics from the largest GWAS meta-analysis (*n ≈* 700, 000) on height and body mass index^49^ that on average across traits the SBayesR prediction accuracy improved by 6.8% relative to that estimated from an individual-level data BayesR analysis of data from the UKB (*n ≈* 450, 000). Compared with commonly used state-of-the-art summary-based methods, SBayesR improved the prediction *R*^2^ by 4.1% relative to LDpred and by 28.7% relative to clumping and *p*-value thresholding. SBayesR gave comparable prediction accuracy to the recent RSS method, which has a similar algorithm, but at a computational time that is two orders of magnitude smaller. The methodology is implemented in a very efficient and user-friendly software tool titled GCTB^30^.

## Materials and Methods

### Data

#### UK Biobank

We used real genotype and phenotype data from the full release of the UK Biobank (UKB). The UKB is a prospective community cohort of over 500,000 individuals from across the United Kingdom and contains extensive phenotypic and genotypic information about its participants^50^. The UKB data contains genotypes for 488,377 individuals (including related individuals) that passed sample quality control (99.9% of total samples). A subset of 456,426 European ancestry individuals was selected using the protocol described in Yengo *et al.*^49^. To exclude related individuals, a genomic relationship matrix (GRM) was constructed with 1,123,943 HM3 variants further filtered for minor allele frequency (MAF) > 0.01, pHWE < 10^−6^ and missingness < 0.05 in the European subset, resulting in a final set of 348,580 unrelated (absolute GRM off-diagonal < 0.05) Europeans. Genotype data were imputed to the Haplotype reference consortium and UK10K panel, which was provided as part of the data release and described in^50^, and contained SNPs, short indels and large structural variants. Variant quality control included: removal of multi-allelic variants, SNPs with imputation info score < 0.3, retained SNPs with hard-call genotypes with > 0.9 probability, removed variants with minor allele count (MAC) ≤ 5, Hardy-Weinberg *p*-value (pHWE) < 10^−5^ and removed variants with missingness > 0.05, which resulted in 46,500,935 SNPs for the 456,426 individuals.

#### Atherosclerosis Risk in Communities, 1000 Genomes and UK10K data

The implemented summary statistics methodology requires an estimate of LD among genetic markers. In addition to the UKB, three data sets were used to calculate LD reference matrices. We used the genotype data from the Atherosclerosis Risk in Communities (ARIC)^51^ and GENEVA Diabetes study obtained via dbGaP. The ARIC+GENEVA data consisted of 12,942 unrelated individuals determined by an absolute GRM off-diagonal relatedness cutoff of < 0.05. After imputation to the Phase 3 of the 1000 Genomes Project (1000G)^52^, 1,182,558 HM3 SNPs (MAF > 0.01) were selected and available for analysis after quality control. Whole-genome sequencing data from the 1000G project was used for LD matrix reference calculation. These data were subsetted to a set of 397 individuals with European ancestry to be consistent with the LD reference used in Zhu and Stephens^42^. Whole-genome sequencing data from the UK10K project^53^ was also used for analysis. The UK10K contains 17.6 million genetic variants (excluding singletons and doubletons) in 3,642 unrelated individuals after quality control, which was performed as per Yang *et al.*^54^.

#### Health and Retirement Study and Estonian Biobank

For out-of-sample validation of genetic predictors we used two cohorts that are independent of the UKB. We used genotypes imputed to the 1000G reference panel and phenotypes from 8,552 unrelated (absolute GRM off-diagonal < 0.05) participants of the Health and Retirement Study (HRS)^55^. After imputation and restricting variants with an imputation quality score > 0.3, MAF > 0.01 and a pHWE > 10^−6^ there were 24,777,992 SNPs available for prediction. The Estonian Biobank^56^ is a cohort study of over 50,000 individuals over 18 years of age with phenotypic and genotypic data. For the prediction analysis we used data from 32,594 individuals genotyped on the Global Screening Array. These data were imputed to the Estonian reference^57^, created from the whole genome sequence data of 2,244 participants. Markers with imputation quality score > 0.3 were selected leaving a total of 11,130,313 SNPs for prediction.

### Overview of summary statistics based Bayesian multiple regression

We relate the phenotype to the set of genetic variants under the multiple linear regression model

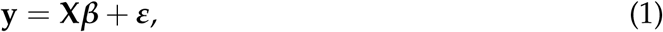

where **y** is an *n* × 1 vector of trait phenotypes, which has been centred, **X** is an *n* × *p* matrix of genotypes coded as 0, 1 or 2 representing the number of copies of the reference allele at each marker, ***β*** is a *p* × 1 vector of multiple regression coefficients (marker effects) and ***e*** is the error term (*n* × 1). We can relate the multiple regression model to the estimates of the regression coefficients from *p* simple linear regressions **b** from GWAS, by multiplying (1) by **D**^−1^**X**′ where 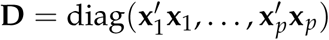 to arrive at

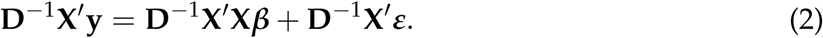

Noting that **b** = **D**^−1^**X**′**y** is the vector (*p* × 1) of least-squares marginal regression effect estimates and the correlation matrix between all genetic markers 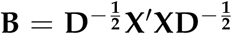, we rewrite the multiple regression model as

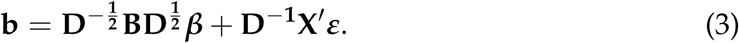

Assuming *ε*_1_, …, *ε_n_* are independent 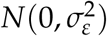, the following likelihood can be proposed for the multiple regression coefficients ***β***

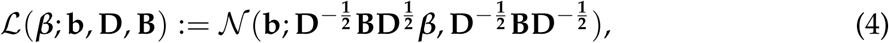

where 𝒩 (***ξ***; ***µ***, **Σ**) represents the multivariate normal distribution with mean vector ***µ*** and covariance matrix **Σ** for ***ξ***. If individual level data are available then inference about ***β*** can be obtained by replacing **D** and **B** with estimates 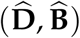 from the individual level data. If individual level data are unavailable then we can replace **D** with 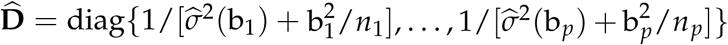, where 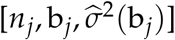 are the sample size used to compute the simple linear regression coefficient, an estimate of the simple linear regression allele effect coefficient and 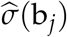 the standard error of the effect for the *j*th variant respectively. This reconstruction of 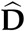 assumes that the markers have been centred to mean 0 (please see the Supplemental Note for a detailed reasoning of this reconstruction of 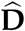). If we make the further assumption that the genetic markers have been scaled to unit variance then we can replace **D** with 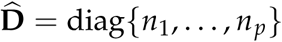. Similarly, we replace **B**, the LD correlation matrix between the genotypes at all markers in the population, which the genotypes in the sample are assumed to be a random sample, with 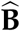 an estimate calculated from a population reference that is assumed to closely resemble the sample used to generate the GWAS summary statistics. Zhu and Stephens^42^ discuss further the theoretical properties of a similar likelihood. We assess the limits of replacing **D** and **B** with these approximations through simulation and real data analysis.

We perform Bayesian posterior inference by assuming a prior on the multiple regression genetic effects and the posterior

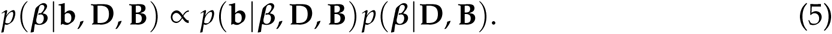

In this paper we implement the BayesR model^28,31^, which assumes that

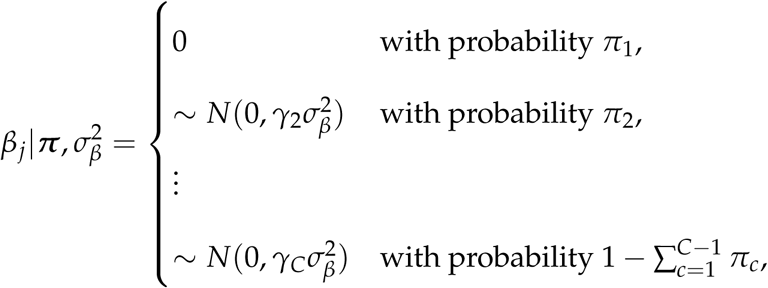

where *C* denotes the maximum number of components in the finite mixture model, which is prespecified. The *γ_c_* coefficients are prespecified and constrain how the common marker effect variance 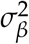 scales in each distribution. In previous implementations of BayesR the variance weights ***γ*** were with respect to the genetic variance 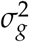. For example, it is common in the BayesR model to assume *C* = 4 such that ***γ*** = (*γ*_1_, *γ*_2_, *γ*_3_, *γ*_4_)′ = (0, 0.0001, 0.001, 0.01)′. This requires the genotypes to be centred and scaled and equates 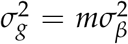, where *m* is the number of variants. We relax this assumption to disentangle the relationship between these parameters and to maintain the flexibility of the model to assume scaled or unscaled genotypes. In this implementation, we let the weights be with respect to 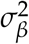 and have a default ***γ*** = (0, 0.01, 0.1, 1.0)′, which maintains the relative magnitude of the variance classes as in the original model. The Supplementary Note details further the hierarchical model and hyperparameter prior specification. The Supplementary Note also details the derivation of the Markov chain Monte Carlo Gibbs sampling routine for sampling of the key model parameters 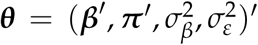 from their full conditional distributions. SNP-based heritability estimation is performed by calculating 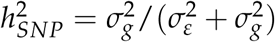, where the genetic variance 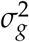 is calculated as Var(**X*β***) for each sampled set of ***β***^(*i*)^ in iteration *i* of the MCMC chain (see Supplemental Note for further details).

To illustrate why the Gibbs sampling routine proposed lends itself to the use of summary statistics, we focus on the full conditional distribution of *β_j_* under the proposed multiple regression model. To facilitate the explanation we make the simplifying assumption that *C* = 2 and ***γ*** = (*γ*_1_, *γ*_2_) = (0, 1). The full conditional distribution of *β_j_* under this assumption (see Supplemental Note) is

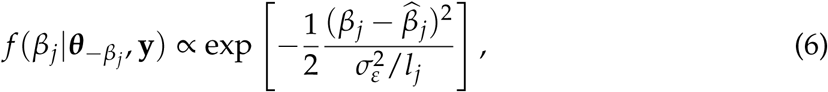

where 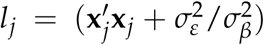 and 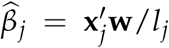. The term *l_j_* only involves the diagonal elements of **X′X** and is easily calculated from summary statistics via 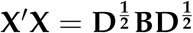. For 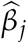, we require 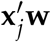, which is defined as

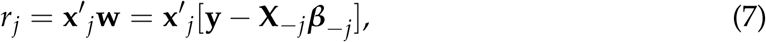

where **X**_*-j*_ is **X** without the *j*th column. This quantity can be efficiently stored and calculated in each MCMC iteration via a right-hand side updating scheme. We define the right-hand side **X**′**y** corrected for all current ***β*** as

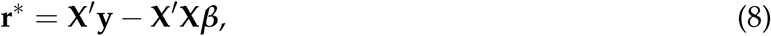

where **r**^*^ is a vector of dimension *p* × 1. The *j*th element of **r**^*^ can be used to calculate

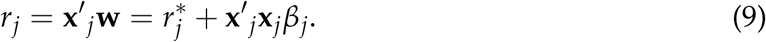

Therefore, once a variant has been chosen to be in the model its effect is sampled from (6), which is the kernel of the normal distribution with mean 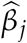 and variance 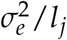 (see the Supplemental Note for more detail). After the effect for variant *j* has been sampled we update

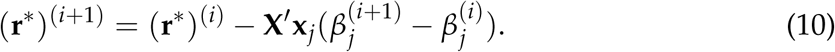

Importantly, after the initial reconstruction of **X**′**y** = **Db** from summary statistics, equation (10) only requires **X**′**x**_*j*_, which is the *j*th column of **X**′**X**. The operation in (10) is a very efficient vector subtraction and only requires the subtraction of the non-zero elements of the shrinkage estimator of the LD correlation matrix from Wen and Stephens^36^, which we perform using sparse matrix operations. The other elements of the Gibbs sampling routine are the same as the individual data model except for the sampling of 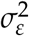, which is outlined in the Supplemental Note.

### Genome-wide simulation study

Before performing simulations using genome-wide variants, we first thoroughly tested and compared individual level and summary statistics based methods using a simulation study on two chromosomes (Supplemental Note and Figures S1, S2, S3 and S4). This small-scale simulation established the implementation of the method by comparing the individual data BayesR method with SBayesR using the full LD matrix constructed from the cohort used to perform the GWAS, which should theoretically give equivalent results. Furthermore, it allowed for a thorough investigation of the method’s properties as a function of genetic architecture and LD reference in reasonable computing time relative to genome-wide analyses. In particular, we observed that SBayesR outperformed other summary statistics methods when the genetic architecture of the simulated trait contained very large genetic effects and a polygenic background, which is expected due to the very flexible SBayesR prior (Supplemental Figure S3). Overall at the scale of two chromosomes, SBayesR generally outperformed other methods in terms of prediction accuracy and performed well at 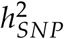 estimation.

To investigate the performance of the methodology at a genome-wide scale, we simulated quantitative phenotypes using 1,094,841 genome-wide HM3 variants and a random subset of 100,000 individuals from the 348,580 unrelated European ancestry individuals in the UKB data set. For the same set of 1,094,841 variants, we generated two independent tuning and validation genotype sets from the remaining 248,580 unrelated European individuals each containing 10,000 individuals. The 1,094,841 variant subset was formed from the 1,365,446 HM3 SNPs further filtered on MAF> 0.01, strand ambiguous SNPs (as do Vilhjálmsson *et al.*^44^ and Bulik-Sullivan *et al.*^39^), removal of long-range LD regions (defined in Bycroft *et al.*^50^ Table S13 and includes the MHC), which increased model stability across a large set of phenotypes, and overlapped with the 1000G genetic map downloaded from joepickrell/1000-genomes-genetic-maps. The 1000G genetic map is required for use in the LD matrix shrinkage estimator^36^. The genetic map files contain interpolated map positions for the CEU population generated from the 1000G OMNI arrays. The shrinkage estimator of the LD matrix^36^, shrinks the off-diagonal entries of the LD correlation matrix toward zero and is required for the Regression with Summary Statistics (RSS)^42^ and SBayesR methods.

The simulation study on two chromosomes established that the LD reference cohort from 50,000 random individuals from the UKB gave the highest prediction accuracy and lowest bias in 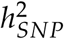 estimation (Supplemental Note). The overlap between this random subsample with the 100,000 random individuals used to generate the simulated phenotypes was 13,967. For this LD reference cohort, chromosome-wise LD matrices i.e., all inter chromosomal LD is ignored, were built and the shrinkage estimator of the LD matrix calculated using an efficient implementation in the GCTB software. The calculation of the shrunk LD matrix requires the effective population sample size, which we set to be 11,400 (as in Zhu and Stephens^42^), the sample size of the genetic map reference, which corresponds to the 183 individuals from the CEU cohort of the 1000G and the hard threshold on the shrinkage value, which we set to 10^−3^. This threshold gave a good balance between computational efficiency and accuracy with, on average, each SNP having 4,113 (SD=1,211) non-zero elements across the autosomes (Figure S5). We further stored the shrunk LD matrix in sparse matrix format (ignoring matrix elements equal to 0) for efficient SBayesR computation. For LDpred^44^, SBLUP^45^ and PLINK clumping and then *p*-value thresholding (P+T) (implemented in the PLINK 2 software^47^), a separate genotype data set is required for LD correlation reference and utilisation within each method’s program. This was set to be the same set of genotypes from 50,000 individual used to calculate the LD reference matrix for SBayesR and RSS.

Two genetic architecture scenarios were generated: 10,000 causal variants sampled under the SBayesR model i.e., 2500, 5000, and 2500 variants from each of 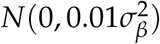, 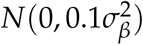, and 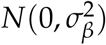 distributions respectively and 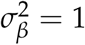. For the second architecture, 50,000 causal variants were sampled from a single standard normal distribution. For each replicate a new sample of causal variants was chosen at random from the set of 1,094,841 variants. For each scenario, 10 simulation replicates were generated under the multiple regression model using the phenotype simulation tool in the GCTA software^58^ and centred and scaled genotypes for all 100,000 individuals. For each architecture the residual variance was scaled such that the total 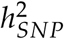 was 0.1, 0.2 and 0.5, which led to a total of six simulation scenarios.

For each of the the six scenarios, simple linear regression for each variant was run using the –`linear` option in the PLINK 2 software for each of the 10 simulation replicates to generate summary statistics. For each of the simulation scenarios the following methods were used to estimate the genetic effects: LDpred, RSS, SBLUP, P+T, BayesR^31^, and SBayesR. For 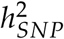 comparison we ran LD score regression (LDSC)^39^ and Haseman-Elston regression (HEreg) in the GCTA software^48,59^. HEreg requires a GRM, which was built from the 1,094,841 genome-wide HM3 variants in the GCTA software. For LDpred, we specified 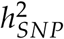 to be equal to the true simulated value, specified the number of SNPs on each side of the focal SNP for which LD should be adjusted to be 350 (approximately 1,094,841/3,000 as suggested by Vilhjálmsson *et al.*^44^), and calculated effect size estimates for all of the 10 fraction of non-zero effects pre-specified parameters, which included LDpred-inf, 1, 0.3, 0.1, 0.03, 0.01, 0.003, 0.001, 0.0003, and 0.0001. For RSS, analyses were performed for each chromosome with the chromosome-wise shrunk LD matrices calculated in GCTB and stored in MATLAB format. The RSS-BSLMM model was run for 2 million MCMC iterations with 1 million as burn in and a thinning rate of 1 in 100 to arrive at 10,000 posterior samples for each of the model parameters. For each chromosome, the posterior mean over posterior samples for the SNP effects and 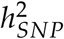 estimates was used. The chromosome wise 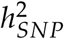 estimates were summed to get the genome-wide estimate. For SBLUP, we used the GCTA software implementation and set the shrinkage parameter 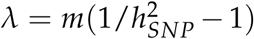 for each true simulated 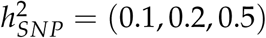 and *m* = 1, 094, 841 and the LD window size specification was set to 1 MB. LDSC was run using LD scores calculated from the 1000G Europeans provided by the software and 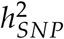 estimation performed. For P+T, we used the PLINK 2 software to clump the GWAS summary statistics discarding variants within 1 MB of and in LD *R*^2^ > 0.1 with the most associated SNP in the region. Using these clumped results, we generated polygenic risk scores for sets of SNPs at the following *p*-value thresholds: 5×10^−8^, 1×10^−6^, 1×10^−4^, 0.001, 0.01, 0.05, 0.1, 0.2, 0.5, and 1.0. BayesR was run using a mixture of four normal distributions model with distribution variance weights ***γ*** = (0, 10^−4^, 10^−3^, 10^−2^)′. BayesR was run for 4,000 iterations with 2,000 taken as burn in and a thinning rate of 1 in 10. For SBayesR, the MCMC chain was run for 4,000 iterations with 2,000 taken as burn in and a thinning rate of 1 in 10 and run with four distributions and variance weights ***γ*** = (0, 0.01, 0.1, 1)’. The posterior mean of the effects and the proportion of variance explained over the 200 posterior samples was taken as the parameter estimate for each scenario replicate for both methods.

To assess prediction accuracy, we calculated the EGV (using the score function in the PLINK 2 software) for each individual using the genotypes from the 10,000 individual tuning and validations data sets and the genetic effects estimated from each method. Parameter tuning was performed for LDpred and P+T, where for each simulation replicate the prediction accuracy was assessed for each of the pre-specified fraction of non-zero effects parameters for LDpred and the *p*-value thresholds for P+T. The parameter that gave the maximum prediction *R*^2^ in the tuning data set was then used for calculating the EGV for each individual in the validation data set. SNP effects from BayesR and SBayesR were estimated using scaled genotypes and thus each variant’s effect was divided by 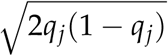, where *q_j_* is the minor allele frequency from the validation cohort of the *j*th variant, before PLINK scoring was performed. The prediction *R*^2^ was calculated via linear regression of the true simulated phenotype on that predicted from each method.

### Application to 10 quantitative traits in the UK Biobank

To assess the methodology in real data, we performed five-fold cross-validation using phenotypes and genotypes from 348,580 unrelated individuals of European ancestry from the full release of the UKB data set. We chose 10 quantitative traits including: standing height (*n*=347,106), basal metabolic rate (BMR, *n*=341,819), heel bone mineral density T-score (hBMD, *n*=197,789), forced vital capacity (FVC, *n*=317,502), body mass index (BMI, *n*=346,738), body fat percentage (BFP, *n*=341,633), forced expiratory volume in one-second (FEV, *n*=317,502), hip circumference (HC, *n*=347,231), waist-to-hip ratio (WHR, *n*=347,198) and birth weight (BW, *n*=197,778). All phenotypes were pre-adjusted for age, sex and the first ten principal components using the R programming language^60^. Principal components were calculated using high-quality genotyped variants as defined in Bycroft *et al.*^50^ that passed additional quality control filters (as applied in the European unrelated UKB data) that were LD pruned (*R*^2^<0.1) and had long-range LD regions removed (Bycroft *et al.*^50^ Table S13) leaving 137,102 SNPs for principal component calculation in the European unrelated individuals using flashPCA^61^. Following covariate correction the residuals were standardised to have mean zero and unit variance and finally rank-based inverse-normal transformed. A set of 5,000 individuals was kept separate for LDpred and P+T parameter tuning. To perform the cross-validation, the remaining 343,580 individuals were randomly partitioned into five equal sized disjoint subsamples. For each fold analysis, a single subsample was retained for validation with the remaining four subsamples used as the training data. This process was repeated five times, with each of the five subsamples used exactly once as the validation data. The SNP set used for analysis was the same set of 1,094,841 HM3 variants described in the genome-wide simulation study.

We generated summary statistics for each pre-adjusted trait in the training sample in each fold by using PLINK 2 to run simple linear regression for all variants. Using the individual level data and the summary statistics we performed analyses using the following methods: LDpred, RSS, SBLUP, P+T, BayesR, and SBayesR. For 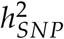 comparison we ran LDSC and HEreg. The same shrunk sparse reference LD correlation matrix from the genome-wide simulation study was used for SBayesR and RSS analyses. For LDpred, we specified the number of SNPs on each side of the focal SNP for which LD should be adjusted to be 350, and calculated effect size estimates for all of the 10 fraction of non-zero effects pre-specified parameters, which included LDpred-inf, 1, 0.3, 0.1, 0.03, 0.01, 0.003, 0.001, 0.0003, and 0.0001. The optimal parameter was chosen by predicting into the independent subset of 5,000 individuals initially partitioned off and choosing that which had the highest prediction *R*^2^ when the predicted phenotype was regressed on the true simulated phenotype. For RSS, analyses were performed for each chromosome with the chromosome-wise shrunk LD matrices calculated in GCTB and stored in MATLAB format. The RSS-BSLMM model was run for 2 million MCMC iterations with 1 million as burn in and a thinning rate of 1 in 100 to arrive at 10,000 posterior samples for each of the model parameters. For each chromosome, the posterior mean over posterior samples for the SNP effects and 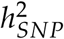 estimates was used. The chromosome wise 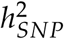 estimates were then summed to get the genome-wide estimate. For SBLUP, we used the GCTA software implementation, which requires the specification of the 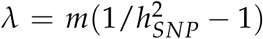 parameter. For each fold, 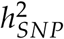 was taken to be the estimate from HEreg and *m* = 1, 094, 841. The LD window size specification was set to 1 MB for ease of computation. SBLUP and LDpred were run on each chromosome separately to improve computational efficiency. LDSC was run using LD scores from the 1000G European data and 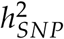 estimation performed. For P+T, we ran the same clumping procedure and calculated polygenic risk scores for the same set of *p*-value thresholds as in the simulation studies. BayesR and SBayesR were run using the same protocols as in the simulation studies. SNP effects from BayesR and SBayesR were again rescaled before PLINK scoring was performed.

To assess prediction accuracy, we calculated EGVs using the genotype data from the independent validation retained set in each fold. The PLINK 2 software was used to calculate EGVs for all methods and the prediction *R*^2^ calculated via linear regression of the true phenotype on that calculated from each method.

### Across biobank prediction analysis

To investigate how the proposed methods scale and perform in very large data sets, we analysed the full set of unrelated and related (*n* = 456, 426) UKB European ancestry individuals and used summary statistics from the largest meta-analysis of height and BMI^49^. For these analyses, the same set of 1,094,841 genome-wide HM3 variants described in the simulations was used. The set of traits was limited to those that were present in the UKB and had large independent validations sets, which included the HRS and the ESTB^56^, which contain imputed genotype and phenotype information on BMI and height.

To generate a baseline for comparison between the individual data BayesR method and the SBayesR method we first analysed data from the same set of individuals and variants from the full set of unrelated and related UKB individuals. BMI and height phenotypes were pre-adjusted for age, sex and the first ten principal components using the R programming language as per the cross-validation. We generated summary statistics for SBayesR analysis for height and BMI using a linear mixed-model to account for sample relatedness in the BOLT-LMM v2.3 software^13,25^ for the 1,094,841 HM3 variants in the full UKB data set. Using these summary statistics, we ran SBayesR for 4,000 iterations with 2,000 taken as burn in and a thinning rate of 1 in 10 and four distributions and variance weights ***γ*** = (0, 0.01, 0.1, 1)′. For comparison in the full UKB data set, we ran the individual level BayesR method using a mixture of four normal distributions model with distribution variance weights ***γ*** = (0, 10^−4^, 10^−3^, 10^−2^)′. BayesR was run for 4,000 iterations with 2,000 taken as burn in and a thinning rate of 1 in 10. The posterior mean of the sampled genetic effects and 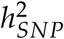 over the 200 posterior samples was taken as the parameter estimate for each trait for both methods.

Motivated by the hypothesis that summary statistics methodologies can increase prediction accuracy over large-scale individual level analyses by utilising publicly available summary statistics from very large GWASs, we took the summary statistics from the largest meta-analysis of BMI and height^49^ and analysed them using SBayesR, RSS and LDpred, which were the best performing summary based methods (in terms of prediction accuracy) in the cross-validation. We subsetted the set of 1,094,841 HM3 variants to 982,074 variants that overlapped with those in both the BMI and height summary statistics sets. The summary based methodology implicitly assumes that the summary statistics have been generated on the same set of individuals^42^. Empirically we observed that the methodology can tolerate deviations from this assumption up to a limit. To improve method convergence we removed variants from the Yengo *et al.*^49^ summary statistics that had a per variant sample size that deviated substantially from the mean of the sample size distribution over all variants, which was also performed by Pickrell *et al.*^62^ and recommended by Zhu and Stephens^42^. To minimise the variants removed, we interrogated the distributions of per variant sample size in each of the BMI and height summary statistics sets and removed variants in the lower 2.5th percentile and upper 5th percentile of the per variant sample size distribution for BMI and in the lower 5th percentile for height (Figure S6). This left 932,969 and 909,293 variants with summary information for height and BMI respectively. These sets of variants were also used in the LDpred and RSS analyses.

SBayesR was run as above with the default ***γ*** for BMI and ***γ*** = (0, 10^−4^, 10^−3^, 1)′ for height. Empirically, we observed that this constraint on the elements of ***γ*** was a further requirement for SBayesR model convergence using these height summary statistics. For LDpred, we specified the number of SNPs on each side of the focal SNP for which LD should be adjusted to be 350, and calculated effects size estimates for all of the 10 fraction of non-zero effects pre-specified parameters, which included LDpred-inf, 1, 0.3, 0.1, 0.03, 0.01, 0.003, 0.001, 0.0003, and 0.0001. The optimal parameter was chosen by predicting into the HRS data set and choosing the parameter that had the highest prediction *R*^2^ when the predicted phenotype was regressed on the true phenotype. This optimal parameter was then used for prediction into the ESTB. For RSS, analyses were performed for each chromosome with the chromosome-wise shrunk LD matrices from the simulation and cross-validation analyses used. The RSS-BSLMM model was run for 2 million MCMC iterations with 1 million as burn in and a thinning rate of 1 in 100 to arrive at 10,000 posterior samples for each of the model parameters. For each chromosome, the posterior mean over posterior samples for the SNP effects and 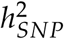 estimates was used. The chromosome-wise 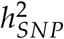 estimates were then summed to get the genome-wide estimate. To assess prediction accuracy, we calculated EGVs using the genotype data from the independent test data sets using the PLINK 2 software for all methods. Prediction *R*^2^ was calculated via linear regression of the true phenotype on that estimated from each method, which was used as a measure of prediction accuracy for each trait.

## Results

### Genome-wide simulation study

Across the simulation scenarios, we observed that BayesR or SBayesR gave the highest or equal highest mean validation prediction *R*^2^ across the 10 replicates (Figure 1). SBayesR showed the highest or equal highest mean prediction *R*^2^ of the summary statistics methodologies across all scenarios. The difference between the mean prediction *R*^2^ from BayesR and that from SBayesR was minimal for less heritable traits with SBayesR showing a marginally higher mean *R*^2^ for lower heritable traits with 50k causal variants. Prediction *R*^2^ for BayesR was maximally greater than SBayesR when 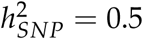 and for the 10k causal variant scenario with a relative increase of 13.2% (from 0.356 to 0.403). P+T performed well across scenarios and showed increased mean prediction *R*^2^ relative to LDpred-inf and SBLUP in the 10k causal variant scenarios but did not perform substantially better than LDpred tuned for the polygenicity parameter across all scenarios. RSS showed the closest mean prediction *R*^2^ to SBayesR in the 10k causal variant simulation scenarios. Similarly, SBLUP showed a mean prediction *R*^2^ close to SBayesR in the 50k causal variant simulation scenarios. SBayesR showed the largest nominally significant (*p*-value=0.015) improvement in prediction *R*^2^ over other summary statistics methodologies in the 10k causal variant scenario and 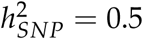 with an relative difference in mean of 3.5% (from 0.344 to 0.356) over RSS.

**Figure 1.**
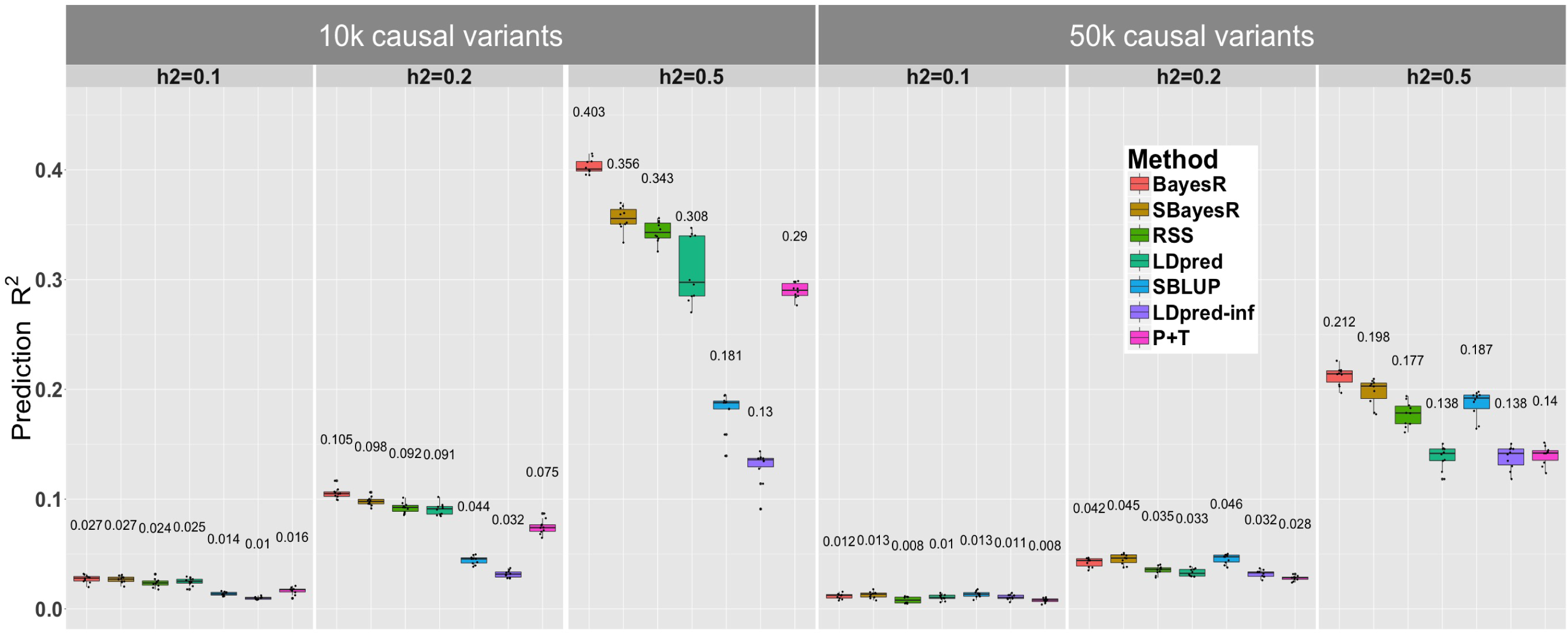
Prediction accuracy performance for the UKB genome-wide simulation. Each panel displays boxplot summaries of the prediction *R*^2^ (y-axis) in the 10,000 individual validation data set for each method (x-axis) across the 10 replicates. The simulation study contained six scenarios that varied in the number of causal variants, 10,000 (10k) and 50,000 (50k), and the true simulated heritability 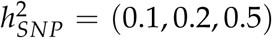. The two genetic architecture scenarios generated were: 10,000 causal variants sampled under the SBayesR model i.e., 2500, 5000, and 2500 variants from each of *N*(0, 0.01), *N*(0, 0.1), and *N*(0, 1) distributions respectively, and 50,000 causal variants sampled from a standard normal distribution. For each replicate a new sample of causal variants was chosen at random from the set of 1,094,841 HapMap 3 variants. In each panel LDpred has two boxplot summaries, one that has been optimised for the polygenicity parameter and the other is LDpred-inf, which is displayed for comparison with SBLUP. The mean prediction accuracy across the 10 replicates is displayed above the boxplot for each method.

Across all simulation scenarios, all methods except RSS showed minimal bias in 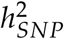 estimation (Figure S7), with HEreg showing the least bias across all scenarios. SBayesR maintained a small upward bias across all simulation scenarios and a maximum upward relative on mean bias of 5.0% (0.105 compared to 0.1) in the 10k causal variant scenarios (Figure S7). Similar to RSS, LDSC maintained a small downward bias in mean 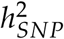 with a maximum of relative deviation of 6.4% (0.468 compared to 0.5) for the 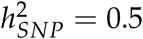 and 10k causal variant scenario.

We compared the CPU time and memory usage between all methods in each scenario. P+T, HEreg and LDSC were not compared as they required minimal relative computational resources but do not estimate the genetic effects. For the Bayesian methodologies, runtime is dependent on the length of the MCMC chain. The chain length of 4,000 MCMC iterations for BayesR was chosen as a compromise between maximum prediction accuracy and computational efficiency. We observed that a marginal relative gain in the mean prediction accuracy of 0.5% (e.g., 0.403 to 0.405) could be achieved if the chain was run for 10,000 iterations (mean runtime of 110 hours) (Figure S8) at a cost of twice the runtime. An MCMC chain length of 4,000 iterations was chosen for SBayesR to allow direct comparison with the results from BayesR with no improvement in mean prediction *R*^2^ if a chain length of 100,000 (mean runtime of 15 hours) was used (Figure S9). We observed substantial differences between prediction accuracy results from RSS when the chain length was reduced to 200,000 iterations (in an attempt to reduce computational time) (Figure S10) and we thus maintained an MCMC chain length of 2 million iterations, which was used in Zhu and Stephens^42^. Across the simulation scenarios, SBayesR had the shortest mean runtime (approximately one hour) with a greater than 10-fold improvement over the second quickest LDpred (Figure S12). SBayesR required *≈* 50 GB of memory usage, which was similar to SBLUP (35-40 GB), although SBLUP had a much longer on mean runtime. SBayesR required half the memory of the individual data BayesR, which has been highly optimised for time and memory efficiency, and showed a seven-fold improvement over LDpred and a 30-fold improvement over RSS (Figure S13). We note that the memory requirements for SBayesR are fixed for this set of variants for an arbitrary number of individuals, which is not the case for the individual level BayesR method. The total time and memory used to compute the SBayesR LD reference is not included in these assessments. The building of the sparse LD reference for SBayesR took in total 13 and 1/3 CPU days and approximately 500 GB of memory. SBayesR can compute the sparse LD matrix in parallel via dividing each chromosome into genomic ‘chunks’. We used 100 CPUs to compute the LD matrix, which brought the average runtime and memory for computing each LD matrix chunk to 3.25 hours and 5 gigabytes. These chromosome-wise LD matrices are a once off computation cost that can be distributed with the program and were used for all SBayesR and RSS analysis in the genome-wide simulation and further analyses using this HM3 variant set.

### Application to 10 quantitative traits in the UK Biobank

We compared all methods in terms of prediction accuracy and 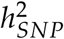 estimation across 10 quantitative traits in the UKB using five-fold cross-validation. SBayesR consistently improved or equalled the mean prediction *R*^2^ of all other methods, including the individual level BayesR method, across the five folds for 8/10 traits (Figure 2). BayesR was the only method to exceed SBayesR in mean prediction *R*^2^ and showed a relative increase of 4.3% (from 0.187 to 0.195) for heel BMD and 4.3% (from 0.349 to 0.364) for height. Heel BMD, height and FVC showed nominal significance (*p*-value = (0.007, 0.029, 0.011) respectively) in prediction accuracy improvement over RSS with a relative improvement in mean prediction *R*^2^ of 2.5% (from 0.182 to 0.187), 2.0% (from 0.342 to 0.349) and 2.5% (from 0.123 to 0.127) respectively (Figure 2). SBayesR showed larger improvements relative to LDpred tuned for the polygenicity parameter with SBayesR showing mean relative prediction *R*^2^ increases over LDpred ranging from 2% (BFP) to 37% (hBMD).

**Figure 2.**
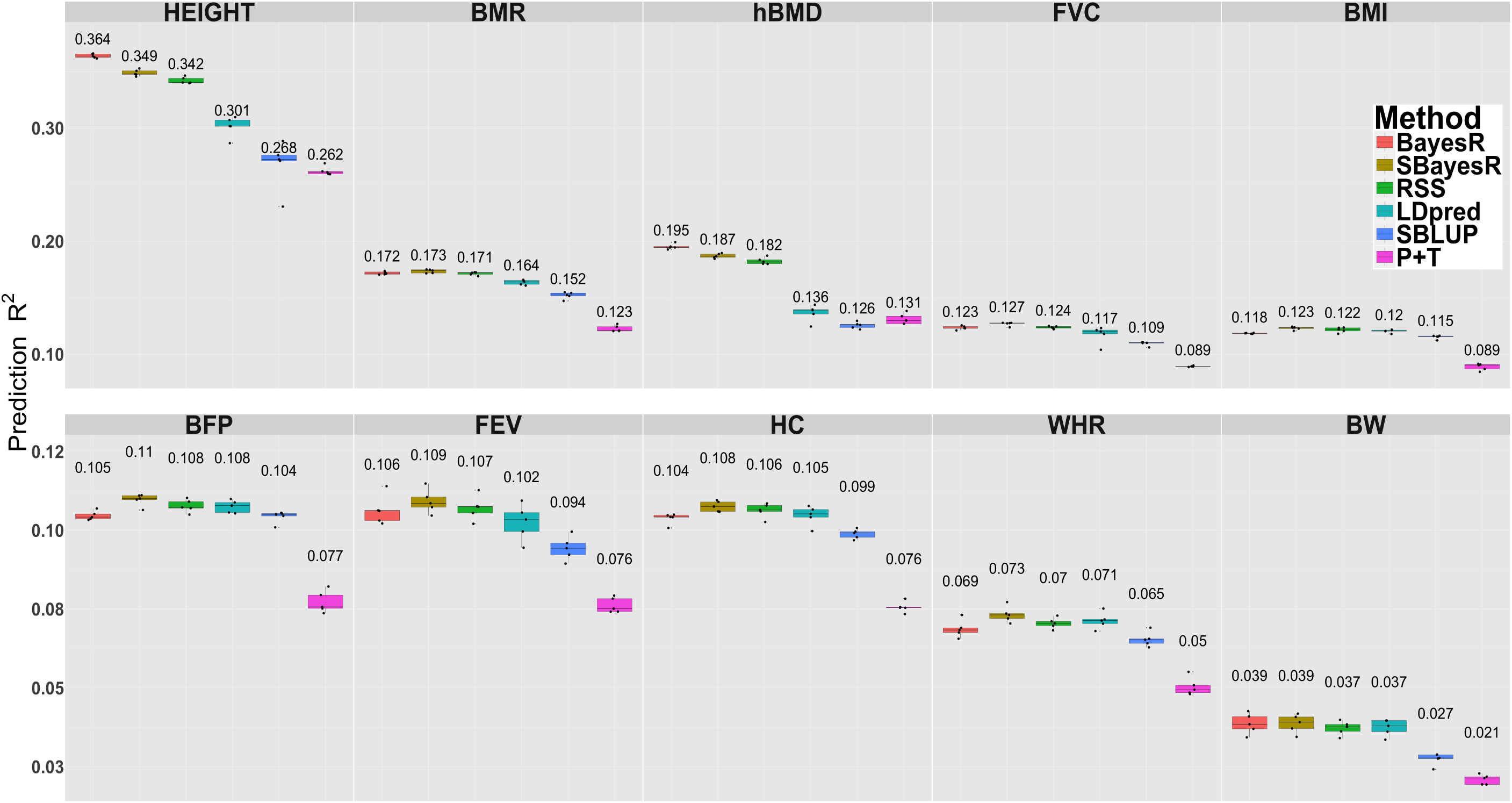
Prediction accuracy in five-fold cross-validation for 10 quantitative traits in the UK Biobank. Panel headings describe the abbreviation for 10 quantitative traits including: standing height (HEIGHT, *n*=347,106), basal metabolic rate (BMR, *n*=341,819), heel bone mineral density T-score (hBMD, *n*=197,789), forced vital capacity (FVC, *n*=317,502), body mass index (BMI, *n*=346,738), body fat percentage (BFP, *n*=341,633), forced expiratory volume in one-second (FEV, *n*=317,502), hip circumference (HC, *n*=347,231), waist-to-hip ratio (WHR, *n*=347,198) and birth weight (BW, *n*=197,778). Each panel shows a boxplot summary of the prediction *R*^2^ across the five folds with the mean across the five folds displayed above each method’s boxplot. Traits are ordered by mean estimated 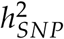 (see Figure S14) from highest to lowest.

For all traits except height, 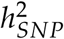 estimates were consistent across all methods (Figure S14). Across all traits except BW and FEV, SBayesR gave the highest mean 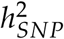 estimate and LDSC the lowest mean value, with the largest deviation in mean LDSC estimates from other methods for hBMD and height. On mean across the five folds, relative deviations in mean 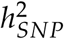 estimates between SBayesR and HEreg were between 1.0%-14.6% with the largest deviations being for WHR (6.4%), BFP (9.7%) and BW (14.7%). Similar ranges in relative deviations from mean HEreg 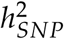 estimates were observed for other methods, with BayesR showing a range of 1.8%-20.1% and RSS 1.2%-23.1%.

We summarised the time and memory requirements of BayesR, SBayesR, RSS, LDpred and SBLUP for all traits across the five folds. P+T, HEreg and LDSC are very time and memory efficient and we therefore did not summarise their resource requirements. SBayesR on mean took approximately one to two hours and required 50 GB of memory to complete a genome wide analysis (1,094,841 HM3 variants) with variability depending on the number of non-zero variants in the model (Figures 3 and S17). For example, BFP and BMI had approximately 120,000 non zero effects whereas hBMD had approximately 30,000 and consequently the shortest runtime (Figure 3). The difference in the number of non-zero effects in the model for these traits may be driven in part by the sample size differences between BMI (*n*=346,738) and hBMD (*n*=197,789). RSS had the longest runtime with a total on mean CPU runtime being in the order of 400 hours. Again, shortening of the chain to 200,000 iterations to reduce runtime decreased the prediction accuracy of RSS with marginal changes in mean 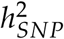 estimates (Figures S15 and S16). LDpred was the closest to SBayesR in terms of runtime with total time being 25 hours on mean across the traits. SBayesR showed a six-fold memory improvement over BayesR and LDpred and a 30 fold improvement over RSS (Figure S17). The improvements in memory between SBayesR, LDpred and SBLUP are likely a result of not having to compute the LD correlations for each fold in each trait. The memory improvement over RSS is due to the sparse matrix storage and computation in SBayesR.

**Figure 3.**
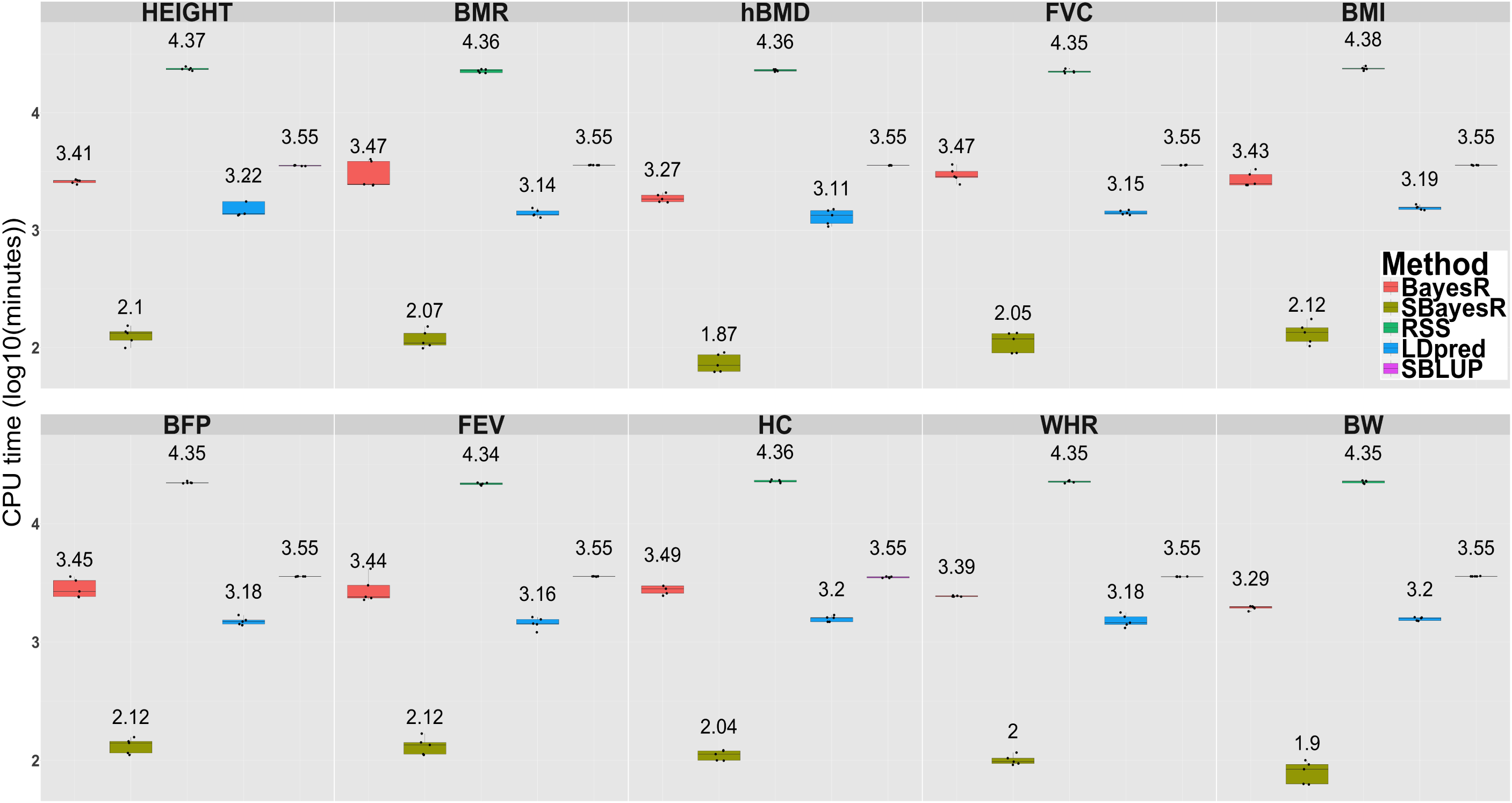
Runtime (log_10_(minutes)) comparison for BayesR, SBayesR, RSS, LDpred and SBLUP in cross-validation analysis of 10 quantitative traits in the UKB. Panel headings describe the abbreviation for 10 quantitative traits including: standing height (HEIGHT, *n*=347,106), basal metabolic rate (BMR, *n*=341,819), heel bone mineral density T-score (hBMD, *n*=197,789), forced vital capacity (FVC, *n*=317,502), body mass index (BMI, *n*=346,738), body fat percentage (BFP, *n*=341,633), forced expiratory volume in one-second (FEV, *n*=317,502), hip circumference (HC, *n*=347,231), waist-to-hip ratio (WHR, *n*=347,198) and birth weight (BW, *n*=197,778). Each panel shows a boxplot summary of runtime with the mean across the five folds displayed above each method’s boxplot. Results for RSS, LDpred and SBLUP represent the sum over time for each chromosome-wise analysis. Results for RSS and SBayesR do not include the time to compute the LD reference matrix. Results for P+T, HEreg and LDSC are not shown as they required relatively minimal computing resources.

### Across biobank prediction analysis

Overall, SBayesR gave similar but consistently higher prediction *R*^2^ values than BayesR for both BMI and height in both the HRS and ESTB samples (Figure 4), when the summary statistics from the full European ancestry (related and unrelated individuals) UKB data set were used (*n* = 453, 458 and *n* = 454, 047 for BMI and height respectively). When the summary statistics from Yengo *et al.*^49^ were used, a further improvement in prediction *R*^2^ was observed for SBayesR and RSS, except for height and in HRS (Figure 4). SBayesR and RSS gave the same prediction *R*^2^ values for BMI with marginal increases of SBayesR over RSS for height, which is consistent with the results from the cross-validation. The maximum increase in SBayesR prediction *R*^2^ relative to the BayesR analysis using just the UKB data for BMI was 11.3% (from 0.106 to 0.118) and 4.9% (from 0.307 to 0.322) for height in the ESTB sample when the summary statistics from the^49^ data set were used. The maximum increase in prediction *R*^2^ relative to that from the predictor built from the GCTA-COJO analysis thresholded at *p*-value< 0.001 performed in Yengo *et al.*^49^ for BMI was 32.5% (from 0.089 to 0.118) in the ESTB. For height, we observed a maximum relative increase of 31.6% (from 0.244 to 0.321) in prediction *R*^2^ over the P+T predictor of Yengo *et al.*^49^ in the HRS sample when the summary statistics from the full UKB data set were used for SBayesR analysis.

**Figure 4.**
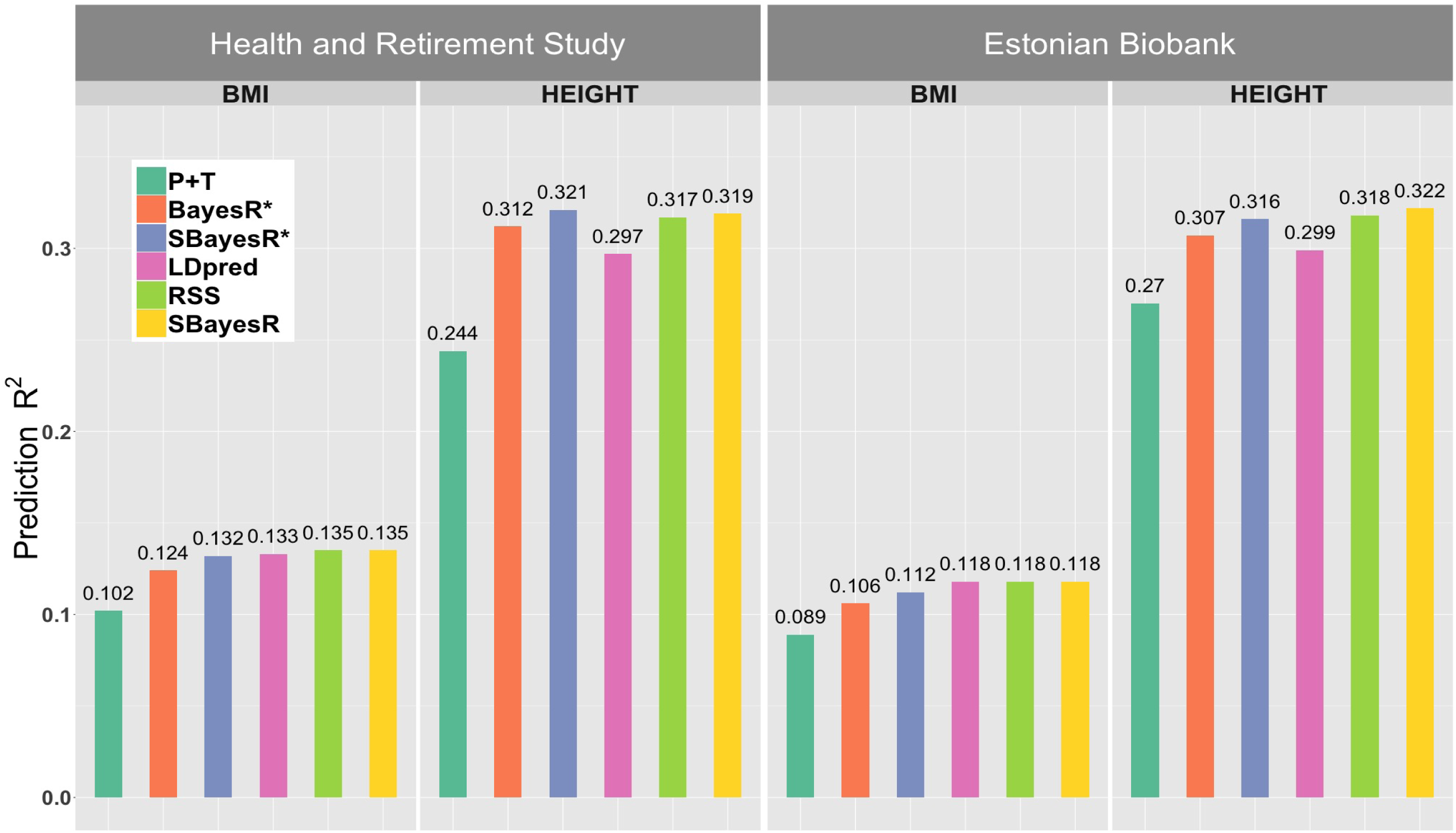
Prediction accuracy for height and body mass index in the independent Health and Retirement Study and Estonian Biobank data sets. Panels depict prediction *R*^2^ (y-axis) generated from regression of the predicted phenotype on the observed phenotype for body mass index (BMI) and height for different methods in the independent HRS and ESTB data sets. P+T refers refers to the prediction *R*^2^ generated from the summary statistics of Yengo et al. 2018 (*n ≈* 700,000), which included 6,781 SNPs for BMI and 11,816 SNPs for height from a GCTA-COJO analysis thresholded at *p*-value < 0.001. The BayesR* and SBayesR* predictions were calculated using 1,094,841 HM3 variants estimated from the full set of unrelated and related UKB European individuals (*n* = 453, 458 and *n* = 454, 047 for BMI and height respectively). Summary statistics for SBayesR analysis for the UKB European individuals were generated using the BOLT-LMM software. All other prediction *R*^2^ results were generated using summary statistics methodology and were calculated from the analysis of summary statistics from Yengo *et al.* ^49^ for 909,293 and 932,969 variants for BMI and height that overlapped with the 1,094,841 HM3 variants set used for the UKB analyses. The overlap of the sets of variants used in each of the analyses and those available in the imputed HRS and ESTB data sets for prediction had a minimum value of 98%.

## Discussion

Clinically relevant genetic predictors for complex traits and disorders will require the analysis of data from large consortia and biobank initiatives, with sample sizes for GWASs set to soon regularly reach into the millions of individuals. Efficient methods that produce theoretically optimal predictors under the multiple regression model will therefore be critical to this goal. We have presented one solution, that rests on an extension of the established summary statistics methodological framework to include a class of point-normal mixture prior Bayesian regression models, which encompasses many previously proposed models^27,28,31^.

We observed that the cohort used to construct the LD reference matrix influenced the prediction accuracy and 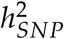 estimation. The LD reference built from a random sample of 50k individuals from the UKB showed the maximum prediction accuracy and smallest upward bias in 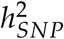 estimation across all scenarios in the small-scale simulation on two chromosomes although these were marginal relative to those from the smaller UK10K sequence reference. We anticipate that the UKB will contribute to future large-scale GWASs and thus we anticipate that the LD reference built from a large subset of this cohort in this study will be highly beneficial to future summary statistics analyses of complex traits.

The simulation studies thoroughly compared prediction methods as a function of genetic architecture, LD reference and other parameters, with SBayesR generally outperforming other methods. In simulation, P+T performed well across scenarios and showed increased mean prediction *R*^2^ relative to SBLUP and LDpred-inf in a subset of the simulation scenarios but did not perform better than LDpred tuned for the polygenicity parameter across all scenarios, which is contrary to observations made by Mak *et al.*^46^. In the five-fold cross-validation, SBayesR consistently improved or equalled the mean prediction *R*^2^ of all other methods, with a marginal improvement over the individual level BayesR method for most traits. SBayesR maintained a minimal upward bias across all simulation scenarios (maximum upward bias of *≈* 5.0%) and showed 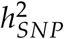 estimates close to that from HEreg in the cross-validation analysis. SBayesR gave consistently higher but similar prediction *R*^2^ values than BayesR for both BMI and height in across biobank predictions into the HRS and ESTB samples. This was both the case when the summary statistics from the full European UKB data set were used with a further improvement in prediction *R*^2^ observed when the summary statistics from Yengo *et al.*^49^ were used. The maximum increase in prediction *R*^2^ relative to the prediction *R*^2^ from Yengo *et al.*^49^ for height was in the the HRS sample when the summary statistics from the full UKB data set were used 31.6% (from 0.244 to 0.321). The maximal prediction accuracy in HRS and ESTB was *R*^2^ = 0.321 (correlation between outcome and predictor of 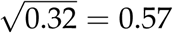), which is starting to reach the initial estimates of 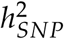 of 0.45 in Yang *et al.*^21^.

The observation that SBayesR improves on the BayesR prediction accuracy in real data cross-validation and independent out-of-sample prediction is contrary to expectation. In the small-scale simulation we observed that SBayesR using the full LD correlation matrix and BayesR, which are theoretically equivalent, returned equal on mean prediction accuracies and 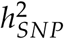 estimates and thus the numerical implementation is not substantially superior. When we scaled the simulation to the whole genome, we observed that BayesR showed relatively smaller improvements over SBayesR for lower heritable traits in the 10k causal variant scenarios and for 50k causal variants scenarios SBayesR improved on BayesR mean prediction *R*^2^ for lower heritability traits, which was also the case for lower heritable traits in the cross-validation. For lower heritable traits the length of the BayesR MCMC chain may play a larger role with marginal improvements in prediction accuracy observed for longer BayesR chains for the 10k causal variants and 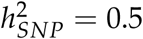 genome-wide simulation scenario. A further factor is the impact of using summary statistics results from a LMM (e.g., Loh *et al.*^25^), where the model is derived under the assumption that the summary statistics have been generated from a least squares analysis. The use of summary statistics from a LMM will affect the reconstruction of **X**′**y**. One further, and likely major, difference between these two methods is the ignorance of interchromosomal LD in the SBayesR method, where interchromosomal LD may result from genetic sampling in finite population sizes, population structure and non-random mating (e.g., assortative mating). The incorporation of this information appears only advantageous for predictions performed within an independent subset from the same population e.g., the partitioning of the UKB in the simulation studies and in cross-validation. The HRS and ESTB data are unlikely to contain the same interchromosomal LD correlation structure and thus its inclusion in the BayesR analysis may be partially detrimental as it comes into the model as informative within data set (UKB) but as noise across data sets (UKB to HRS/ESTB). One hypothesis for this is that the HRS and ESTB populations have different patterns of assortative mating for specific traits than in the UKB, or individuals in HRS or ESTB are more randomly mated than in those in the UKB.

The method is implemented in a very efficient and user-friendly software tool that maximises computational efficiency via precomputing and efficiently storing sparse LD matrices that account for the variation in the number of LD ‘friends’ for each variant. In simulation and cross-validation we showed large fold improvements in time and memory over current state-of-the-art individual and summary data methods. The improvements in efficiency are not just a result of the computational implementation but are a contributed to by the faster convergence of the the Gibbs sampling algorithm. This is evidenced by the comparison with RSS, which requires a much longer chain length to arrive at maximum prediction accuracy. Importantly, once the GWAS effect size estimates have been generated the method’s runtime is independent of the sample size making it applicable to an arbitrary number of individuals.

We found that model convergence is sensitive to inconsistencies in summary statistics generated from external consortia and meta-analyses. We observed that the shrinkage estimator of the LD matrix^36^ can assist with more stable model convergence. We observed a persistent small upward bias in 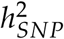 estimation, which was also observed by Zhu and Stephens^42^. We did not observe this upward bias in the RSS analyses, which may in part be attributed to the much larger LD reference used. Zhu and Stephens^42^ hypothesised that the persistent upward inflation to be due to deviations from the assumption of small effects underlying the RSS model. However, we did not observe large differences in upward bias in 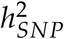 estimation between simulation scenarios containing very large effects compared to scenarios with effect sizes similar to those for very polygenic traits. It is difficult to assess the impact of the small effect assumption versus the contribution from the replacement of the **D** and LD matrices with estimates reconstructed from GWAS summary statistics from external references or a subset of the GWAS data. Through simulation, we observed that this upward bias can be minimised through an optimally sparse and sufficiently large LD reference. The impact from residual population stratification in the GWAS summary statistics is another potential source in upward bias in 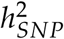 estimates but was not investigated via simulation.

There are distinct practical advantages in estimating 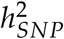 and the genetic effects within one framework with the method encompassing many available summary statistics methodologies. Zhu and Stephens^42^ presented a similar omnibus method and showed the capacity of this similar methodology for variant mapping. Although we haven’t assessed our method’s effectiveness for mapping causal variants we expect it to be capable of performing this task, which is to be inherited from the individual-level BayesR method’s capacity to perform this task^31,63,64^. SBayesR estimates all parameters from the data and does not require any post-hoc tuning of prediction relevant parameters in a test data subset (as in the polygenicity parameter in LDpred or P+T), which has practical advantages in terms of relieving the analytical burden of tuning these parameters in an external data set. Furthermore, this leads to more generalisable predictors as the parameters have been optimised over all possible values rather than selected from a finite grid.

The method assumes certain ideal data constraints such as summary data computed from a single set of individuals at fully observed genotypes as well as minimal imputation error and data processing errors such as allele coding and frequency mismatch. Summary data in the public domain often substantially deviate from these ideals and can contain residual population stratification, which is not accounted for in this model. Practical solutions to these ideal data deviations include the use of data that are imputed and the restriction of analyses to variants that are known to be imputed with high accuracy as in Bulik-Sullivan *et al.*^39^ and Zhu and Stephens^42^. We found that the simple filtering of SNPs with sample sizes that deviate substantially from the mean across all variants from an analysis, as in Pickrell *et al.*^62^, when using summary statistics from the public domain substantially improved model convergence. We explored LD pruning of variants to remove variants in very high LD (*R*^2^ > 0.99) but found that this did not substantially improve model convergence or parameter estimates although this was not formally assessed. However, removal of high LD regions, such as the MHC region improved model convergence for real traits. High LD regions are expected to have the potential to be extreme sources of model misspecification with the model expecting summary data in to be very similar for variants in high LD. Small deviations due to data error not expected in the model likelihood at these loci thus have high potential to lead to model divergence (see Zhu and Stephens^42^ for further discussion). Future research into efficient diagnostic tools and methods that can assist analysts with the assessment of sources of bias and error and summary data quality would be highly beneficial.

We expect that as GWAS sample sizes continue to grow that polygenic predictions will become more accurate. We expect that they will be important in future clinical settings, for improving prediction in diverse populations and for understanding quantitative genetics more generally. The very efficient implementation of our method makes the analysis of millions of variants and an arbitrary number of individuals possible. The implementation and model are very flexible and can easily incorporate other model formalisations such as understanding the contributions of genomic annotations to prediction and 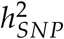 enrichment such as in^41,65^ or understanding genetic architecture via summary statistics versions of models such as those presented in Gazel *et al.*^66^ and Zeng *et al.*^30^.

## Acknowledgements

The authors would like to thank the members of the Program in Complex Genetics for their insights and helpful discussion. We are grateful to Xiang Zhu for assistance and discussion concerning the Residual with Summary Statistics methodology. The authors would like to acknowledge the support from the Australian Research Council (DP160102400), the Australian National Health and Medical Research Council (1113400, 1078037, 1078901 and 1080157), the National Institute of Health (R21 ES025052 and R01 MH100141) and the Sylvia & Charles Viertel Charitable Foundation. We gratefully acknowledge CQU’s eResearch support and the use of the High Performance Computing facility (www.cqu.edu.au/hpc) in developing the updated BayesR software. **UKB**: This study has been conducted using UK Biobank resource under Application Number 12514. UK Biobank was established by the Wellcome Trust medical charity, Medical Research Council, Department of Health, Scottish Government and the Northwest Regional Development Agency. It has also had funding from the Welsh Assembly Government, British Heart Foundation and Diabetes UK. **ARIC**: The Atherosclerosis Risk in Communities Study is carried out as a collaborative study supported by National Heart, Lung, and Blood Institute contracts (HHSN268201100005C, HHSN268201100006C, HHSN268201100007C, HHSN268201100008C, HHSN268201100009C, HHSN268201100010C, HHSN268201100011C, and HHSN268201100012C), R01HL087641, R01HL59367 and R01HL086694; National HumanGenome Research Institute contract U01HG004402; and National Institutes of Health contract HHSN268200625226C. The authors thank the staff and participants of the ARIC study for their important contributions. Infrastructure was partly supported by Grant Number UL1RR025005, a component of the National Institutes of Health and NIH Roadmap for Medical Research. **UK10K**: The UK10K project was funded by the Wellcome Trust award WT091310. Twins UK (TUK): TUK was funded by the Wellcome Trust and ENGAGE project grant agreement HEALTH-F4-2007-201413. The study also receives support from the Department of Health via the National Institute for Health Research (NIHR)-funded BioResource, Clinical Research Facility and Biomedical Research Centre based at Guy’s and St. Thomas’ NHS Foundation Trust in partnership with King’s College London. Dr Spector is an NIHR senior Investigator and ERC Senior Researcher. Funding for the project was also provided by the British Heart Foundation grant PG/12/38/29615 (Dr Jamshidi). A full list of the investigators who contributed to the UK10K sequencing is available from www.UK10K.org. **HRS**: HRS is supported by the National Institute on Aging (NIA U01AG009740). The genotyping was funded separately by the National Institute on Aging (RC2 AG036495, RC4 AG039029). Genotyping was conducted by the NIH Center for Inherited Disease Research (CIDR) at Johns Hopkins University. Genotyping quality control and final preparation of the data were performed by the Genetics Coordinating Center at the University of Washington. **ESTB**: The Estonian Genome Centre of University of Tartu Study was supported by EU Horizon 2020 grants 692145, 676550, and 654248; Estonian Research Council Grant IUT20-60, NIASC, EIT Health; NIH BMI grant 2R01DK075787-06A1; and the European Regional Development Fund (project 2014-2020.4.01.15-0012 GENTRANSMED).

## Author contributions

P.M.V., J.Y., M.E.G. and N.R.W. conceived the study. P.M.V., J.Y., L.R.L-J and J.Z. designed the experiment. J.Z., L.R.L-J and M.E.G. derived the analytical methods. L.R.L-J and J.Z. conducted all analyses with assistance from J.S. and guidance from P.M.V., J.Y., L.Y., G.M., and H.W. J.Z. and L.R.L-J developed the GCTB software. G.M. developed the updated version of the BayesR software. K.E.K, L.Y. and Z.Z. performed the initial preparation and quality control of the UK Biobank data. J.S., R.M., T.E., and A.M supplied and performed initial quality control on the Estonian Biobank data. L.R.L-J wrote the manuscript with the participation of all authors in particular P.M.V., J.Y., and J.Z. All authors reviewed and approved the final manuscript.

