## Supplemental Figures and Note for "Improved polygenic prediction by Bayesian multiple regression on summary statistics"

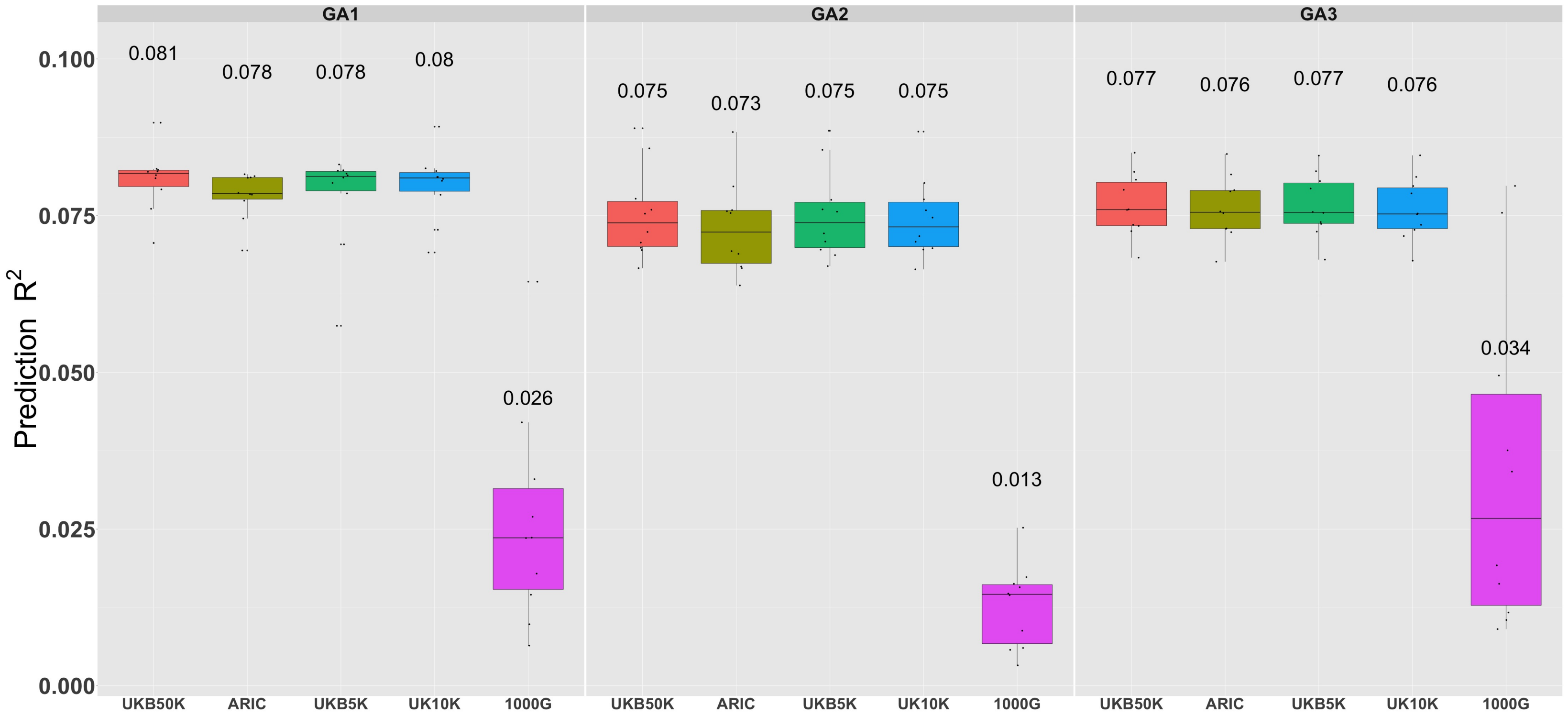


**Figure S1 Prediction accuracy $(R^2)$ for SBayesR using different LD matrix reference cohorts in the simulation on chromosomes 21 and 22.** Each panel displays boxplot summaries of the prediction $R^2$ (y-axis) from the SBayesR method in the 10,000 individual validation data set for each LD reference cohort (x-axis). Each boxplot summarises results from 10 replicates for each of the three simulation scenarios in the two chromosome simulation i.e., the first genetic architecture (GA1) contained two causal variants of large effect explaining 3% and 2% of the phenotypic variance respectively and a polygenic tail of 1,498 causal variants sampled from a N(0, 0.05/1, 498) distribution such that the expected total genetic variance explained by all variants was 0.1. The second architecture (GA2) was simulated under a BayesR model with three sets of causal variants: the first contained 1,445 causal variants sampled from a $N(0, 0.06/1445)$ distribution, the second contained 50 causal variants sampled from a $N(0, 0.02/50)$ distribution and the third 5 causal variants sampled from $N(0, 0.02/5)$ distribution. The third architecture (GA3) contained 1,500 variants sampled from a $N(0, 0.1/1500)$ distribution. The mean $R^2$ across the 10 replicates is displayed above the boxplot for each cohort. Poorer prediction accuracies for the 1000G cohort are hypothesised to be primarily driven by the small sample size ($n = 378$) of this reference.

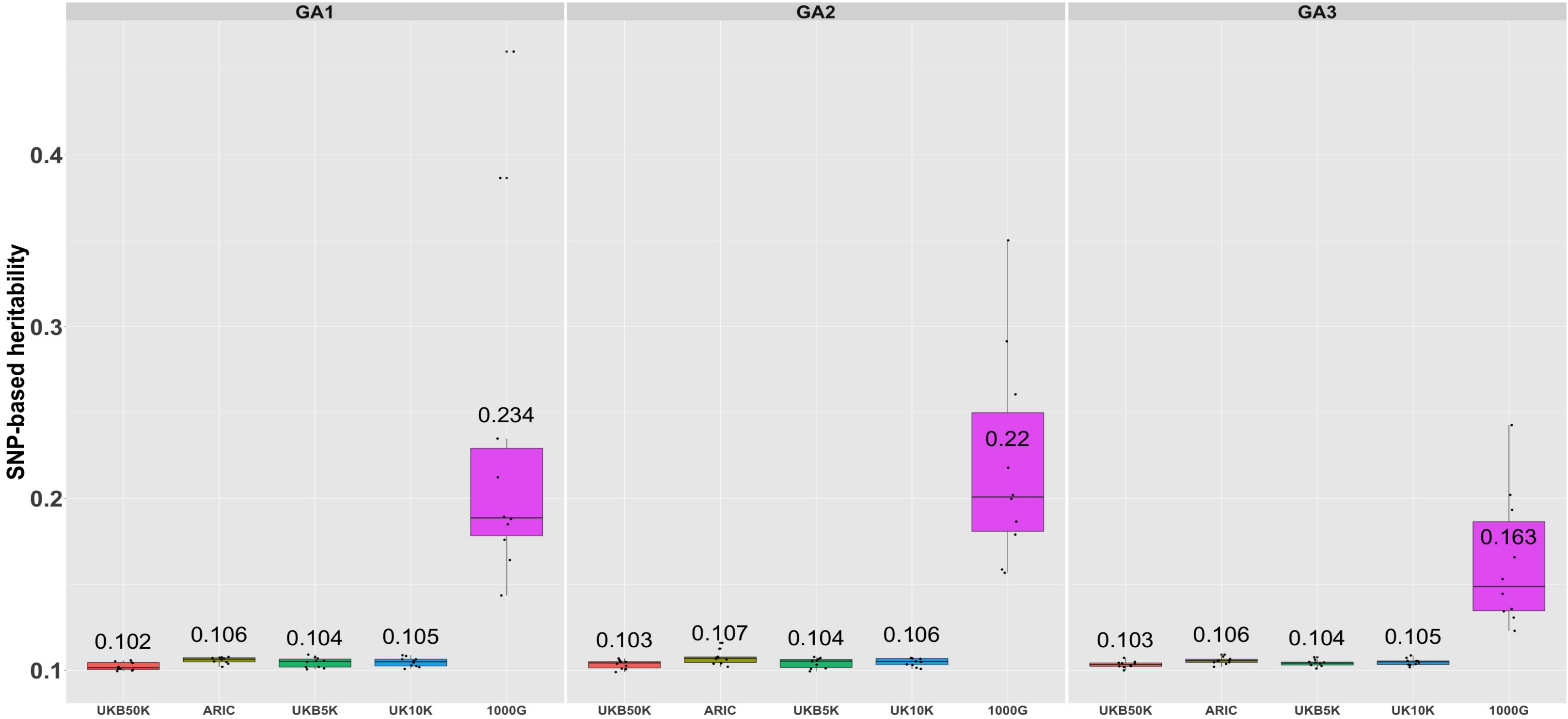


**Figure S2 SNP-based heritability estimation $(h^2_{SNP})$ for SBayesR using different LD matrix reference cohorts in the simulation on chromosomes 21 and 22.** Each panel displays boxplot summaries of $h^2_{SNP}$ estimates (y-axis) for each LD reference cohort (x-axis) across the 10 replicates for each of the three simulation scenarios in the two chromosome simulation. Each trait has a simulated true $h^2_{SNP} = 0.1$ and 1,500 causal variants. The mean $h^2_{SNP}$ across the 10 replicates is displayed above the boxplot for each cohort. Inflated $h^2_{SNP}$ estimates for the 1000G cohort are hypothesised to be primarily driven by the small sample size ($n = 378$) of this reference. See Figure S1 for descriptions of the genetic architectures (GA1, GA2, GA3).

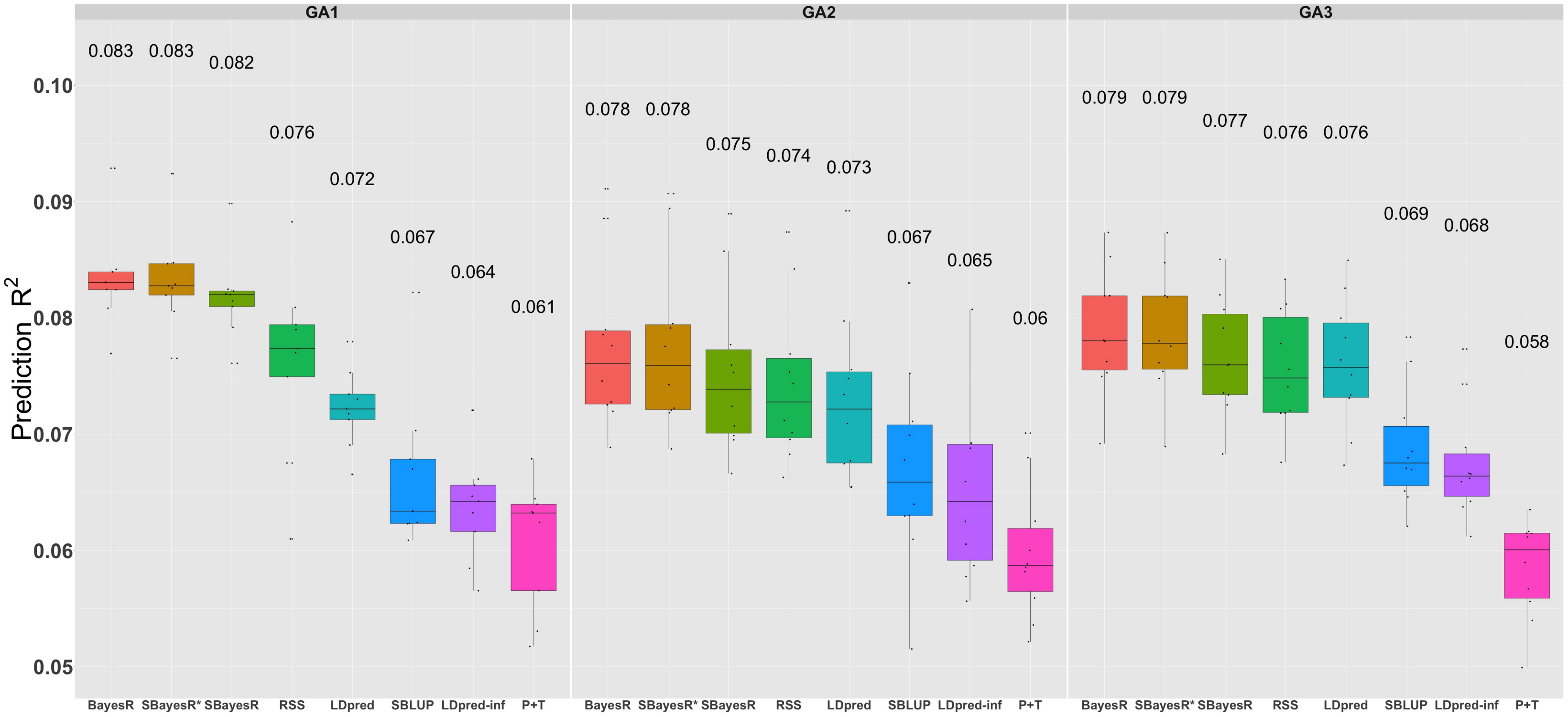


**Figure S3 Prediction accuracy performance using different methods in the simulation on chromosomes 21 and 22.** Each panel displays boxplot summaries of the prediction $R^2$ (y-axis) in the 10,000 individual validation data set and an LD reference generated from a random subset of 50,000 individuals from the UKB. Each boxplot shows the prediction $R^2$ across the 10 replicates for each of the three simulation scenarios in the two chromosome simulation i.e., the first genetic architecture (GA1) contained two causal variants of large effect explaining 3% and 2% of the phenotypic variance respectively and a polygenic tail of 1,498 causal variants sampled from a N(0, 0.05/1, 498) distribution such that the expected total genetic variance explained by all variants was 0.1. The second architecture (GA2) was simulated under a BayesR model with three sets of causal variants: the first contained 1,445 causal variants sampled from a $N(0, 0.06/1445)$ distribution, the second contained 50 causal variants sampled from a $N(0, 0.02/50)$ distribution and the third five causal variants sampled from $N(0, 0.02/5)$ distribution. The third architecture (GA3) contained 1,500 variants sampled from a $N(0, 0.1/1500)$ distribution. SBayesR* corresponds to the analysis using the SBayesR model and the full set of 100,000 individuals used in the GWAS analysis to create the LD matrix. This LD matrix includes all pairwise correlations i.e., includes inter-chromosomal LD. The mean $R^2$ across the 10 replicates is displayed above the boxplot for each cohort.

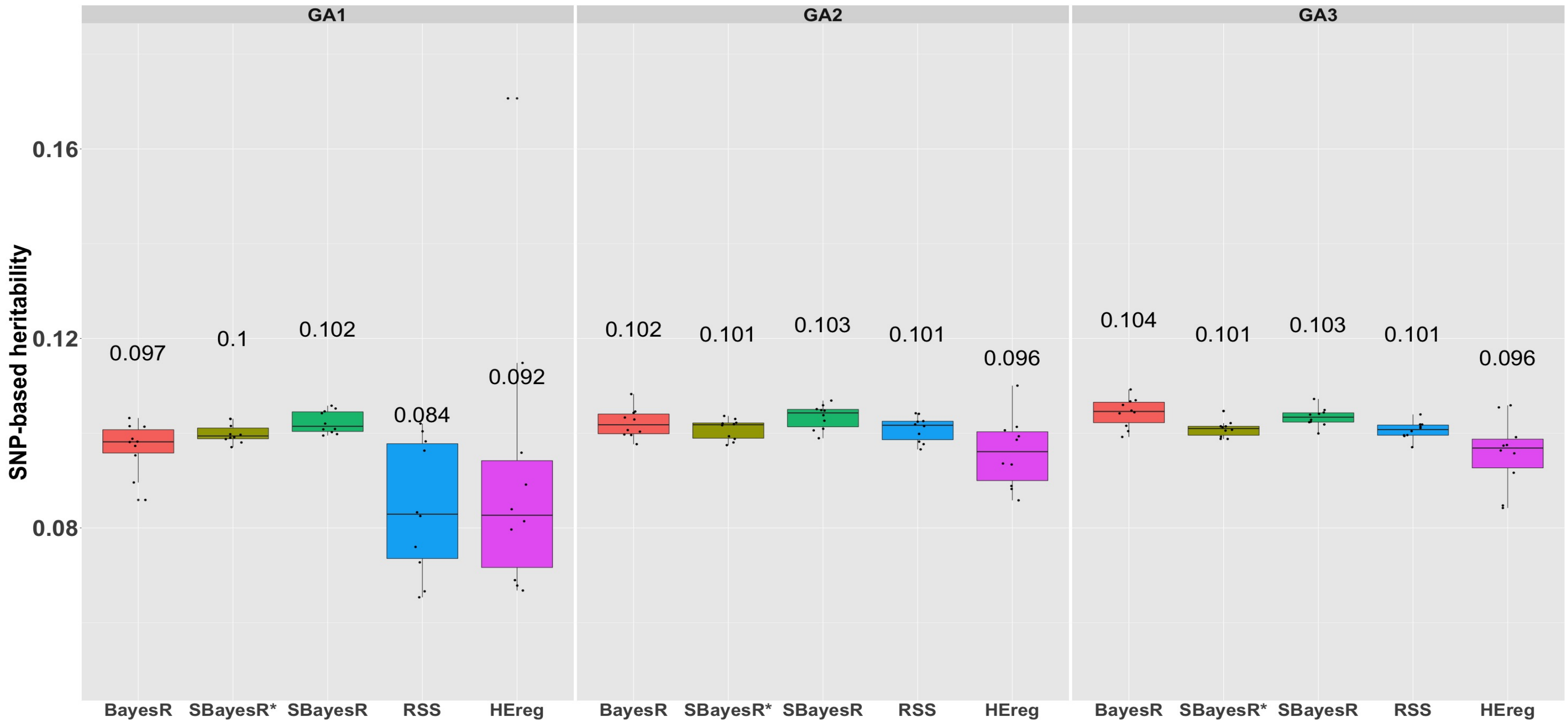


**Figure S4 SNP-based heritability ($h^2_{SNP}$) estimation for different methods in the simulation on chromosomes 21 and 22.** Each panel displays boxplot summaries of the $h^2_{SNP}$ estimates (y-axis) for each method (x-axis) across the 10 replicates in each of the three two chromosome simulation scenarios. Each trait has a simulated true $h^2_{SNP} = 0.1$ and 1,500 causal variants. SBayesR* corresponds to the analysis using the SBayesR model and the full set of 100,000 individuals used in the GWAS analysis to create the LD matrix. The mean $h^2_{SNP}$ across the 10 replicates is displayed above the boxplot for each method. See Figure S1 for descriptions of the genetic architectures (GA1, GA2, GA3)

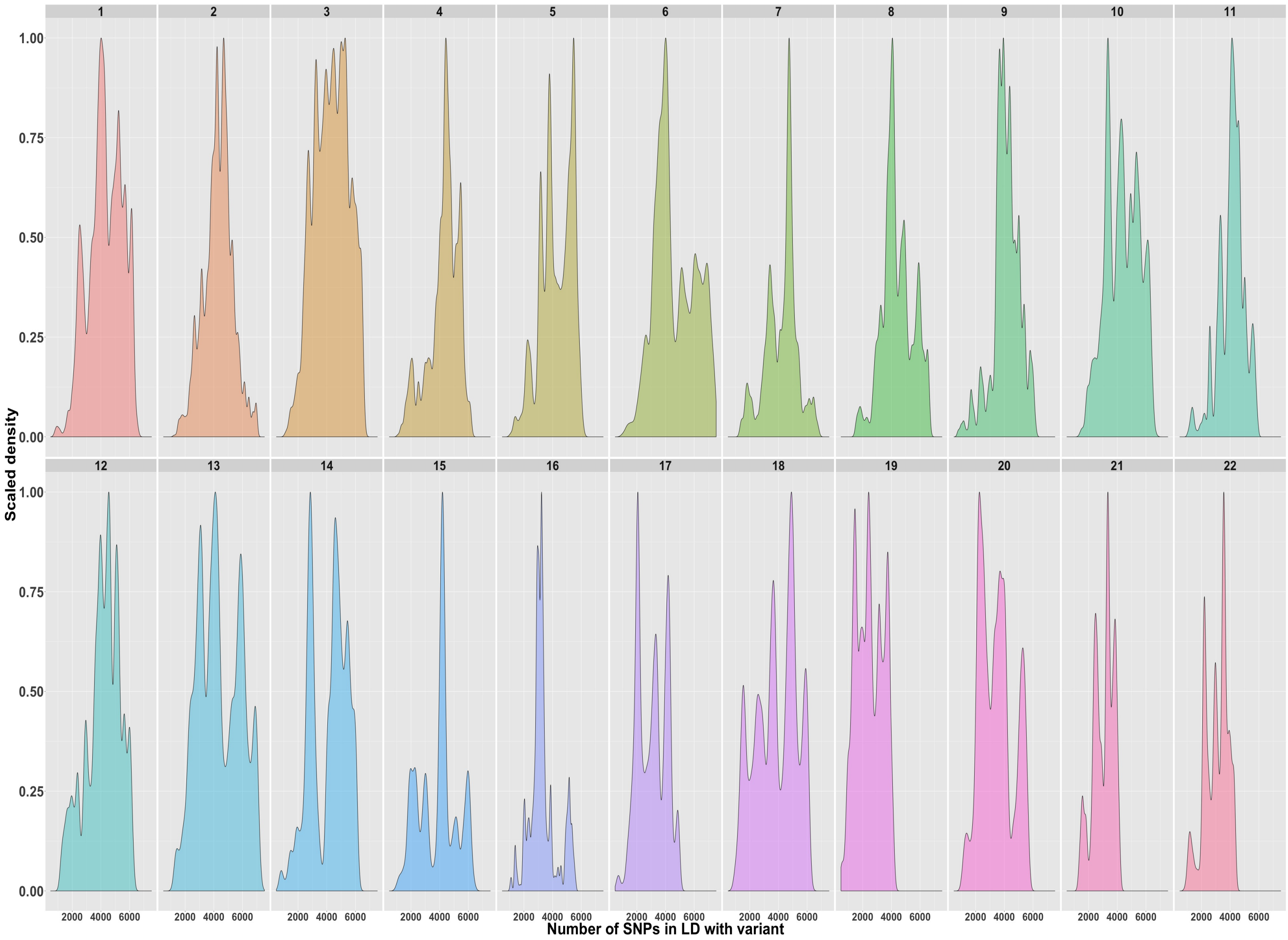


**Figure S5 Variability in the number of 'LD friends' from the shrunk-sparse LD correlation matrix within chromosome for each of 1.09 million HapMap3 variants in the UKB.** The chromosome-wise LD matrices were calculated using imputed genotype data for a random set of 50,000 individuals from the unrelated European individuals in the UKB data. The LD matrices depicted were shrunk and the number of LD friends represents the number of non-zero correlations for each variant.

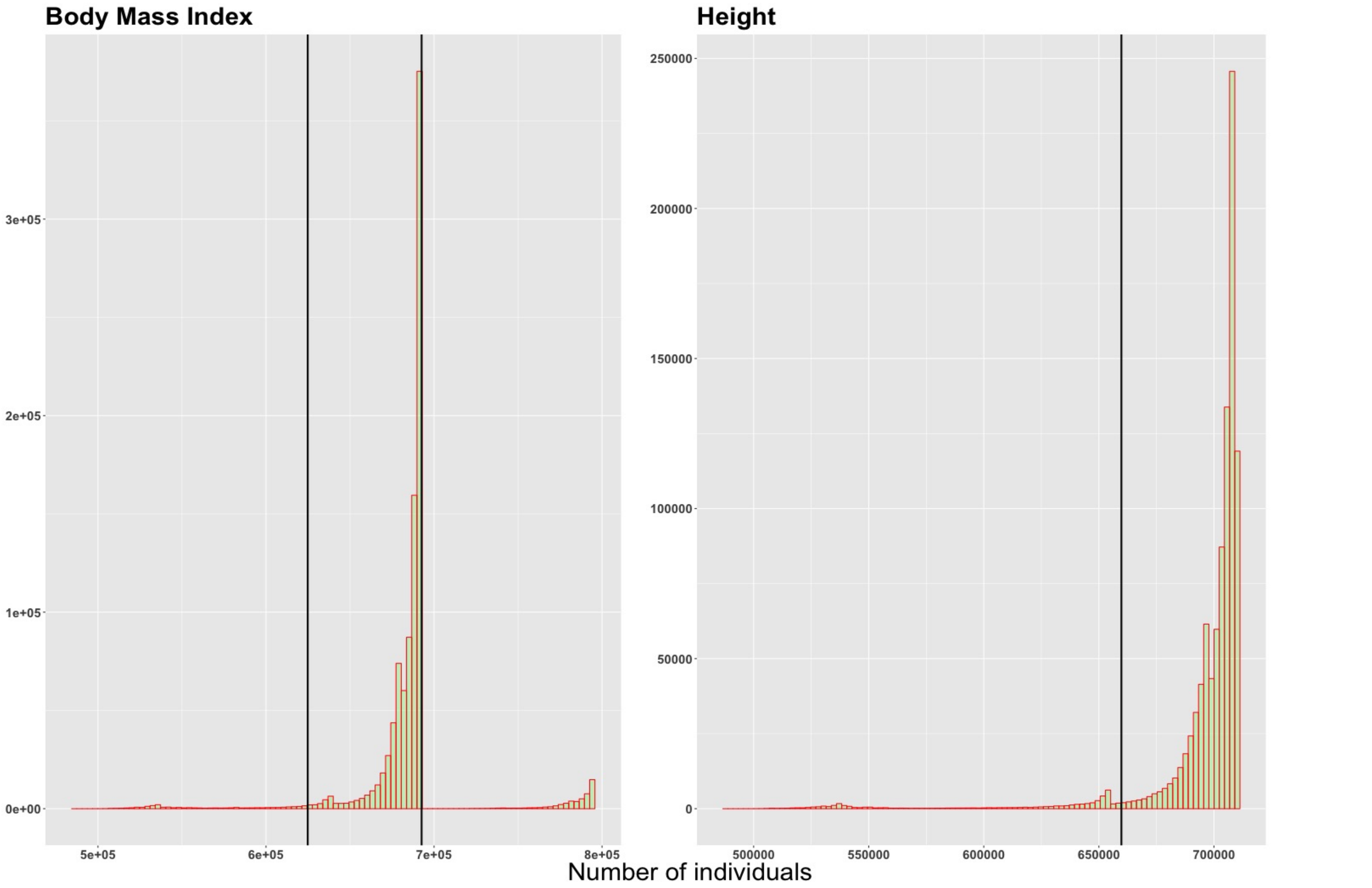


**Figure S6 Distribution and truncation of per-variant sample size from BMI and height summary statistics for 982,000 HapMap3 variants from Yengo *et al.*[1].** The 982,000 variants are those that overlap between the summary statistics made available from Yengo *et al.*[1] and the 1.09 M HM3 variants used in the simulation and cross-validation analyses. Vertical bars indicate the 0.025 and 0.95 percentiles for BMI and the 0.05 percentile for height. These truncations on $n$ reduced the variant sets to 909,293 and 932,969 for BMI and height respectively. This truncation is required for model stability as the RSS and SBayesR models assume that the summary data were generated from the same set of individuals.

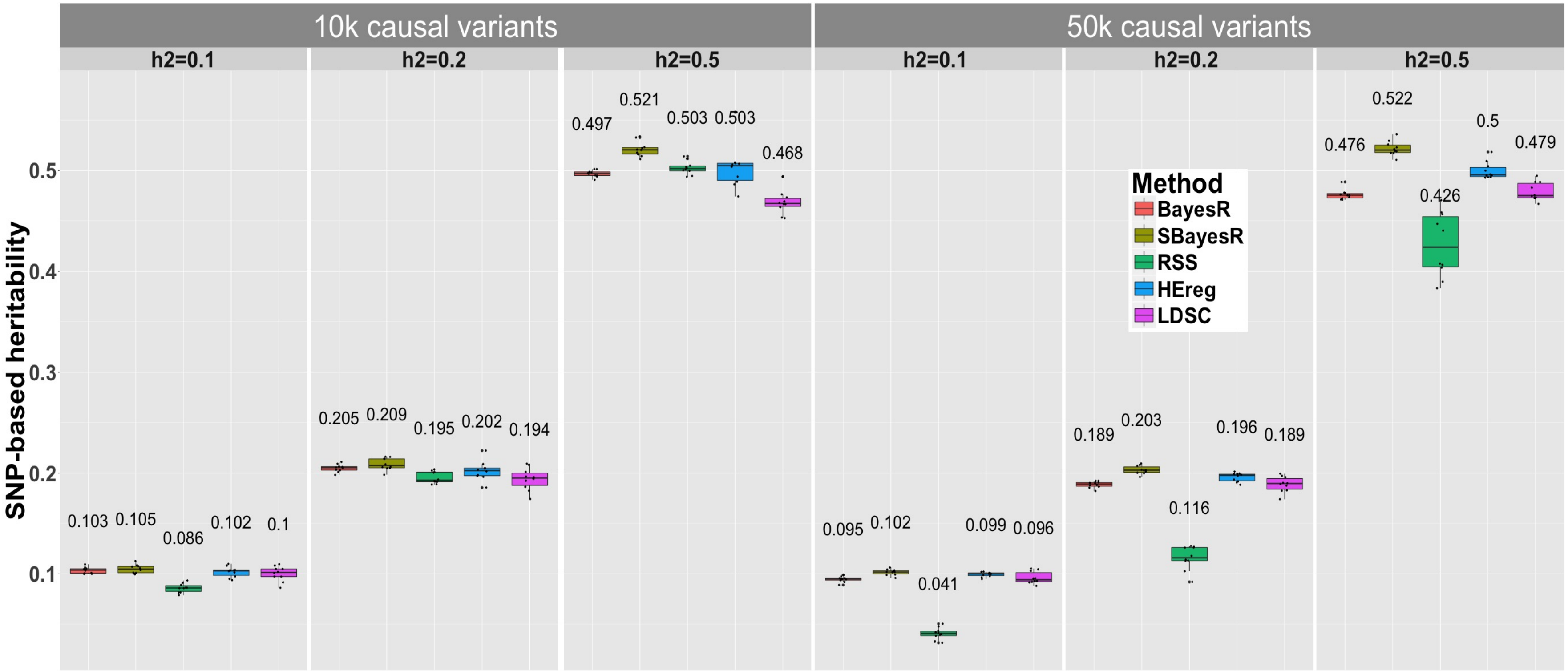


**Figure S7 SNP-based heritability ($h^2_{SNP}$) estimation performance for different methods in UKB genome-wide simulation.** Each panel displays boxplot summaries of $h^2_{SNP}$ estimates (y-axis) for each method (x-axis) across the 10 replicates for each of the six simulation scenarios that varied in the number of causal variants, 10k and 50k, and the true simulated $h^2_{SNP} = (0.1, 0.2, 0.5)$. Two genetic architecture scenarios were generated: 10,000 causal variants sampled under the SBayesR model i.e., 2500, 5000, and 2500 variants from each of $N(0, 0.01\sigma^2_\beta)$, $N(0, 0.1\sigma^2_\beta)$, and $N(0, \sigma^2_\beta)$ distributions respectively and $\sigma^2_\beta = 1$. For the second architecture, 50,000 causal variants were sampled from a standard normal distribution. For each replicate a new sample of causal variants was chosen at random from the set of 1,094,841 HapMap 3 variants. The mean $h^2_{SNP}$ estimate across the 10 replicates is displayed above the boxplot for each method.

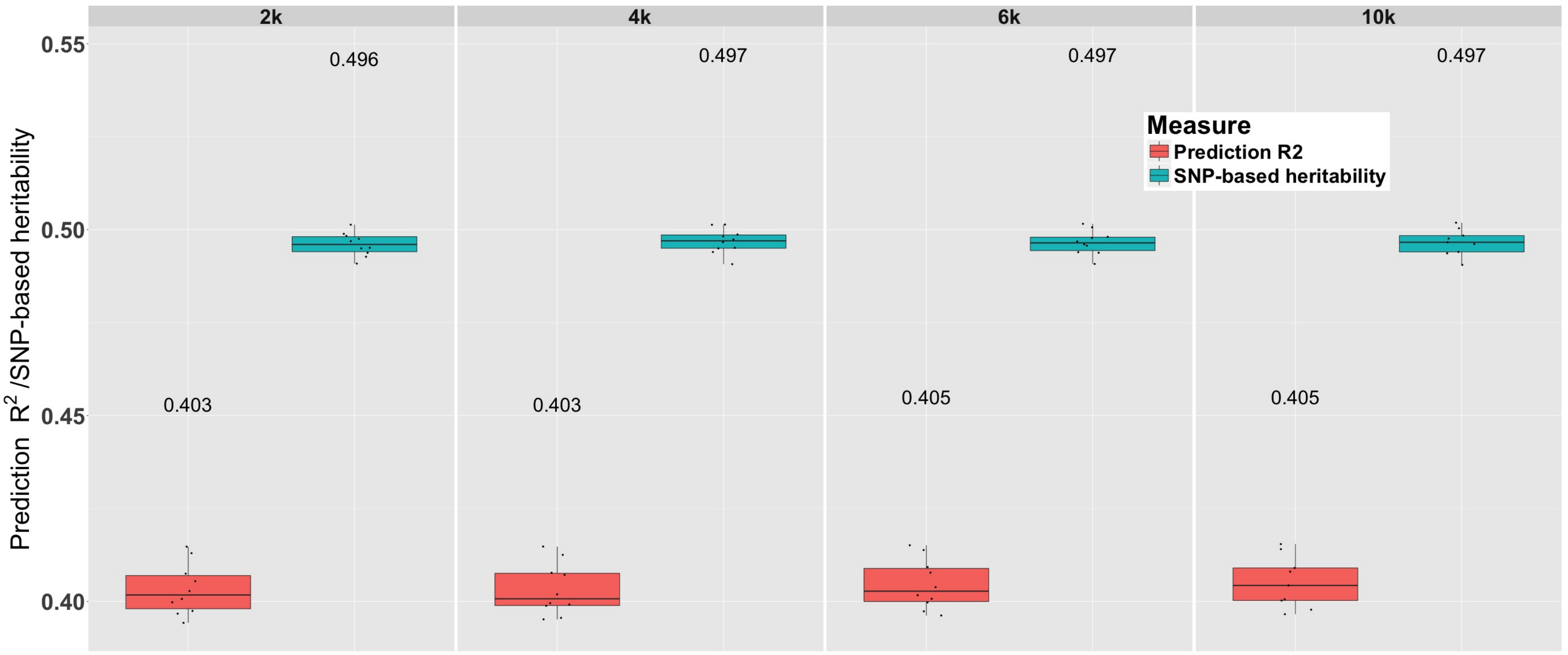


**Figure S8 BayesR prediction accuracy and SNP-based heritability ($h^2_{SNP}$) estimation change with MCMC chain length for one scenario of UKB genome-wide simulation.** Each panel displays boxplot summaries of prediction $R^2$ and $h^2_{SNP}$ estimates (y-axis) for 2,000 (2k), 4,000 (4k), 6,000 (6k) and 10,000 (10k) MCMC iterations of the BayesR method[2]. Each boxplot shows the results from the 10 replicates in the 10k (simulated under a BayesR model) causal variant and the true simulated $h^2_{SNP} = 0.5$ scenario. The mean prediction $R^2$ and $h^2_{SNP}$ estimates across the 10 replicates are displayed above the relevant boxplot. The mean run time for each of the 2k, 4k, 6k and 10k MCMC iterations scenarios was 32.5, 56.7, 77.9 and 109.8 hours respectively.

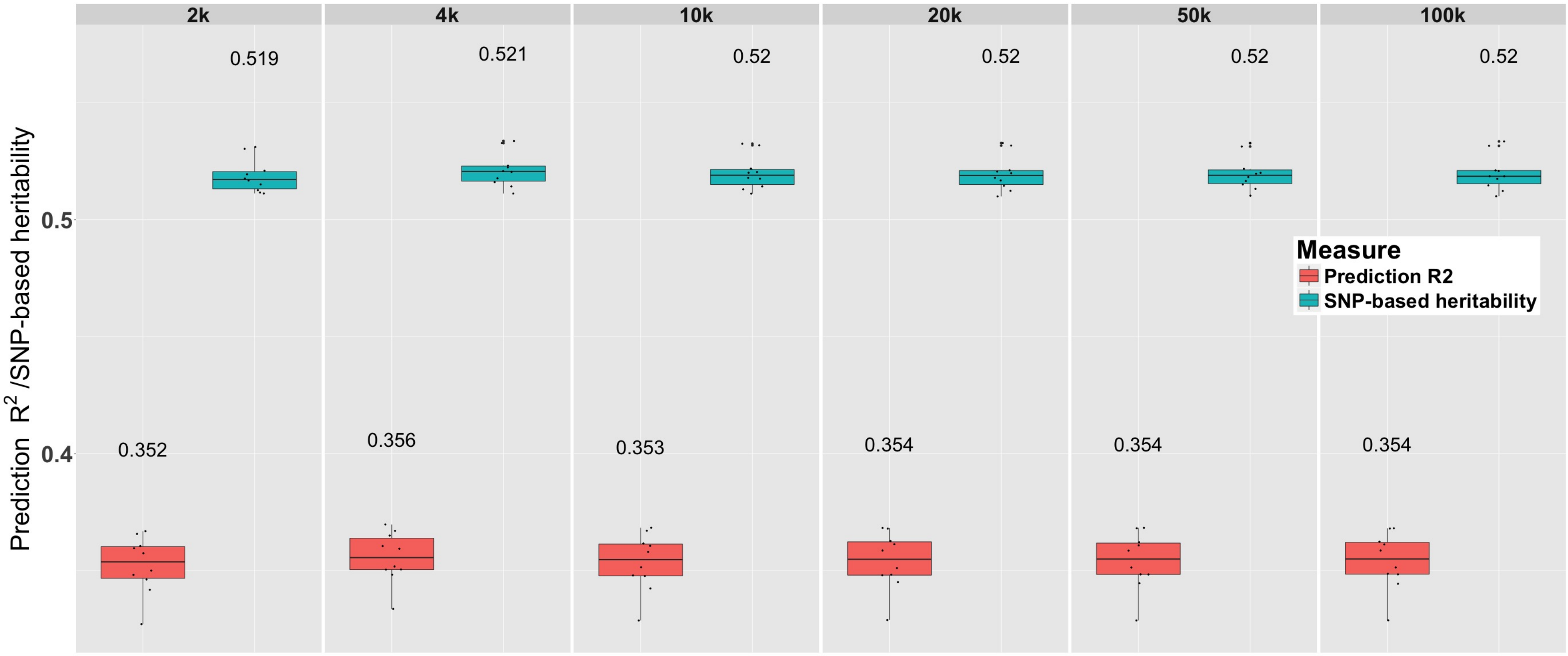


**Figure S9 SBayesR prediction accuracy and SNP-based heritability $(h^2_{SNP})$ estimation change with MCMC chain length for one scenario of UKB genome-wide simulation.** Each panel displays boxplot summaries of prediction $R^2$ and $h^2_{SNP}$ estimates (y-axis) for 2,000 (2k), 4,000 (4k), 10,000 (10k), 20,000 (20k), 50,000 (50k), and 100,000 (100k) MCMC iterations of the SBayesR method. Each boxplot shows the results from the 10 replicates in the 10k (simulated under a BayesR model) causal variant and the true simulated $h^2_{SNP} = 0.5$ scenario. The mean prediction $R^2$ and $h^2_{SNP}$ estimates across the 10 replicates are displayed above the relevant boxplot. The mean run time for each of the 2k, 4k, 10k, 20k, 50k and 100k MCMC iterations scenarios was 0.35, 0.78, 4.4, 4.5, 7.7, and 14.6 hours respectively.

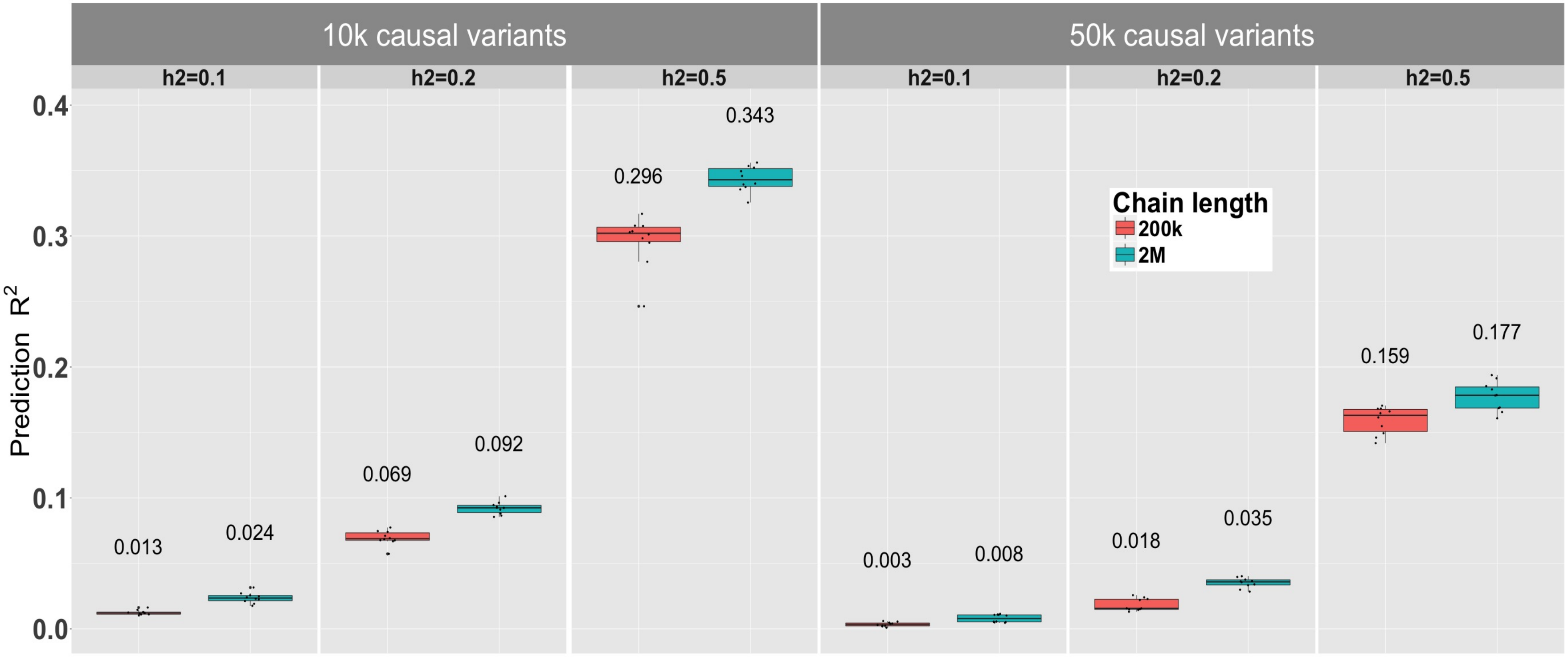


**Figure S10 Regression with Summary Statistics (RSS)[3] prediction accuracy for results generated from 200,000 (200k) and 2,000,000 (2M) iterations of the MCMC chain for all scenarios of the UKB genome-wide simulation.** Each panel displays boxplot summaries of the prediction $R^2$ (y-axis) for RSS across the 10 replicates for each of the six simulation scenarios that varied in the number of causal variants, 10k and 50k, and the true simulated $h^2_{SNP} = (0.1, 0.2, 0.5)$. Two genetic architecture scenarios were generated: 10,000 causal variants sampled under the SBayesR model i.e., 2500, 5000, and 2500 variants from each of $N(0, 0.01\sigma^2_\beta)$, $N(0, 0.1\sigma^2_\beta)$, and $N(0, \sigma^2_\beta)$ distributions respectively and $\sigma^2_\beta = 1$. For the second architecture, 50,000 causal variants were sampled from a standard normal distribution. The mean prediction $R^2$ value across the 10 replicates is displayed above the relevant boxplot.

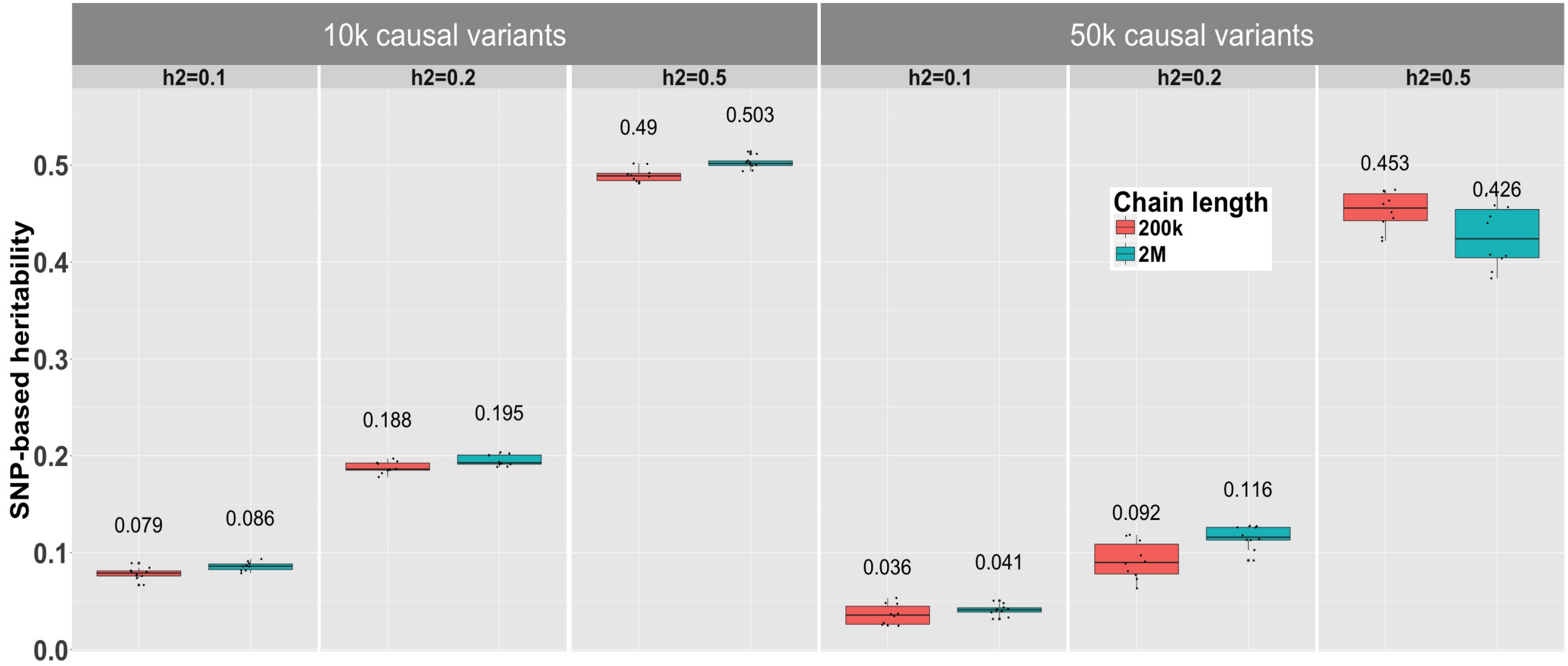


**Figure S11 Regression with Summary Statistics (RSS)[3] SNP-based heritability $(h^2_{SNP})$ estimates for results generated from 200,000 (200k) and 2,000,000 (2M) iterations of the MCMC chain for all scenarios of the UKB genome-wide simulation.** Each panel displays boxplot summaries of $h^2_{SNP}$ estimates (y-axis) for RSS across the 10 replicates for each of the six simulation scenarios that varied in the number of causal variants, 10k and 50k, and the true simulated $h^2_{SNP} = (0.1, 0.2, 0.5)$. Two genetic architecture scenarios were generated: 10,000 causal variants sampled under the SBayesR model i.e., 2500, 5000, and 2500 variants from each of $N(0, 0.01\sigma^2_\beta)$, $N(0, 0.1\sigma^2_\beta)$, and $N(0, \sigma^2_\beta)$ distributions respectively and $\sigma^2_\beta = 1$. For the second architecture, 50,000 causal variants were sampled from a standard normal distribution. The mean prediction $h^2_{SNP}$ estimate across the 10 replicates is displayed above the relevant boxplot.

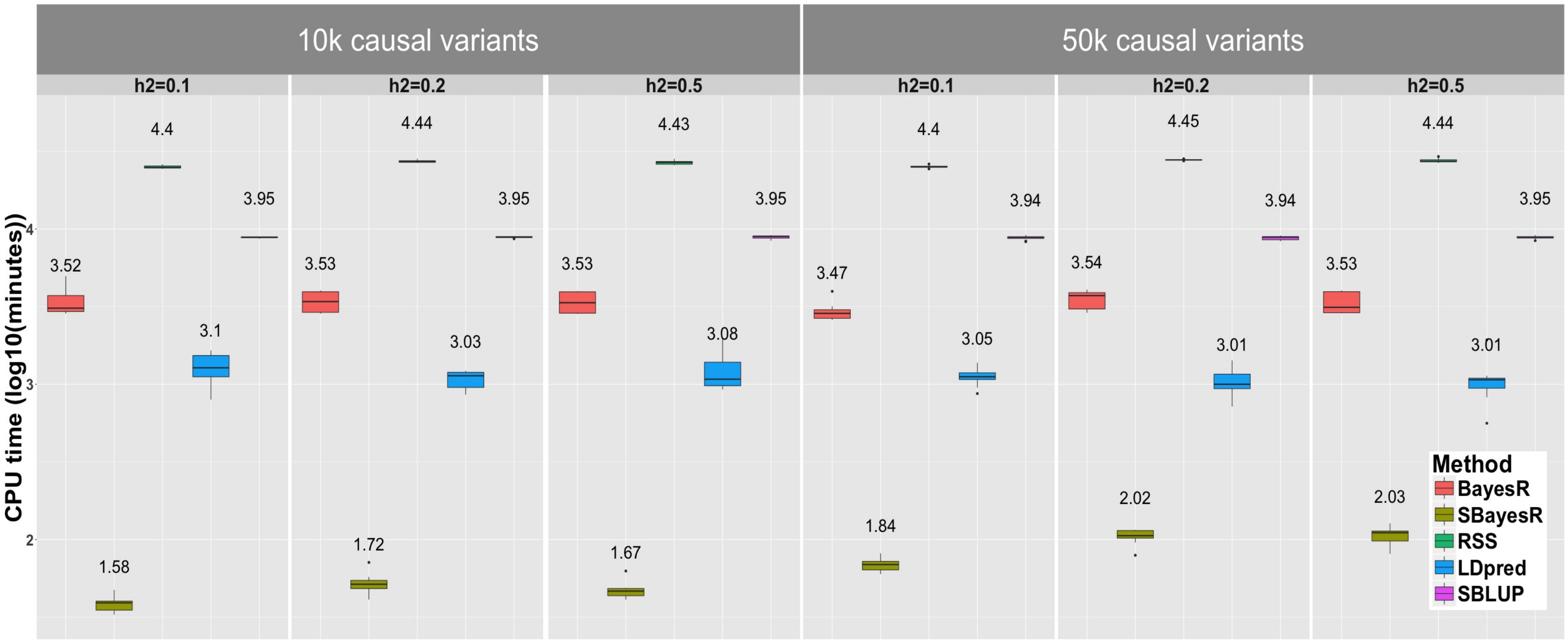


**Figure S12 Runtime** ($\log_{10}$**(minutes)) comparison for BayesR, SBayesR, RSS, LDpred and SBLUP for UKB genome-wide simulation.** Each panel shows a boxplot summary of runtime across the 10 replicates for each scenario with the mean runtime displayed above each method's boxplot. The runtime for RSS, LDpred and SBLUP represents the sum over the runtimes for each chromosome. Results for P+T, HEreg and LDSC are not shown as they required relatively minimal computing resources.

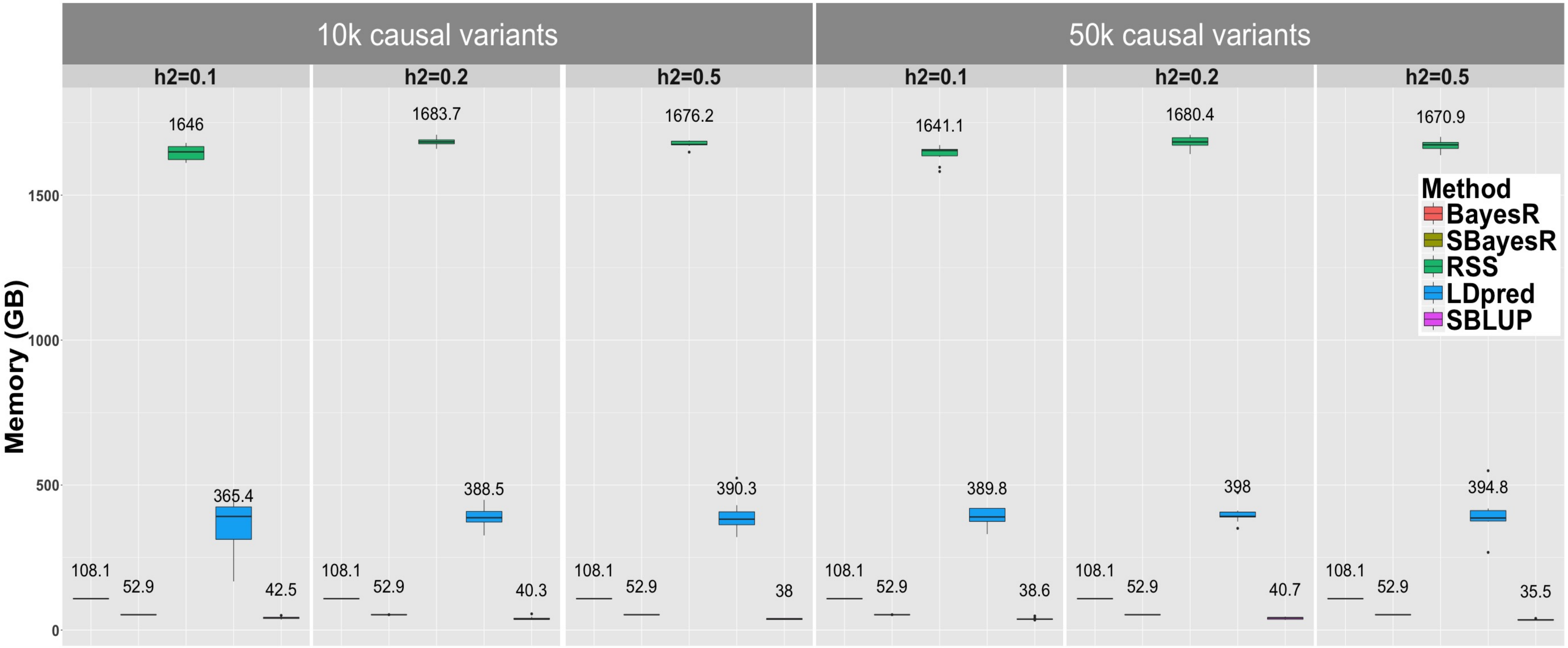


**Figure S13 Memory usage in gigabytes (GB) comparison for BayesR, SBayesR, RSS, LDpred and SBLUP for UKB genome-wide simulation.** Each panel shows a boxplot summary of memory usage across the 10 replicates for each scenario with the mean memory displayed above each method's boxplot. The memory for RSS, LDpred and SBLUP represents the sum over the memory usage for each chromosome. Results for P+T, HEreg and LDSC are not shown as they required relatively minimal computing resources.

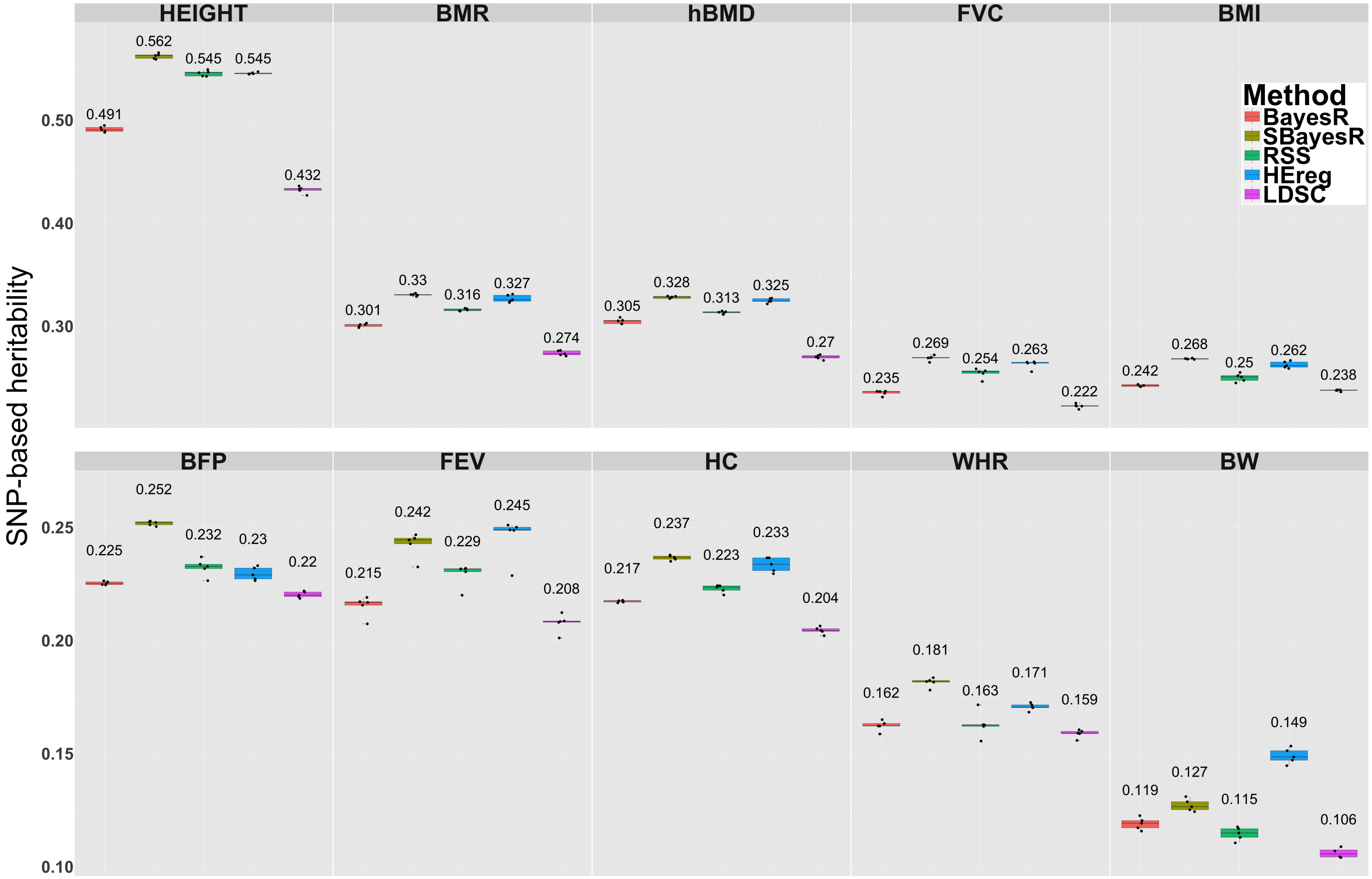


**Figure S14 SNP-based heritability $(h^2_{SNP})$ estimation performance for different methods in the 5-fold cross-validation analysis of 10 quantitative traits in the UKB.** Panel headings describe the abbreviation for the 10 quantitative traits including: standing height (HEIGHT), basal metabolic rate (BMR), heel bone mineral density T-score (hBMD), forced vital capacity (FVC), body mass index (BMI), body fat percentage (BFP), forced expiratory volume in one-second (FEV), hip circumference (HC), waist-to-hip ratio (WHR) and birth weight (BW). Each panel shows a boxplot summary of the $h^2_{SNP}$ estimates across the five folds with the mean across the five folds displayed above each method's boxplot.

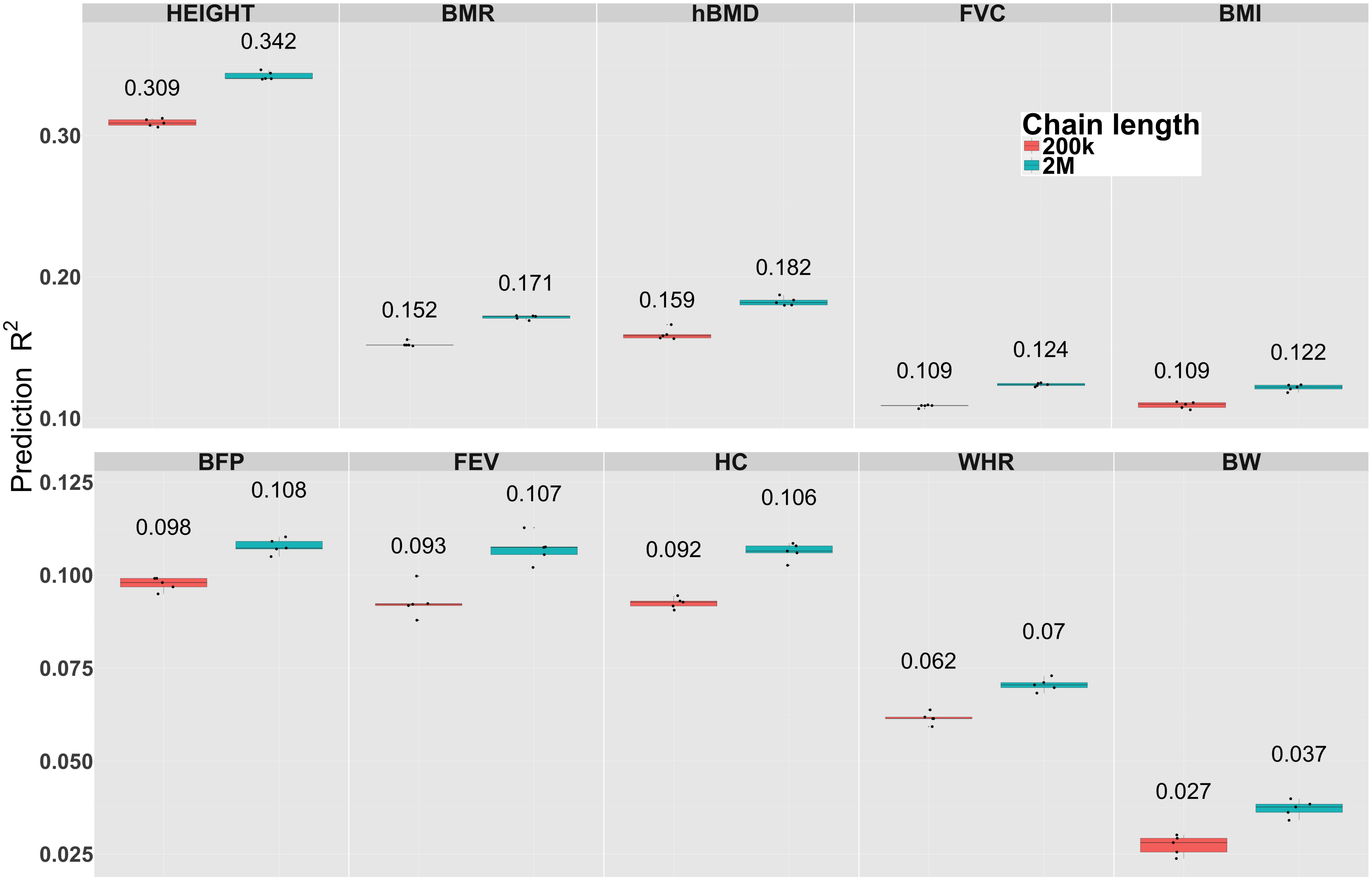


**Figure S15 Regression with Summary Statistics (RSS)[3] prediction accuracy for results generated from 200,000 (200k) and 2,000,000 (2M) iterations of the MCMC chain in the 5-fold cross-validation analysis of 10 quantitative traits in the UKB.** Panel headings describe the abbreviation for the 10 quantitative traits including: standing height (HEIGHT), basal metabolic rate (BMR), heel bone mineral density T-score (hBMD), forced vital capacity (FVC), body mass index (BMI), body fat percentage (BFP), forced expiratory volume in one-second (FEV), hip circumference (HC), waist-to-hip ratio (WHR) and birth weight (BW). Each panel shows a boxplot summary of the prediction $R^2$ across the five folds with the mean across the five folds displayed above each boxplot.

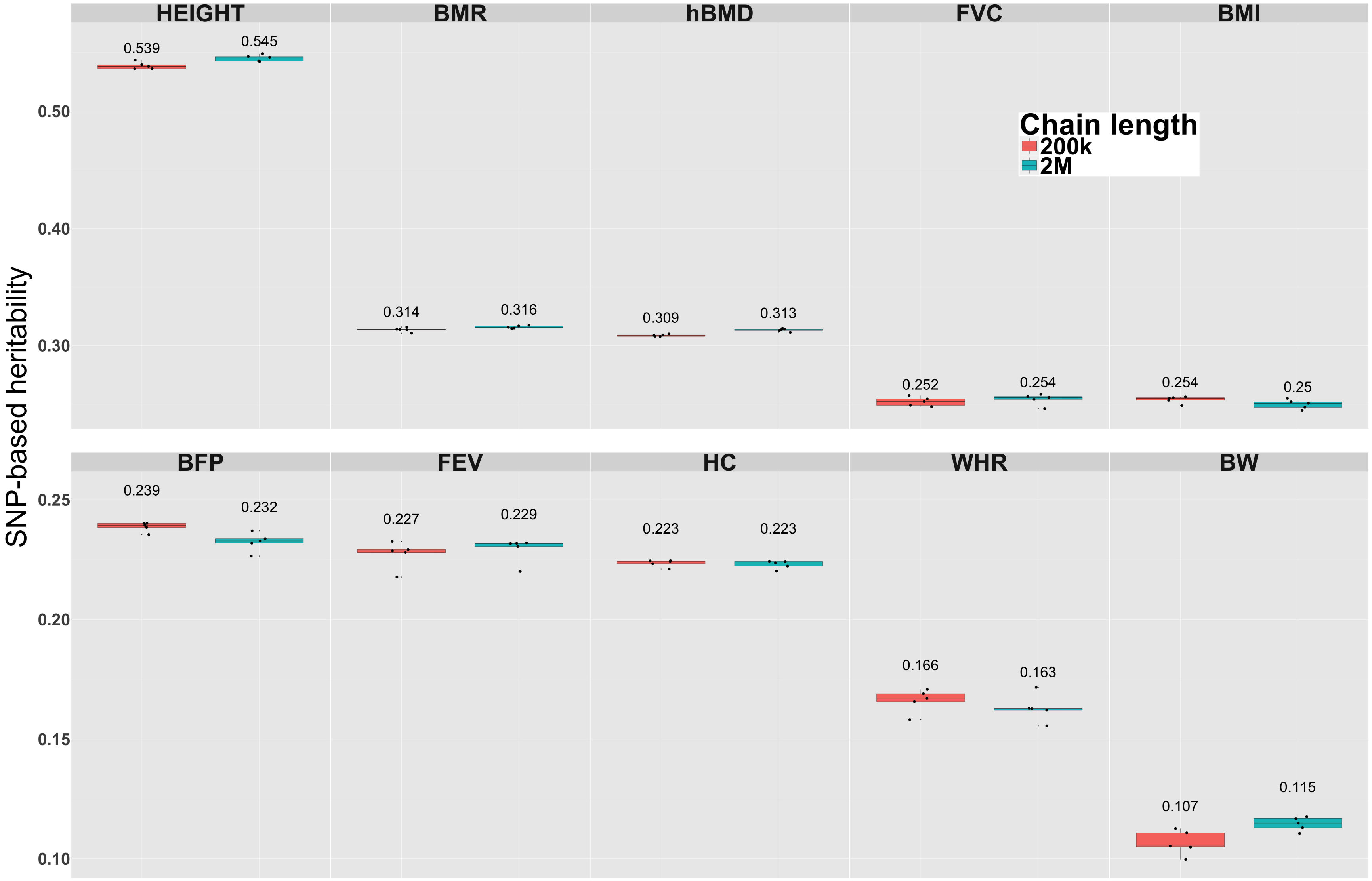


**Figure S16 Regression with Summary Statistics (RSS)[3] SNP-based heritability ($h^2_{SNP}$) estimates for results generated from 200,000 (200k) and 2,000,000 (2M) iterations of the MCMC chain in the 5-fold cross-validation analysis of 10 quantitative traits in the UKB.** Panel headings describe the abbreviation for the 10 quantitative traits including: standing height (HEIGHT), basal metabolic rate (BMR), heel bone mineral density T-score (hBMD), forced vital capacity (FVC), body mass index (BMI), body fat percentage (BFP), forced expiratory volume in one-second (FEV), hip circumference (HC), waist-to-hip ratio (WHR) and birth weight (BW). Each panel shows a boxplot summary of the $h^2_{SNP}$ estimates across the five folds with the mean across the five folds displayed above each boxplot.

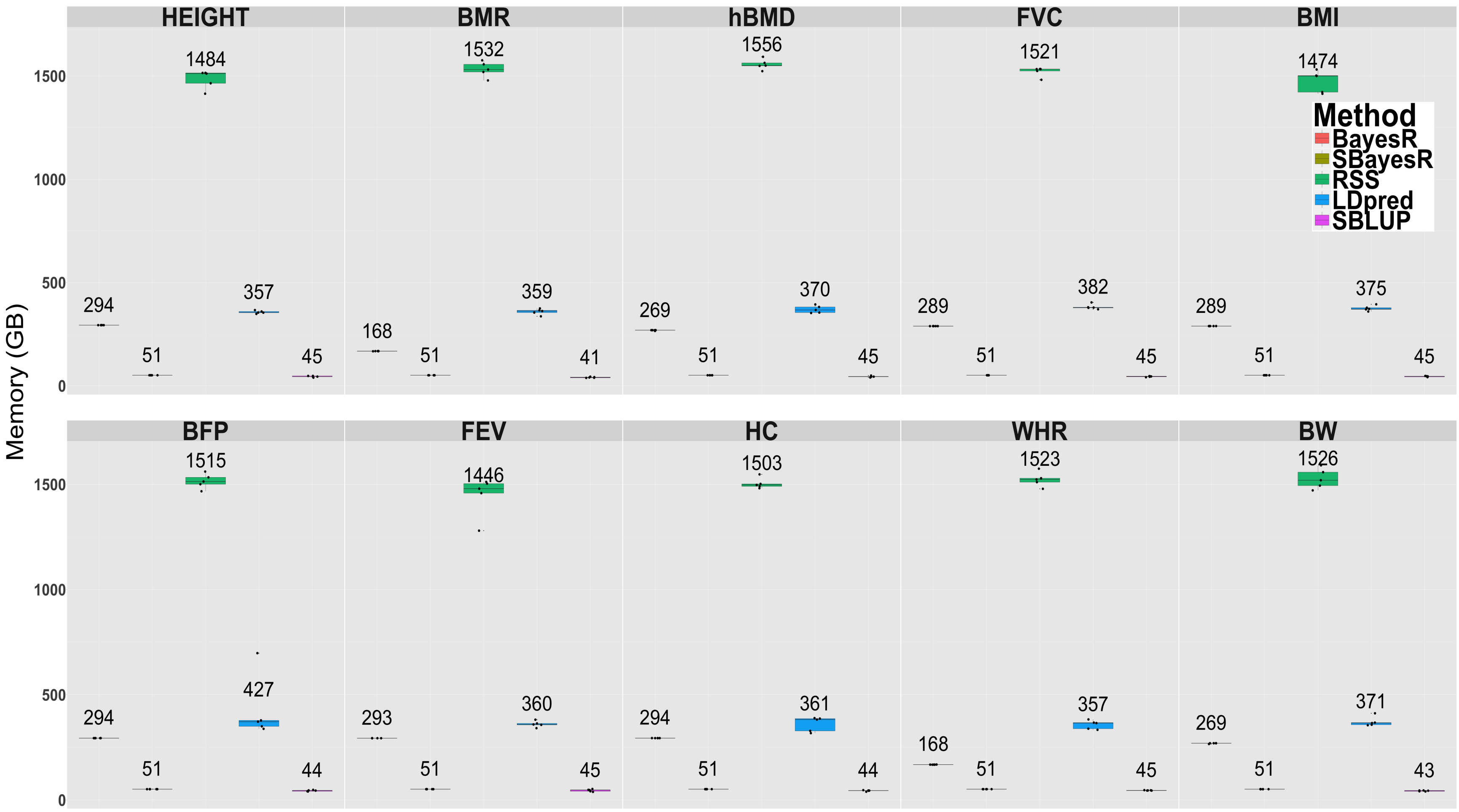


**Figure S17 Memory usage comparison in gigabytes (GB) for cross validation analysis of 10 quantitative traits in the UKB.** Panels headings describe the abbreviation for the 10 quantitative traits. Each panel shows a boxplot summary of memory usage across the five folds with the mean across the five folds displayed above each method's boxplot. Results for RSS, LDpred and SBLUP represent the sum over memory for each chromosome-wise analysis. Results for RSS and SBayesR do not include the memory required to compute the LD reference matrix. See Figure S14 for description of trait abbreviations. Results for HEreg and LDSC are not shown as they required relatively minimal computing resources.

### Supplemental Note

#### Simulation study on two chromosomes

##### *Description*

To initially investigate the performance of the SBayesR methodology, we simulated quantitative phenotypes using 30,122 HM3 variants from chromosomes 21 and 22 for a random subset of 100,000 individuals from the 348,580 unrelated Europeans in the version three UKB data set. The variants on chromosomes 21 and 22 were taken from the list of 1,365,446 HM3 SNPs, which included a final filter that excluded SNPs with MAC $\geqslant 5$ and pHWE $< 1 \times 10^{-5}$ and missingness $> 0.05$, in the UKB data set. Taking the overlap between the HM3 variants on these chromosomes and the 1000G genetic map downloaded from joepickrell/1000-genomes-genetic-maps left 30,122 variants. The 1000G genetic map is required for use in the LD matrix shrinkage estimator of Wen and Stephens[4]. The genetic map files contain interpolated map positions for the CEU population generated from the 1000G OMNI arrays. The shrinkage estimator of the LD matrix, shrinks the off-diagonal entries of the an LD correlation matrix toward zero and is required for the Regression with Summary Statistics (RSS) method[3].

Using these genotypes, three genetic architecture scenarios were generated under the multiple regression model $y_i = \sum_{j=1}^{p} w_{ij}\beta_j + \varepsilon_i$, where $w_{ij} = (x_{ij} - 2p_j)/\sqrt{2p_j(1-p_j)}$ with $x_{ij}$ being the reference allele count for the $i$th individual at the $j$th SNP, $p_j$ the allele frequency of the $j$th variant and $\varepsilon_i$ was sampled from a normal distribution with mean 0 and variance $\text{Var}(\mathbf{W}\boldsymbol{\beta})(1/h^2_{SNP} - 1)$ such that $h^2_{SNP} = 0.1$ for each simulation replicate, which is larger than the contribution to the genome-wide SNP-based heritability ($h^2_{SNP}$) estimate for these chromosomes for most quantitative traits. All phenotypes were generated using the R programming language.

For each scenario replicate, we randomly sampled a new set of 1,500 causal variants, which is the approximate number of causal variants that are expected given the proportion of HM3 variants on chromosomes 21 and 22 and a trait with 50,000 genome-wide causal variants. The first genetic architecture (GA1) contained two causal variants of large effect explaining 3% and 2% of the phenotypic variance respectively and a polygenic tail of 1,498 causal variants sampled from a N(0, 0.05/1, 498) distribution such that the expected total genetic variance explained by all variants was 0.1. The second architecture (GA2) was simulated under a BayesR model with three sets of causal variants: the first contained 1,445 causal variants sampled from a $N(0, 0.0.06/1445)$ distribution, the second contained 50 causal variants sampled from a $N(0, 0.02/50)$ distribution and the third five causal variants sampled from $N(0, 0.02/5)$ distribution. The third architecture (GA3) contained 1,500 variants sampled from a $N(0, 0.1/1500)$ distribution. For each of the three genetic architecture scenarios, 10 simulation replicates were generated for the 100,000 individuals. We generated two independent tuning and validation genotype sets from the remaining 248,580 unrelated European individuals each containing 10,000 individuals. The tuning genotype data set is required for parameter tuning, for example, the $p$-value threshold when performing clumping and then $p$-value thresholding Tuning and validation phenotypes were generated using the effects generated from the training data. For each of the 10 simulation replicates in the three scenarios, simple linear regression for each variant was run using the PLINK 2 software[5] to generate summary statistics.

For each of the simulation scenarios the following methods were applied: LDpred[6], RSS[3], summary BLUP (SBLUP)[7], LD clumping and then $p$-value thresholding (P+T) implemented in PLINK 2, individual data BayesR[2] and the summary data implementation of BayesR (SBayesR) implemented in the GCTB software. For $h^2_{SNP}$ comparison we ran Haseman-Elston regression (HEreg) in the GCTA software[8–10]. LD score regression (LDSC)[11] was not run for $h^2_{SNP}$ estimation comparison in this simulation due to the small number of variants used, which can lead to unreliable estimates. The SBayesR and RSS methods require precomputed reference LD correlation matrices.

To assess the influence of LD data reference on prediction performance and parameter estimation, we generated LD correlation matrices for the 30,122 HM3 variants using genotypes from the 1000G, ARIC and UK10K cohorts and two random subsamples from the UKB genotype data with 5,000 (UKB5K) and 50,000 (UKB50K) individuals. The overlap between these random subsamples with the 100,000 random individuals from the GWAS was 1,467 and 13,967 individuals respectively. For each LD reference cohort chromosome-wise LD matrices i.e., all inter chromosomal LD is ignored, were built and the shrinkage estimator of the LD matrix[4] calculated using an efficient implementation in the GCTB software. The calculation of the shrunk LD matrix requires the effective population sample size, which we set to be 11,400 (as in Zhu and Stephens[3]), the sample size of the genetic map reference, which corresponds to the 183 individuals from the CEU cohort of the 1000G and the hard threshold on the shrinkage value, which we set to $10^{-3}$. This threshold gave a good balance between computational efficiency and accuracy with on average each SNP having 3,033 (SD=756) non-zero elements across these two chromosomes (Figure S5). We further stored the shrunk LD matrix in sparse matrix format (ignoring matrix elements equal to 0) for efficient SBayesR computation. SBayesR was run for each of the simulation scenarios using each reference LD matrix.

The PLINK 2 software was used to calculate the estimate genetic values for each individual for all LD matrix cohorts and the prediction $R^2$, calculated via linear regression of the true simulated phenotype in the validation data set on that predicted values from SBayesR, used as a measure of prediction accuracy. The UKB50K reference showed marginal improvements over the other cohorts in prediction accuracy in the validation data set and had the smallest upward bias in $h^2_{SNP}$ estimation (Figure S1). We therefore selected this LD reference cohort for all methods. For LDpred, SBLUP and P+T, a separate genotype data set is required to be specified for LD correlation reference and utilisation within each method's program. This was set to be the same 50,000 individual genotype set used for SBayesR and RSS. Furthermore, the full LD matrix that incorporates inter-chromosomal LD information was generated using the full 100,000 individuals such that the individual data BayesR model could be compared with the SBayesR model run with the full LD matrix, which is expected to produce equivalent results.

For LDpred, we specified $h^2_{SNP}$ to be equal to the true 0.1, specified the number of SNPs on each side of the focal SNP for which LD should be adjusted to be 3,500, which equated to an approximate 10 megabase (MB) window, and calculated effects size estimates for all of the 10 fraction of non-zero effects pre-specified parameters, which included LDpred-inf, 1, 0.3, 0.1, 0.03, 0.01, 0.003, 0.001, 0.0003, and 0.0001. A larger LD window size than recommended was chosen because of the large effects simulated, it was computationally feasible in this small simulation and to be comparable with the LD references used for SBayesR and RSS. For RSS, analyses were performed for each chromosome to limit the computational burden of running these analyses using MATLAB, as in Zhu and Stephens[3]. For each chromosome, the RSS-BSLMM model was run for 2 million MCMC iterations with 1 million as burn in and a thinning rate of 1 in 100 to arrive at 10,000 posterior samples for each of the model parameters. For each chromosome, the posterior mean over posterior samples for the SNP effects and $h^2_{SNP}$ estimates was used. The chromosome wise $h^2_{SNP}$ estimates were then summed to get the total estimate. For SBLUP, we used the GCTA software implementation, which required the specification of the $\lambda = m(1/h^2_{SNP} - 1)$ parameter, which was calculated using $h^2_{SNP} = 0.1$ and $m = 30,122$ and the LD correlation window size specification was set to 10 MB. For P+T, we used the PLINK 2 software to clump the GWAS summary statistics discarding variants within 1 MB (using 10 MB gave very similar results) of and in LD $R^2 > 0.1$ with the most associated SNP in the region. Using these clumped results, we generated PRSs for sets of SNPs at the following $p$-value thresholds: $5\times10^{-8}$, $1\times10^{-6}$, $1\times10^{-4}$, 0.001, 0.01, 0.05, 0.1, 0.2, 0.5, and 1.0. BayesR was run using a mixture of four normal distributions model with distribution variance weights $\gamma = (0, 10^{-4}, 10^{-3}, 10^{-2})'$. BayesR was run for 4,000 iterations with 2,000 taken as burn in and a thinning rate of 1 in 10. The posterior mean of the effects and the proportion of variance explained over the 200 posterior samples was taken as the parameter estimate for each scenario replicate. For SBayesR the MCMC chain was run for 4,000 iterations with 2,000 taken as burn in and a thinning rate of 1 in 10 and run with four distributions and variance weights $\gamma = (0, 0.01, 0.1, 1)'$. HEreg requires a genetic relatedness matrix, which was built from 30,122 HM3 variants from chromosomes 21 and 22 using the GCTA software.

To assess prediction accuracy, the estimated genetic value (EGV) for each individual was calculated using the genotype data from the 10,000 individual tuning and validations data sets and genetic effect estimates from each method. Tuning was performed for LDpred and P+T where for each simulation replicate the prediction accuracy was assessed for each of the pre-specified fraction of non-zero effects parameters for LDpred and the $p$-value thresholds for P+T. The parameter that gave the optimal prediction $R^2$ in the tuning data set was then used for calculating the EGV in the validation data set. SNP effects from BayesR and SBayesR were estimated using scaled genotypes and thus each variant's effect was divided by $\sqrt{2q_j(1-q_j)}$, where $q_j$ is the allele frequency from the validation cohort of the $j$th variant, before PLINK scoring was performed. The PLINK 2 software was used to perform the EGV calculation for all methods and the prediction $R^2$ calculated via linear regression of the true simulated phenotype on that estimated from each method used as a measure of prediction accuracy.

##### *Results*

The choice of LD matrix reference cohort for use in SBayesR analysis led to differences in absolute prediction accuracy and bias in $h^2_{SNP}$ estimation (Figures S1 and S2). The 1000G cohort showed the poorest prediction accuracy and upward bias in on mean $h^2_{SNP}$ estimates. The ARIC and UK10K LD reference cohorts showed similar on mean prediction $R^2$ and bias in $h^2_{SNP}$ estimation as the UKB50K and UKB5K cohorts. However, overall the UKB50K cohort showed the maximum prediction accuracy and smallest upward bias in $h^2_{SNP}$ estimation across all scenarios and thus was chosen as the LD reference cohort for all analyses.

Across the three simulation scenarios we observed that the individual level analysis using BayesR, or SBayesR with the full LD matrix, gave the highest mean validation data set prediction $R^2$ (Figure S3). The relative difference between the individual data BayesR model mean prediction accuracy and the highest performing summary statistics method, SBayesR, ranged from 1.2% to 3.9% (Figure S3). P+T showed the lowest on mean prediction accuracy across scenarios but showed similar mean prediction accuracies to the LDpred infinitesimal model and SBLUP for scenario one, which contains variants of very large effect and a polygenic tail. SBayesR showed substantial improvement in prediction accuracy relative to other summary statistics methodologies particularly in scenario one, with RSS showing the closest prediction $R^2$ compared to SBayesR in all scenarios. RSS outperformed SBLUP and LDpred-inf in all scenarios but showed a smaller relative improvement in prediction $R^2$ as the simulated traits had fewer large effects. SBLUP outperformed LDpred-inf in each of the simulation scenarios, which is the most similar LDpred model to SBLUP (Figure S3). Overall, on mean prediction $R^2$ improvement ranged from 1.3% to 7.9% when comparing SBayesR with the best alternative summary statistic method (RSS) across all scenarios (Figure S3).

Across all simulation scenarios all methods showed minimal bias in $h^2_{SNP}$ estimation (Figure S4). Overall SBayesR using the full LD matrix showed the smallest bias, with HEreg showing the largest bias in scenario one with the bias diminishing as the scenarios became more similar to the infinitesimal model. RSS showed a downward bias in GA1 and was unbiased for GA2 and GA3. SBayesR maintained a marginal upward bias across all simulation scenarios and a maximum relative upward on mean bias of 3% in GA2 and GA3 (Figure S4).

#### Bayesian multiple regression

The starting point is the multiple linear model of the form

$$\mathbf{y} = \mathbf{X}\boldsymbol{\beta} + \boldsymbol{\varepsilon}, \tag{1}$$

where $\mathbf{y}$ is an $n \times 1$ vector (centred) of trait phenotypes, $\mathbf{X}$ is an $n \times p$ matrix of genotype covariates initially coded 0, 1, 2 representing the number of copies of a reference allele at each marker, and we consider that the columns of $\mathbf{X}$ can either be centred or centred and scaled. The vector $\boldsymbol{\beta}$ is a $p \times 1$ vector of random partial regression coefficients of the $p$ SNPs (marker effects) and $\boldsymbol{\varepsilon}$ is a vector ($n \times 1$) of residuals.

We wish to optimise the parameters of the stated linear model using Bayesian posterior inference, which requires the specification of prior distributions for $\boldsymbol{\beta}$ and $\boldsymbol{\varepsilon}$. We assume that the error term $\boldsymbol{\varepsilon}|\sigma_\varepsilon^2 \sim \text{MVN}(\mathbf{0}, \mathbf{R}\sigma_\varepsilon^2)$, where MVN denotes the multivariate normal distribution, $\mathbf{0}$ is a column vector of zeroes of length $n$, and $\mathbf{R}$ is a covariance matrix, which is assumed here to be a diagonal matrix of ones. The parameter $\sigma_\varepsilon^2$ is treated as an unknown with a scaled inverse chi-square distribution prior with scale parameter $s_\varepsilon^2$ and degrees of freedom $\nu_\varepsilon$.

Members of the Bayesian alphabet for genomic selection including BayesA and BayesB[12], BayesC and BayesC$\pi$[13], BayesR[2,14], BSLMM[15], BayesS[16] among others, differ largely in the prior used for $\boldsymbol{\beta}$. In this work, we will focus on the BayesR model, which assumes that $\beta_j$ comes from a finite mixture of normals distribution, which includes a point mass at zero. This prior is motivated by the capacity of the mixture distribution to be flexible and thus model a diverse set of underlying genetic effect distributions.

Inferences on marker associations are based on the posterior distribution of the marker effects $f(\boldsymbol{\beta}|\mathbf{y})$. Closed form expressions are not available for making inferences from $f(\boldsymbol{\beta}|\mathbf{y})$ and instead they are drawn from the posterior of interest. The following derivation describes a similar Markov chain Monte Carlo (MCMC) algorithm as in Habier *et al.*[13], which is discussed in detail in Fernando and Garrick[17].

Let $\boldsymbol{\theta} = (\boldsymbol{\beta}', \boldsymbol{\pi}', \sigma_\beta^2, \sigma_\varepsilon^2)'$ denote all the unknowns in the model including the random marker effects, mixing proportions of the mixture of normals, the variance of the marker effects, and the residual variance. For each model parameter, we draw posterior samples using the single site Gibbs sampler, which draws samples for each element $i$ of the vector $\boldsymbol{\theta}$ from its full conditional posterior: $f(\theta_i|\boldsymbol{\theta}_{-i}, \mathbf{y})$. The full conditional can be expressed as

$$f(\theta_i|\boldsymbol{\theta}_{-i}, y) \propto f(\theta_i, \boldsymbol{\theta}_{-i}, \mathbf{y}). \tag{2}$$

The joint density in (2) can be written as

$$f(\theta_i, \boldsymbol{\theta}_{-i}, \mathbf{y}) = f(\mathbf{y}|\boldsymbol{\theta})f(\theta_i)f(\boldsymbol{\theta}_{-i}),$$

where $f(\mathbf{y}|\boldsymbol{\theta})$ is the density function of the conditional distribution of $\mathbf{y}|\boldsymbol{\theta}$, and $f(\theta_i)$ and $f(\boldsymbol{\theta}_{-i})$ are the densities of the prior distributions of $\theta_i$ and $\theta_{-i}$. Ignoring factors that are constant with respect to $\theta_i$ gives the kernel of the full-conditional posterior for each parameter of interest, which we will derive for each element of $\boldsymbol{\theta}$.

The conditional distribution of $\mathbf{y}$ given all the unknowns is MVN with expectation $\mathbf{X}\boldsymbol{\beta}$ and covariance matrix $\mathbf{R}\sigma_e^2$. The MVN density is thus

$$f(\mathbf{y}|\boldsymbol{\theta}) = (2\pi\sigma_e^2)^{-n/2} \exp\left[-\frac{(\mathbf{y}-\mathbf{X}\boldsymbol{\beta})'(\mathbf{y}-\mathbf{X}\boldsymbol{\beta})}{2\sigma_e^2}\right]. \tag{3}$$

Formally, under the BayesR model we assume the following prior on the genetic effects

$$\beta_j|\boldsymbol{\pi}, \sigma_\beta^2 = \begin{cases} 0 & \text{with probability } \pi_1, \\ \sim N(0, \gamma_2\sigma_\beta^2) & \text{with probability } \pi_2, \\ \vdots & \\ \sim N(0, \gamma_C\sigma_\beta^2) & \text{with probability} 1 - \sum_{c=1}^{C-1} \pi_c, \end{cases}$$

where $C$ denotes the maximum number of components in the finite mixture model, which is prespecified. The $\gamma_c$ coefficients are prespecified and constrain how the common marker effect variance $\sigma_\beta^2$ scales in each distribution. For example it is common in BayesR to assume $C = 4$ such that $\boldsymbol{\gamma} = (\gamma_1, \gamma_2, \gamma_3, \gamma_4)' = (0, 0.0001, 0.001, 0.01)'$ representing a class of effects with no effect and three further classes of small, medium and large effects. Under this prior assumption, we derive the MCMC Gibbs sampling routine for sampling of the key model parameters $\boldsymbol{\theta} = (\boldsymbol{\beta}', \boldsymbol{\pi}', \sigma_\beta^2, \sigma_\varepsilon^2)'$ from their full conditional distributions.

We introduce the dummy variable $\delta_j$ which is a random variable that takes values $1, 2, \ldots, C$ depending on which mixture distribution marker $j$ is sampled in. The prior distribution for the marker effects conditional on the marker effect variance $\sigma_\beta^2$ and mixture class is

$$f(\boldsymbol{\beta}|\delta = c, \gamma_c\sigma_\beta^2) = \prod_{j=1}^{k_c} (2\pi\gamma_c\sigma_\beta^2)^{-1/2} \exp\left[-\frac{\beta_j^2}{2\gamma_c\sigma_\beta^2}\right],$$

which represents the product over those markers sampled in class $c$ denoted $k_c$. We assume that the prior for $\sigma_\beta^2$ is a scaled inverse chi-square distribution with density

$$f(\sigma_\beta^2; \nu_\beta, S_\beta^2) = \frac{(S_\beta^2 \nu_\beta/2)^{\nu_\beta/2}}{\Gamma(\nu_\beta/2)} \frac{\exp(-\nu_\beta S_\beta^2/2\sigma_\beta^2)}{(\sigma_\beta^2)^{1+\nu_\beta/2}},$$

where $S_\beta^2$ and $\nu_\beta$ are the scale parameter and degrees of freedom respectively. As stated above the residual variance $\sigma_\varepsilon^2$ is assumed to have scaled inverse chi-square distribution prior with distribution

$$f(\sigma_e^2; \nu_\varepsilon, S_\varepsilon^2) = \frac{(S_\varepsilon^2 \nu_\varepsilon/2)^{\nu_\varepsilon/2}}{\Gamma(\nu_\varepsilon/2)} \frac{\exp(-\nu_\varepsilon S_\varepsilon^2/2\sigma_e^2)}{(\sigma_e^2)^{1+\nu_\varepsilon/2}}.$$

The full conditional posterior of $\beta_j$ is proportional to the product of the likelihood, the prior distribution for $\beta_j$, and the prior distributions of the variances. The variances don't contain $\beta_j$ and nor do the other components of the product for the prior for $\beta_j$. Therefore, the full conditional for $\beta_j$ can be written as

$$f(\beta_j|\delta_j = c, \boldsymbol{\theta}_{-\beta_j}, \mathbf{y}) \propto \exp\left[-\frac{(\mathbf{y}-\mathbf{X}\boldsymbol{\beta})'(\mathbf{y}-\mathbf{X}\boldsymbol{\beta})}{2\sigma_e^2}\right] \exp\left[-\frac{\beta_j^2}{2\gamma_c\sigma_\beta^2}\right].$$

We define $\mathbf{w} = \mathbf{y} - \sum_{k\neq j} \mathbf{x}_k\beta_k$ to be the vector of trait phenotypes corrected for all effects other than that being sampled. Given this we can write

$$\begin{aligned} f(\beta_j|\delta_j = c, \boldsymbol{\theta}_{-\beta_j}, \mathbf{y}) &\propto \exp\left\{-\frac{1}{2\sigma_e^2}\left[(\mathbf{w}-\mathbf{x}_j\beta_j)'(\mathbf{w}-\mathbf{x}_j\beta_j) + \frac{\beta_j^2\sigma_e^2}{2\gamma_c\sigma_\beta^2}\right]\right\} \\ &\propto \exp\left[-\frac{1}{2\sigma_e^2}\left(\mathbf{w}'\mathbf{w} - 2\mathbf{x}_j'\mathbf{w}\beta_j + \mathbf{x}_j'\mathbf{x}_j\beta_j^2 + \frac{\beta_j^2\sigma_e^2}{\sigma_\beta^2}\right)\right]. \end{aligned} \tag{4}$$

We can complete the square with respect to $\beta_j$ in (4) to obtain

$$f(\beta_j|\delta_j = c, \boldsymbol{\theta}_{-\beta_j}, \mathbf{y}) \propto \exp\left[-\frac{1}{2\sigma_e^2}\left(\mathbf{w}'\mathbf{w} - l_{jc}\widehat{\beta}_j^2 + l_{jc}(\beta_j - \widehat{\beta}_j)^2\right)\right],$$

where $l_{jc} = \mathbf{x}_j'\mathbf{x}_j + \sigma_e^2/(\gamma_c\sigma_\beta^2)$ is the left hand side of the well known mixed model equations

(MME)[18] for $\beta_j$, $\widehat{\beta}_j = \mathbf{x}_j'\mathbf{w}/l_{jc}$, and $\mathbf{x}_j'\mathbf{w}$ is the right hand side of the MME. Dropping terms that are free from $\beta_j$ the full conditional becomes

$$f(\beta_j|\delta_j = c, \boldsymbol{\theta}_{-\beta_j}, \mathbf{y}) \propto \exp\left[-\frac{1}{2}\frac{(\beta_j - \widehat{\beta}_j)^2}{\frac{\sigma_e^2}{l_{jc}}}\right].$$

This can be seen to be the kernel of the normal distribution and within each iteration of the Gibbs sampler we sample the genetic effect from a normal distribution with mean $\widehat{\beta}_j$ and variance $\sigma_e^2/l_{jc}$.

In BayesR the prior assumption is that the marker effects have IID Gaussian mixture distributions, with a point mass at zero with probability $\pi_1$, a univariate normal distribution with variance $\gamma_2\sigma_\beta^2$ with probability $\pi_2$, a univariate normal distribution with variance $\gamma_3\sigma_\beta^2$ with probability $\pi_3$ etc. up to a univariate normal distribution with variance $\gamma_C\sigma_\beta^2$ with probability $\pi_C$ , such that $\pi_C = 1 - \sum_{c=1}^{C-1}\pi_c$, where $C$ here denotes the maximum number of components in the finite mixture model. The vector $\boldsymbol{\pi} = (\pi_1, \pi_2, \ldots, \pi_C)'$ is treated as an unknown and is assumed to have a Dirichlet prior, which is the extension of the concept in BayesC$\pi$[13] that the $\pi$ is treated as an unknown with a uniform prior. BayesR classically assumes that there are $C = 4$ classes but $C$ can be chosen to be arbitrarily large with some scaling (not necessarily an exponential function) of each of the variance components assumed.

To derive the posterior update for $\boldsymbol{\pi}$, we treat the indicator variables $\delta_j$ as a random variable that takes values $1, 2, 3, \ldots, C$ depending on which class marker $j$ is sampled in. Therefore, $\delta_j$ can be modelled as a categorical random variable, which implies that

$$f(\delta_j|\boldsymbol{\pi}) = \prod_{c=1}^{C}\pi_c^{[\delta_j = c]} \quad ,$$

where $[\delta_j = c]$ evaluates to 1 if $\delta_j = c$ and 0 otherwise. Therefore,

$$f(\delta|\boldsymbol{\pi}) = \prod_{j=1}^{p}\prod_{c=1}^{C}\pi_c^{[\delta_j = c]}.$$

The Dirichlet distribution is the conjugate prior distribution of the categorical distribution. Given, $\delta_j$ has a categorical distribution if we assume that $\boldsymbol{\pi}$ has a $Dirichlet(1,1,\ldots,1)$ prior, which assumes that the prior probability of a SNP being in any distribution is the same, then the posterior distribution of $\delta_j$ is also Dirichlet. Then, more generally in our setting $\boldsymbol{\alpha} = (\alpha_1, \ldots, \alpha_c, \ldots, \alpha_C)$ where $\alpha_c = 1$ and we can write our prior as $\boldsymbol{\pi}|\boldsymbol{\alpha} \sim Dirichlet(C, \boldsymbol{\alpha})$. Given an initial vector $\boldsymbol{\pi}$ then $\boldsymbol{\delta}|\boldsymbol{\pi} \sim Categorical(C, \boldsymbol{\pi})$. From the form of the Dirichlet distribution we have

$$f(\pi_1, ...., \pi_C; \alpha_1, \ldots, \alpha_C) = \frac{1}{B(\boldsymbol{\alpha})} \prod_{c=1}^{C} \pi_c^{\alpha_c - 1}.$$

As always we can ignore the normalising constant and look at

$$f(\boldsymbol{\pi}|\boldsymbol{\delta}, \boldsymbol{\alpha}) \propto f(\boldsymbol{\delta}|\boldsymbol{\pi}) f(\boldsymbol{\pi}|\boldsymbol{\alpha}) = \prod_{j=1}^{p} \prod_{c=1}^{C} \pi_c^{[\delta_j = c]} \prod_{c=1}^{C} \pi_c^{\alpha_c - 1} = \prod_{j=1}^{p} \prod_{c=1}^{C} \pi_i^{[\delta_j = c] + \alpha_c - 1},$$

which is the kernel of a $Dirichlet(C, \mathbf{c} + \boldsymbol{\alpha})$, where $\mathbf{c}$ is a vector of length $C$ with the count of the number of variants in each class and $\mathbf{c} + \boldsymbol{\alpha}$ is a vector with elements $(c_1 + \alpha_1, c_2 + \alpha_2, \ldots, c_C + \alpha_C)$. Therefore, in the Gibbs sampler we sample $\boldsymbol{\pi}$ from a $Dirichlet(C, \mathbf{c} + \boldsymbol{\alpha})$ conditional on the number of variants sampled in each of the $C$ mixture classes.

[14] do not sample the marker effect variance $\sigma^2_\beta$ but instead scale and centre the genotypes and equate the genetic variance $\sigma^2_g = m\sigma^2_\beta$, where $m$ is the number of causal loci[19] and substitute a pre-estimated value of $\sigma^2_g$, from a previous $h^2_{SNP}$ study.[2] also equate the genetic variance with the marker effect variance and sample $\sigma^2_g$ from a scaled inverse chi-square distribution with parameters $\nu_0 + m_g$ and $\frac{m_g \sum_{j=1}^{p} \beta_j^2 + \nu_0 S_0^2}{\nu_0 + m_g}$, where $m_g$ is the number of SNPs included in the current model. Moser *et al.*[2] specify prior values of $\nu_0$ and $S_0^2$ are to be $-2$ and 0, which are proposed to lead to a uninformative prior. For polygenic traits this is likely to be reasonable but a more general hypothesis is to not to connect the genetic variance with the marker effect variance under this assumption.

To derive a new update, we note that the common marker effect variance is only present in the normal density functions of $\beta_j$ when $\delta_j \neq 1$ and its own prior. Therefore,

$$f(\boldsymbol{\beta}|\sigma_\beta^2, \boldsymbol{\delta}) = \prod_{j=1}^{q} \phi(\beta_j; 0, \gamma_{\delta_j}\sigma_\beta^2),$$

where $\boldsymbol{\delta} = (\delta_2, \ldots, \delta_p)$, $\delta_j \in (1, 2, \ldots, C)$, $\phi$ is the normal probability density function and the number of non-zero effects in the model $q = |\boldsymbol{\beta}_{-\beta_j:\delta_j=1}|$. Given this

$$\begin{aligned} f(\sigma_\beta^2; \nu_\beta, S_\beta^2) f(\boldsymbol{\beta}|\sigma_\beta^2, \boldsymbol{\delta}) &= \frac{(S_\beta^2 \nu_\beta/2)^{\nu_\beta/2}}{\Gamma(\nu_\beta/2)} \frac{\exp(-\nu_\beta S_\beta^2/\sigma_\beta^2)}{(\sigma_\beta^2)^{1+\nu_\beta/2}} \prod_{j=1}^{q} (2\pi\gamma_{\delta_j}\sigma_\beta^2)^{-1/2} \exp\left[-\frac{\beta_j^2}{2\gamma_{\delta_j}\sigma_\beta^2}\right] \\ &= \frac{(S_\beta^2 \nu_\beta/2)^{\nu_\beta/2}}{\Gamma(\nu_\beta/2)} \frac{\exp(-\nu_\beta S_\beta^2/2\sigma_\beta^2)}{(\sigma_\beta^2)^{1+\nu_\beta/2}} \gamma_{\delta_2}^{-c_2/2} \ldots \gamma_{\delta_C}^{-c_C/2} (2\pi\sigma_\beta^2)^{-q/2} \exp\left[-\frac{1}{2\sigma_\beta^2}\sum_{j=1}^{q}\frac{\beta_j^2}{\gamma_{\delta_j}}\right], \end{aligned}$$

where $(c_2, \ldots, c_C)$ are the number of variants in each of the non-zero classes. Retaining only those elements that contain $\sigma_\beta^2$

$$\begin{aligned} &\propto \frac{\exp(-\nu_\beta S_\beta^2/2\sigma_\beta^2)}{(\sigma_\beta^2)^{1+\nu_\beta/2}} (\sigma_\beta^2)^{-q/2} \exp\left[-\frac{\sum_{j=1}^{q}\beta_j^2}{2\gamma_{\delta_j}\sigma_\beta^2}\right] \\ &\propto \exp\left[-\frac{1}{2\sigma_\beta^2}\left(\nu_\beta S_\beta^2 + \sum_{j=1}^{q}\frac{\beta_j^2}{\gamma_{\delta_j}}\right)\right] (\sigma_\beta^2)^{-1-\nu_\beta/2-q/2}, \end{aligned}$$

which is the kernel of a scale inverse chi-squared distribution with degrees of freedom $\nu_\beta + q$, where $q$ is the number of non-zero markers in the model. The scale parameter can be determined by letting $\widetilde{\nu}_\beta = \nu_\beta + q$. The expression inside must be equal to $\widetilde{\nu}_\beta \widetilde{S}_\beta^2 = \nu_\beta S_\beta^2 + \sum_{j=1}^{q} \frac{\beta_j^2}{\gamma_{\delta_j}}$. The new scale parameter is thus now

$$\widetilde{S}_\beta^2 = \frac{\nu_\beta S_\beta^2 + \sum_{j=1}^{q} \frac{\beta_j^2}{\gamma_{\delta_j}}}{\nu_\beta + q}.$$

This is only equivalent to that presented in Moser *et al.*[2] when each $\gamma_{\delta_j}$ is equal to $1/q$.

##### ***Joint sampling of $\delta_j$ and $\beta_j$***

We employ a similar strategy as[16] to jointly sample $\delta_j$ and $\beta_j$ by first sampling $\delta_j$ unconditional on $\beta_j$ and then sample $\beta_j$ conditional on $\delta_j$. Mathematically

$$f(\beta_j, \delta_i | \boldsymbol{\theta}_{-\beta_j,\delta_j}, \mathbf{y}) = f(\beta_j | \delta_j, \boldsymbol{\theta}_{-\beta_j}, \mathbf{y}) f(\delta_j | \boldsymbol{\theta}_{-\delta_j}, \mathbf{y}),$$

and then sample $\beta_j$ from $f(\beta_j | \delta_j = c, \boldsymbol{\theta}_{-\beta_j}, \mathbf{y})$. The categorical random variable $\delta_j$ appears in the likelihood and in its own prior

$$f(\delta_j | \boldsymbol{\pi}) = \prod_{c=1}^{C} \pi_j^{[\delta_j = c]}.$$

Samples can be drawn from this categorical distribution by calculating the membership probabilities

$$\mathbb{P}(\delta_j = c | \boldsymbol{\theta}_{-\delta_j}, \mathbf{y}) = \frac{f(\mathbf{w} | \delta_j = c, \boldsymbol{\theta}, \mathbf{y}) \mathbb{P}(\delta_j = c)}{\sum_{c=1}^{C} f(\mathbf{w} | \delta_j = c, \boldsymbol{\theta}, \mathbf{y}) \mathbb{P}(\delta_j = c)},$$

and then use the sampling routine for a categorical distribution once the probabilities are known. If we would like to use this then if $\delta_j \neq 1$ then we need to integrate out $\beta_j$ from $f_j(\mathbf{w} | \delta_j = c, \boldsymbol{\theta}, \mathbf{y})$, which requires the following integral

$$\begin{aligned} f(\mathbf{w} | \delta_j = c, \boldsymbol{\theta}, \mathbf{y}) &= \int f(\mathbf{w} | \beta_j, \sigma_\varepsilon^2) f(\beta_j | \delta_j, \sigma_\alpha^2) d\beta_j \\ &= \int (2\pi\sigma_\varepsilon^2)^{-n/2} \exp\left[ -\frac{(\mathbf{w} - \mathbf{x}_j\beta_j)'(\mathbf{w} - \mathbf{x}_j\beta_j)}{2\sigma_\varepsilon^2} \right] (2\pi\gamma_c\sigma_\beta^2)^{-1/2} \exp\left[ -\frac{\beta_j^2}{2\gamma_c\sigma_\beta^2} \right] d\beta_j. \end{aligned}$$

Expanding the product terms and combining the exponential terms we have

$$f(\mathbf{w} | \delta_j = c, \boldsymbol{\theta}, \mathbf{y}) = \int (2\pi\gamma_c\sigma_\beta^2)^{-1/2} (2\pi\sigma^2)^{-n/2} \exp\left[ -\frac{1}{2\sigma_\varepsilon^2} \left( \mathbf{w}'\mathbf{w} - 2\mathbf{x}_j'\mathbf{w}\beta_j + \mathbf{x}_j'\mathbf{x}_j\beta_j^2 + \frac{\beta_j^2\sigma_\varepsilon^2}{\gamma_c\sigma_\beta^2} \right) \right] d\beta_j.$$

We let $l_{jc} = \mathbf{x}_j'\mathbf{x}_j + \sigma_e^2 / (\gamma_c\sigma_\beta^2)$ and thus

$$= \int (2\pi\gamma_c\sigma_\beta^2)^{-1/2} (2\pi\sigma_\varepsilon^2)^{-n/2} \exp\left[ -\frac{1}{2\sigma_\varepsilon^2} \left( \mathbf{w}'\mathbf{w} - 2\mathbf{x}_j'\mathbf{w}\beta_j + \beta_j^2 l_{jc} \right) \right] d\beta_j.$$

Letting $\widehat{\beta}_j = \mathbf{x}_j'\mathbf{w}/l_{jc}$ we again look to complete the square and take out from the integral those elements that do not involve $\beta_j$

$$
\begin{aligned}
&= \int (2\pi\gamma_c\sigma_\beta^2)^{-1/2}(2\pi\sigma_\varepsilon^2)^{-n/2}\exp\left[-\frac{1}{2\sigma_e^2}\left(\mathbf{w}'\mathbf{w} - 2\widehat{\beta}_j l_{jc}\beta_j + \beta_j^2 l_{jc} + \widehat{\beta}_j^2 l_{jc} - \widehat{\beta}_i^2 l_{jc}\right)\right] d\beta_j \\
&= (2\pi\frac{\sigma_\varepsilon^2}{l_j})^{1/2}(2\pi\gamma_c\sigma_\beta^2)^{-1/2}(2\pi\sigma_\varepsilon^2)^{-n/2}\exp\left[-\frac{1}{2\sigma_\varepsilon^2}\left(\mathbf{w}'\mathbf{w} - l_{jc}\widehat{\beta}_j^2\right)\right] \times \\
&\quad \int (2\pi\frac{\sigma_\varepsilon^2}{l_{jc}})^{-1/2}\exp\left[-\frac{1}{2\frac{\sigma_\varepsilon^2}{l_{jc}}}(\beta_j - \widehat{\beta}_j)^2\right] d\beta_j.
\end{aligned}
$$

The integral component is now a normal distribution and thus integrates to 1. The term left over after cleaning is

$$
f(\mathbf{w}|\delta_j = c, \boldsymbol{\theta}, \mathbf{y}) = \left(\frac{\gamma_{\delta_j}\sigma_\beta^2 l_j}{\sigma_\varepsilon^2}\right)^{-1/2}(2\pi\sigma_\varepsilon^2)^{-n/2}\exp\left[-\frac{1}{2\sigma_e^2}\left(\mathbf{w}'\mathbf{w} - l_{jc}\widehat{\beta}_j^2\right)\right].
$$

Our goal was to derive a form for $f(\mathbf{w}|\delta_j = c, \boldsymbol{\theta}, \mathbf{y})$ that was independent of $\beta_j$ for use in the probability calculations in

$$
\mathbb{P}(\delta_j = c|\boldsymbol{\theta}_{-\delta_j}, \mathbf{y}) = \frac{f(\mathbf{w}|\delta_j = c, \boldsymbol{\theta}, \mathbf{y})\mathbb{P}(\delta_j = c)}{\sum_{c=1}^{C} f(\mathbf{w}|\delta_j = c, \boldsymbol{\theta}, \mathbf{y})\mathbb{P}(\delta_j = c)}.
$$

We only need to calculate $C - 1$ of these as the $C$th probability is $1 - \sum_{c=1}^{C}\mathbb{P}(\delta_j = c|\boldsymbol{\theta}, \mathbf{y})$. We calculate the probabilities for multiple terms similarly to the following example

$$
\begin{aligned}
\frac{f(\mathbf{w}|\delta_j = 1, \boldsymbol{\theta}, \mathbf{y})}{f(\mathbf{w}|\delta_j = 2, \boldsymbol{\theta}), \mathbf{y})} &= \frac{f(\mathbf{w}|\delta_j = 1, \boldsymbol{\theta}, \mathbf{y})}{\int f(\mathbf{w}|\delta_j = 2, \beta_j, \boldsymbol{\theta}, \mathbf{y}) f(\beta_j|\delta_j, \sigma_\beta^2) d\beta_j} \\
&= \frac{(2\pi\sigma_\varepsilon^2)^{-n/2}\exp\left[-\frac{1}{2\sigma_\varepsilon^2}\left(\mathbf{w}'\mathbf{w}\right)\right]}{\left(\frac{\gamma_2\sigma_\beta^2 l_{j2}}{\sigma_\varepsilon^2}\right)^{-1/2}(2\pi\sigma_\varepsilon^2)^{-n/2}\exp\left[-\frac{1}{2\sigma_\varepsilon^2}\left(\mathbf{w}'\mathbf{w} - l_{j2}\widehat{\beta}_j^2\right)\right]} \\
&= \left(\frac{\gamma_2\sigma_\beta^2 l_{j2}}{\sigma_\varepsilon^2}\right)^{1/2}\exp\left[-\frac{1}{2\sigma_\varepsilon^2}(\mathbf{w}'\mathbf{w}) + \frac{1}{2\sigma_\varepsilon^2}(\mathbf{w}'\mathbf{w}) - \frac{1}{2\sigma_\varepsilon^2}(l_{j2}\widehat{\beta}_j^2)\right] \\
&= \left(\frac{\gamma_2\sigma_\beta^2 l_{j2}}{\sigma_\varepsilon^2}\right)^{1/2}\exp\left[-\frac{1}{2\sigma_\varepsilon^2}(l_{j2}\widehat{\beta}_j^2)\right],
\end{aligned}
$$

where $l_{j2} = [\mathbf{x}_j'\mathbf{x}_j + \sigma_\varepsilon^2/(\gamma_2\sigma_\beta^2)]$ is the left hand side of the MME for $\beta_j$ given $\delta_j = c$, $\widehat{\beta}_j = \mathbf{x}_j'\mathbf{w}/l_{j2}$, and $\mathbf{x}_j'\mathbf{w}$ is the right hand side of the MMEs. This form only depends on the scalar right hand side of the MMEs, which can be updated efficiently, and fixed constants for each iteration.

In the above example we observed a cancelling of the computationally difficult component $\mathbf{w}'\mathbf{w}$, which will happen for all forms of the ratio $\frac{f(\mathbf{w}|\delta_j=c,\boldsymbol{\theta},\mathbf{y})}{f(\mathbf{w}|\delta_j=\sim c,\boldsymbol{\theta}),\mathbf{y})}$. To calculate the probability updates for $\delta_j$ we will attempt to observe the form using $C = 2$ and extrapolate to an arbitrary $C$ value. Therefore,

$$\mathbb{P}(\delta_j = 1|\boldsymbol{\theta},\mathbf{y}) = \frac{f_j(\mathbf{w}|\delta_j = 1,\boldsymbol{\theta},\mathbf{y})\pi_1}{f_j(\mathbf{w}|\delta_j = 1,\boldsymbol{\theta},\mathbf{y})\pi_1 + f_j(\mathbf{w}|\delta_j = 2,\boldsymbol{\theta},\mathbf{y})\pi_2}.$$

Let $\sigma_c^2 = \gamma_c\sigma_\beta^2$

$$\mathbb{P}(\delta_j = 1|\boldsymbol{\theta},\mathbf{y}) = \frac{\left(\frac{\sigma_1^2\mathbf{x}_j'\mathbf{x}_j+\sigma_\varepsilon^2}{\sigma_\varepsilon^2}\right)^{-1/2}(2\pi\sigma_\varepsilon^2)^{-1/2}\exp\left[-\frac{1}{2\sigma_\varepsilon^2}\left(\mathbf{w}'\mathbf{w} - \frac{(\mathbf{x}_j'\mathbf{w})^2}{(\mathbf{x}_j'\mathbf{x}_j+\sigma_\varepsilon^2/\sigma_1^2)}\right)\right]\pi_1}{\left(\frac{\sigma_1^2\mathbf{x}_j'\mathbf{x}_j+\sigma_\varepsilon^2}{\sigma_\varepsilon^2}\right)^{-1/2}(2\pi\sigma_\varepsilon^2)^{-1/2}\exp\left[-\frac{1}{2\sigma_\varepsilon^2}\left(\mathbf{w}'\mathbf{w} - \frac{(\mathbf{x}_j'\mathbf{w})^2}{(\mathbf{x}_j'\mathbf{x}_j+\sigma_\varepsilon^2/\sigma_1^2)}\right)\right]\pi_1 + \left(\frac{\sigma_2^2\mathbf{x}_j'\mathbf{x}_j+\sigma_\varepsilon^2)}{\sigma_\varepsilon^2}\right)^{-1/2}(2\pi\sigma_\varepsilon^2)^{-1/2}\exp\left[-\frac{1}{2\sigma_\varepsilon^2}\left(\mathbf{w}'\mathbf{w} - \frac{(\mathbf{x}_j'\mathbf{w})^2}{(\mathbf{x}_j'\mathbf{x}_j+\sigma_\varepsilon^2/\sigma_2^2)}\right)\right]\pi_2}.$$

The component $(2\pi\sigma_e^2)^{-1/2}\exp\left[-\frac{1}{2\sigma_e^2}(\mathbf{w}'\mathbf{w})\right]$ is common to all the distributions and thus they can be partitioned out in the likelihood calculations. We therefore have

$$\mathbb{P}(\delta_j = 1|\boldsymbol{\theta},\mathbf{y}) = \frac{(\frac{\sigma_1^2(\mathbf{x}_j'\mathbf{x}_j)+\sigma_\varepsilon^2)}{\sigma_\varepsilon^2})^{-1/2}\exp\left[\frac{\sigma_1^2}{2\sigma_\varepsilon^2}\left(\frac{(\mathbf{x}_j'\mathbf{w})^2}{(\sigma_1^2\mathbf{x}_j'\mathbf{x}_j+\sigma_\varepsilon^2)}\right)\right]\pi_1}{(\frac{\sigma_1^2(\mathbf{x}_j'\mathbf{x}_j)+\sigma_\varepsilon^2)}{\sigma_\varepsilon^2})^{-1/2}\exp\left[\frac{\sigma_1^2}{2\sigma_\varepsilon^2}\left(\frac{(\mathbf{x}_j'\mathbf{w})^2}{(\sigma_1^2\mathbf{x}_j'\mathbf{x}_j+\sigma_\varepsilon^2)}\right)\right]\pi_1 + (\frac{\sigma_2^2(\mathbf{x}_j'\mathbf{x}_j)+\sigma_\varepsilon^2)}{\sigma_\varepsilon^2})^{-1/2}\exp\left[\frac{\sigma_2^2}{2\sigma_\varepsilon^2}\left(\frac{(\mathbf{x}_j'\mathbf{w})^2}{(\sigma_2^2\mathbf{x}_j'\mathbf{x}_j+\sigma_\varepsilon^2)}\right)\right]\pi_2}.$$

This form is computationally important as it cancels the $\mathbf{w}'\mathbf{w}$ component so that we do no have to calculate it in each iteration. In order to calculate the posterior probability updates for an arbitrary number of components all that is required is $r_j = \mathbf{x}_j'\mathbf{w}$ and $\sigma_c^2 l_{jc} = \sigma_c^2(\mathbf{x}_j'\mathbf{x}_j) + \sigma_\varepsilon^2$. Then with respect to taking ratios of the components of interest it is sufficient to write

$$\log(\mathcal{L}_c) = \log[f(\mathbf{w}|\delta_j = c,\boldsymbol{\theta})] = -\frac{1}{2}\left[\log\left(\frac{\sigma_c^2 l_{jc}}{\sigma_\varepsilon^2}\right) - \frac{r_j^2}{\sigma_\varepsilon^2 l_{jc}}\right] + \log(\pi_c) \qquad (5)$$

and

$$\mathbb{P}(\delta_j = c|\boldsymbol{\theta}, \mathbf{y}) = \frac{\exp[\log(\mathcal{L}_c)]}{\sum_{c=1}^{C} \exp[\log(\mathcal{L}_c)]},$$

where $C$ is the total number of mixture components. With the mixing proportions $\pi_c$ being sampled in the previous iteration from the Dirichlet distribution. We use the numerically more stable version of the updates from Erbe *et al.*[14]

$$\mathbb{P}(\delta_j = c|\boldsymbol{\theta}, \mathbf{y}) = \frac{1}{\sum_{l=1}^{C} \exp[\log(\mathcal{L}_l) - \log(\mathcal{L}_c)]},$$

which can be shown to be equivalent by

$$\begin{aligned}
\mathbb{P}(\delta_j = c|\boldsymbol{\theta}, \mathbf{y}) &= \frac{\exp[\log(\mathcal{L}_c)]}{\exp[\log(\mathcal{L}_1)] + \exp[\log(\mathcal{L}_2)] + \ldots + \exp[\log(\mathcal{L}_C)]} \\
&= \frac{1}{\frac{\exp[\log(\mathcal{L}_1)]}{\exp[\log(\mathcal{L}_c)]} + \frac{\exp[\log(\mathcal{L}_2)]}{\exp[\log(\mathcal{L}_c)]} + \ldots + \frac{\exp[\log(\mathcal{L}_C)]}{\exp[\log(\mathcal{L}_c)]}} \\
&= \frac{1}{\sum_{l=1}^{C} \exp[\log(\mathcal{L}_l) - \log(\mathcal{L}_c)]}. \qquad (6)
\end{aligned}$$

Combining (6) with the simple calculation of $\log(\mathcal{L}_c)$ using (5) we have all that is required to calculate the probabilities for the categorical distribution for an arbitrary number of mixture components.

Given these probabilities we need to sample from the categorical distribution, which determines which class the variant will be sampled from. With this sampled we can sample the effect from the relevant normal distribution or give it a zero effect. To sample from the categorical distribution we

- Create a vector of cumulative probabilities calculated from above $\mathbb{P}(\delta_j = 1|\boldsymbol{\theta}, \mathbf{y}), \mathbb{P}(\delta_j = 2|\boldsymbol{\theta}, \mathbf{y}), \ldots, \mathbb{P}(\delta_j = C|\boldsymbol{\theta}, \mathbf{y})$ ordered by category
- Accept the lowest $c$ such that the cumulative probability $\geqslant u$, where $u$ is sample form a $U(0, 1)$ distribution. For example if the sampled uniform value is 0.8 and the second group has a cumulative probability of 0.81 then we set $\delta_j = 2$.

Once we have sampled the distribution membership then we can sample the effect from

$$\beta_j|\delta_j = c, \boldsymbol{\theta}, \mathbf{y} \sim N(\mathbf{x}_j'\mathbf{w}/l_{jc}, \sigma_\varepsilon^2/l_{jc}), \tag{7}$$

where $l_{jc} = (\mathbf{x}_j'\mathbf{x}_j + \frac{\sigma_\varepsilon^2}{\sigma_c^2})$.

All that is required to do the sampling is the knowledge of $\mathbf{x}_j'\mathbf{x}_j$, which can be reconstructed from the LD matrix or estimated from the data, and $\mathbf{x}_j'\mathbf{w}$, which is the $j$th element of the right hand side, which can be updated efficiently using residual updating and reconstructed from summary statistics. This will highlighted in the next section.

**Algorithm 1** – Individual level data algorithm

Initialise parameters and read genotypes and phenotypes in PLINK binary format
Initialise $\mathbf{y}^* = \mathbf{y} - \mathbf{X}\boldsymbol{\beta}$
**for** i :=1 **to** number of iterations **do**
  **for** i :=1 **to** $p$ **do**
    Calculate $r_j^* = \mathbf{x}_j'\mathbf{y}^*$
    Calculate $r_j = r_j^* + \mathbf{x}_j'\mathbf{x}_j\beta_j^{(i-1)}$
    Calculate $\sigma_c^2 = \sigma_\beta^2\gamma_{\delta_j=c}$ for each of $C$ classes (e.g., BayesR C=4 and $\boldsymbol{\gamma} = (0, 0.0001, 0.001, 0.01)$)
    Calculate the left hand side $l_{jc} = \mathbf{x}_j'\mathbf{x}_j + \frac{\sigma_\varepsilon^2}{\sigma_c^2}$ for each of the $C$ classes
    Calculate the log densities of given $\delta_j = c$ using $\log(\mathcal{L}_c) = -\frac{1}{2}\left[\log\left(\frac{\sigma_c^2 l_{jc}}{\sigma_\varepsilon^2}\right) - \frac{r_j^2}{\sigma_\varepsilon^2 l_{jc}}\right] + \log(\pi_c)$, where $\pi_c$ is the current
    Calculate the full conditional posterior probability for $\delta_j = c$ for $C$ classes with $\mathbb{P}(\delta_j = c|\boldsymbol{\theta}, \mathbf{y}) = \frac{1}{\sum_{l=1}^{C}\exp[\log(\mathcal{L}_l) - \log(\mathcal{L}_c)]}$
    Using full conditional posterior probabilities sample class membership for $\beta_j^{(i)}$ using categorical random variable sampler
    Given class sample SNP effect $\beta_j^{(i)}$ from $N\left(\frac{r_j}{l_{jc}}, \frac{\sigma_\varepsilon^2}{l_{jc}}\right)$
    Given SNP effect adjust corrected phenotype side $(\mathbf{y}^*)^{(i)} = (\mathbf{y}^*)^{(i-1)} - \mathbf{x}_j\left(\beta_j^{(i)} - \beta_j^{(i-1)}\right)$
  **od**
  Sample update from full conditional for $\sigma_\beta^2$ from scaled inverse chi-squared distribution $\widetilde{\nu}_\beta = \nu_\beta + q$ and $\widetilde{S}^2{}_\beta = \frac{\nu_\beta S_\beta^2 + \sum_{j=1}^{q}\frac{\beta_j^2}{\gamma_c}}{\nu_\beta + q}$, where $q$ is the number of non-zero variants
  Sample update from full conditional for $\sigma_\varepsilon^2$ from scaled inverse chi-squared distribution $\widetilde{\nu}_e = n + \nu_e$ and scale parameter $\widetilde{S}_\varepsilon^2 = \frac{SSE + \nu_\varepsilon S_\varepsilon^2}{n + \nu_\varepsilon}$ and $SSE = \mathbf{y}^{*\prime}\mathbf{y}^*$
  Sample update from full conditional for $\boldsymbol{\pi}$, which is Dirichlet($C, \mathbf{c} + \boldsymbol{\alpha}$), where $\mathbf{c}$ is a vector of length $C$ and contains the counts of the number of variants in each variance class and $\boldsymbol{\alpha} = (1, \ldots, 1)$
  Calculate genetic variance for $h_{SNP}^2$ calculation using $\sigma_g^2 = \mathrm{Var}(\mathbf{X}\boldsymbol{\beta})$
  Calculate $h_{SNP}^2 = \frac{\sigma_g^2}{\sigma_g^2 + \sigma_e^2}$
**od**

#### Summary statistics based Bayesian multiple regression

We relate the phenotype to the set of genetic variants under the multiple linear regression model stated in (1). We can relate the multiple regression model to the the regression coefficients estimated from $p$ simple linear regressions $\mathbf{b}$ from GWAS, by multiplying (1) by $\mathbf{D}^{-1}\mathbf{X}'$ where $\mathbf{D} = \text{diag}(\mathbf{x}_1'\mathbf{x}_1, \ldots, \mathbf{x}_p'\mathbf{x}_p)$ (assuming column centred genotypes) to arrive at

$$\mathbf{D}^{-1}\mathbf{X}'\mathbf{y} = \mathbf{D}^{-1}\mathbf{X}'\mathbf{X}\boldsymbol{\beta} + \mathbf{D}^{-1}\mathbf{X}'\boldsymbol{\varepsilon}. \tag{8}$$

Noting that the correlation matrix between all genetic markers $\mathbf{B} = \mathbf{D}^{-\frac{1}{2}}\mathbf{X}'\mathbf{X}\mathbf{D}^{-\frac{1}{2}}$ we rewrite the multiple regression model as

$$\mathbf{b} = \mathbf{D}^{-\frac{1}{2}}\mathbf{B}\mathbf{D}^{\frac{1}{2}}\boldsymbol{\beta} + \mathbf{D}^{-1}\mathbf{X}'\boldsymbol{\varepsilon}. \tag{9}$$

Assuming $\varepsilon_1, \ldots, \varepsilon_n$ are $N(0, \sigma_\varepsilon)$ the following likelihood can be proposed for the multiple regression coefficients $\boldsymbol{\beta}$

$$\mathcal{L}(\boldsymbol{\beta}; \mathbf{b}, \mathbf{D}, \mathbf{B}) := \mathcal{N}(\mathbf{b}; \mathbf{D}^{-\frac{1}{2}}\mathbf{B}\mathbf{D}^{\frac{1}{2}}\boldsymbol{\beta}, \mathbf{D}^{-\frac{1}{2}}\mathbf{B}\mathbf{D}^{-\frac{1}{2}}), \tag{10}$$

where $\mathcal{N}(\boldsymbol{\xi}; \boldsymbol{\mu}, \boldsymbol{\Sigma})$ represents the multivariate normal distribution with mean vector $\boldsymbol{\mu}$ and covariance matrix $\boldsymbol{\Sigma}$ for $\boldsymbol{\xi}$. If individual level data are available then inference about $\boldsymbol{\beta}$ can be obtained by replacing $\mathbf{D}$ and $\mathbf{B}$ with estimates $(\widehat{\mathbf{D}}, \widehat{\mathbf{B}})$ from the individual level data.

If individual level data are unavailable then we can replace $\mathbf{D}$ with $\widehat{\mathbf{D}} = \text{diag}[2n_1p_1(1 - p_1), \ldots, 2n_jp_j(1 - p_j)]$, where $(n_j, p_j)$ are the sample size used to compute the simple linear regression coefficient and the variant allele frequency respectively. Furthermore, if we assume that genotype column $j$ has been centred and scaled by $\sqrt{2p_j(1 - p_j)}$ then $\widehat{\mathbf{D}} = \text{diag}[n_1, \ldots, n_j]$. These approximations to $\mathbf{D}$, assume the the variant is in Hardy-Weinberg equilibrium, which may not be the true for all variants. Furthermore, summary statistics in the public domain often do not include allele frequencies or report allele frequencies from a reference population. The methodology is susceptible to these deviations from the desired summary statistics for $p_j$, which is the allele frequency for the variant used in the analysis. Motivated by this drawback and the implementation of[3] we seek an approximation to $\mathbf{D}$ that does not depend on $p_j$. For an individual variant from GWAS, the expression for the squared standard error of the estimated effect can be rearranged to arrive at

$$\mathbf{x}_j'\mathbf{x}_j = \frac{(\mathbf{y}'\mathbf{y})_j}{\widehat{\sigma}^2(\mathrm{b}_j)n_j + \mathrm{b}_j^2}. \tag{11}$$

Multiplying top and bottom of the right-hand side by $1/n$

$$\mathbf{x}_j'\mathbf{x}_j = \frac{(\mathbf{y}'\mathbf{y})_j/n_j}{\widehat{\sigma}^2(\mathrm{b}_j) + \mathrm{b}_j^2/n_j}, \tag{12}$$

and we note the numerator is the sample variance of the phenotype assuming it has been centred or a mean term fitting in the marginal regression. It is often the case that GWAS are performed on phenotypes that have been standardised to unit variance, which leads to

$$\mathbf{x}_j'\mathbf{x}_j = \frac{1}{\widehat{\sigma}^2(\mathrm{b}_j) + \mathrm{b}_j^2/n_j}. \tag{13}$$

If this is not the case we note that $\mathbf{y}'\mathbf{y}$ is a constant that can be shown to not contribute to the updates of any parameter in the sampling routine, which was also observed and proven in Mak *et al.*[20]. One further consequence of this assumption is that the parameters are always estimated assuming that the phenotypic variance is unity and thus the genetic variance will be equal to the $h^2_{SNP}$.

Similarly, we replace $\mathbf{B}$, the LD correlation matrix between the genotypes at all markers in the population, which the genotypes in the sample are assumed to be a random sample, with $\widehat{\mathbf{B}}$ an estimate calculated from a population reference that is assumed to closely resemble the sample used to generate the GWAS summary statistics. Zhu and Stephens[3] discuss further the theoretical properties of a similar likelihood and approximate reconstruction.

##### *Sampling $\beta$*

As shown in the previous section the update of the $j$th SNP effect involves the calculation of $r_j = \mathbf{x}_j'\mathbf{w}$ and $l_{jc} = \sigma_c^2(\mathbf{x}_j'\mathbf{x}_j) + \sigma_\varepsilon^2$. We require

$$r_j = \mathbf{x'}_j\mathbf{w} = \mathbf{x'}_j[\mathbf{y} - \mathbf{X}_{-j}\boldsymbol{\beta}_{-j}]$$

for use in (5) and (7). To find this we define the corrected right hand side as $\mathbf{X'y}$ corrected for all current $\boldsymbol{\beta}$

$$\mathbf{r}^* = \mathbf{X'y} - \mathbf{X'X}\boldsymbol{\beta} = \mathbf{X'y} - \mathbf{X'X_{-j}}\boldsymbol{\beta}_{-j} - \mathbf{X'x}_j\beta_j,$$

where $\mathbf{r}^*$ is a vector of dimension $p \times 1$ and $\mathbf{X}_{-j}$ is $\mathbf{X}$ minus the $j$th column. We reconstruct $\mathbf{X'y}$ using the GWAS effect estimates $\mathbf{b}$ and $\mathbf{D}$ such that $\mathbf{X'y} = \mathbf{Db}$. The $j$th element of $\mathbf{r}^*$ is

$$r_j^* = \mathbf{x}_j'\mathbf{y} - \mathbf{x}_j'\mathbf{X}_{-j}\boldsymbol{\beta}_{-j} - \mathbf{x'}_j\mathbf{x}_j\beta_j.$$
$$r_j^* + \mathbf{x'}_j\mathbf{x}_j\beta_j = \mathbf{x}_j'\mathbf{y} - \mathbf{x}_j'\mathbf{X}_{-j}\boldsymbol{\beta}_{-j} = \mathbf{x}_j'(\mathbf{y} - \mathbf{X}_{-j}\boldsymbol{\beta}_{-j}) = r_j.$$

Therefore, for each SNP we calculate

$$r_j = \mathbf{x'}_j\mathbf{w} = r_j^* + \mathbf{x'}_j\mathbf{x}_j\beta_j.$$

This can be used in conjunction with (5) to calculate the class membership probabilities and then update the SNP effect using (7), which only requires $r_j$ and the diagonal elements of $\mathbf{X'X}$. The matrix $\mathbf{X'X}$ is easily calculated from summary statistics via $\mathbf{X'X} = \mathbf{D}^{\frac{1}{2}}\mathbf{B}\mathbf{D}^{\frac{1}{2}}$. Given a new $\beta_j$ in MCMC iteration $m$ we can update the corrected right hand side ($\mathbf{r}^*$) by

$$(\mathbf{r}^*)^{(m+1)} = (\mathbf{r}^*)^{(m)} - \mathbf{X'x}_j[\beta_j^{(m+1)} - \beta_j^{(m)}].$$

This can be shown by noting

$$\begin{aligned}(\mathbf{r}^*) &= \mathbf{X}'\mathbf{y} - \mathbf{X}'\mathbf{X}\boldsymbol{\beta} = \mathbf{X}'\mathbf{y} - \mathbf{X}'\mathbf{X}_{-\mathbf{j}}\boldsymbol{\beta}_{-j} - \mathbf{X}'\mathbf{x}_j\beta_j \\ (\mathbf{r}^*)^{(m)} &= \mathbf{X}'\mathbf{y} - \mathbf{X}'\mathbf{X}_{-\mathbf{j}}\boldsymbol{\beta}_{-j} - \mathbf{X}'\mathbf{x}_j\beta_j^{(m)} \\ (\mathbf{r}^*)^{(m+1)} - (\mathbf{r}^*)^{(m)} &= \mathbf{X}'\mathbf{y} - \mathbf{X}'\mathbf{X}_{-\mathbf{j}}\boldsymbol{\beta}_{-j} - \mathbf{X}'\mathbf{x}_j\beta_j^{(m+1)} - \mathbf{X}'\mathbf{y} + \mathbf{X}'\mathbf{X}_{-\mathbf{j}}\boldsymbol{\beta}_{-j} + \mathbf{X}'\mathbf{x}_j\beta_j^{(m)} \\ (\mathbf{r}^*)^{(m+1)} &= (\mathbf{r}^*)^{(m)} - \mathbf{X}'\mathbf{x}_j(\beta_j^{(m+1)} - \beta_j^{(m)}).\end{aligned}$$

This forms the basis of the RHS updating scheme. The beauty of this updating scheme is that it only requires scalar operations and one vector subtraction. If the effect is 0 for the $j$th SNP no sampling of the effect is required.

The LD matrix only enters into the sampling routine through the diagonal elements in the calculation of $l_{jc}$, which are scalar and efficiently stored, and in the $\mathbf{X}'\mathbf{x}_j$ of the residual update. If we are only updating elements of the $\mathbf{r}^*$ that are in LD with the current SNP then the update is even more efficient as we only have to do the vector subtraction on the non-zero elements. The LD approximation using a sparse matrix or formed by block diagonalising the matrix using a window based approach leads to such a scenario. The efficient updating scheme and the fact that we don't have to store and read the genotype matrix makes this method very efficient. We also avoid the dot product computation of $\mathbf{x}'\mathbf{y}^*$, which was present in the previous BayesR algorithm (Algorithm 1).

###### Sampling $\sigma_\varepsilon^2$

The updating of the $\sigma_\varepsilon^2$ is one of the most critical parts of the summary statistics Gibbs sampling algorithm because its update requires the reconstruction of unobservables, which if we had the full LD matrix could be approximated very well. However, when a sparse LD matrix is used the updating of $\sigma_\varepsilon^2$ can become very unstable due to the approximation.

We begin by deriving the full conditional distribution for $\sigma_\varepsilon^2$. The parameter $\sigma_\varepsilon^2$ appears only in its prior

$$f(\sigma_\varepsilon^2; \nu_\varepsilon, S_\varepsilon^2) = \frac{(S_\varepsilon^2 \nu_\varepsilon/2)^{\nu/2}}{\Gamma(\nu_\varepsilon/2)} \frac{\exp(-\nu_\varepsilon S_\varepsilon^2/2\sigma_\varepsilon^2)}{(\sigma_\varepsilon^2)^{1+\nu_\varepsilon/2}},$$

and in the conditional distribution of $\mathbf{y}$ given all the unknowns,

$$f(\mathbf{y}|\boldsymbol{\theta}) = (2\pi\sigma_\varepsilon^2)^{-n/2} \exp\left[-\frac{(\mathbf{y}-\mathbf{X}\boldsymbol{\beta})'(\mathbf{y}-\mathbf{X}\boldsymbol{\beta})}{2\sigma_\varepsilon^2}\right].$$

Therefore,

$$\begin{aligned} f(\sigma_\varepsilon^2|\boldsymbol{\theta}_{-\sigma_\varepsilon^2}, \mathbf{y}) &= \frac{(S_\varepsilon^2\nu_\varepsilon/2)^{\nu/2}}{\Gamma(\nu_\varepsilon/2)} \frac{\exp(-\nu_\varepsilon S_\varepsilon^2/2\sigma_\varepsilon^2)}{(\sigma_\varepsilon^2)^{1+\nu_\varepsilon/2}} (2\pi\sigma_\varepsilon^2)^{-n/2} \exp\left[-\frac{(\mathbf{y}-\mathbf{X}\boldsymbol{\beta})'(\mathbf{y}-\mathbf{X}\boldsymbol{\beta})}{2\sigma_\varepsilon^2}\right] \\ &\propto \frac{\exp(-\nu_\varepsilon S_\varepsilon^2/2\sigma_\varepsilon^2)}{(\sigma_\varepsilon^2)^{1+\nu_\varepsilon/2}} (\sigma_\varepsilon^2)^{-n/2} \exp\left[-\frac{SSE}{2\sigma_\varepsilon^2}\right] \\ &\propto (\sigma_\varepsilon^2)^{-(n+2+\nu_\varepsilon)/2} \exp\left[-\frac{SSE+\nu_\varepsilon S_\varepsilon^2}{2\sigma_\varepsilon^2}\right], \end{aligned}$$

where $SSE$ is the sum of squared errors of prediction. This can be recognised as the kernel of a scaled inverse chi-square distribution with degrees of freedom $\widetilde{\nu}_\varepsilon = n + \nu_\varepsilon$ and scale parameter $\widetilde{S}_\varepsilon^2 = \frac{SSE+\nu_\varepsilon S_\varepsilon^2}{n+\nu_\varepsilon}$.

If we don't have the individual level data then we cannot observe certain components of the $SSE$. Assuming that the fixed effects have already been corrected from the phenotype then the $SSE$ can be written as

$$\begin{aligned} SSE &= \mathbf{y}'\mathbf{y} - 2(\mathbf{X}\boldsymbol{\beta})'\mathbf{y} + (\mathbf{X}\boldsymbol{\beta})'\mathbf{X}\boldsymbol{\beta} \\ &= \mathbf{y}'\mathbf{y} - 2\boldsymbol{\beta}'\mathbf{D}\mathbf{b} + \boldsymbol{\beta}'\mathbf{X}'\mathbf{X}\boldsymbol{\beta}, \end{aligned} \tag{14}$$

where $\mathbf{D} = \mathrm{diag}(\mathbf{X}'\mathbf{X})$ and $\mathbf{b} = \mathbf{D}^{-1}\mathbf{X}'\mathbf{y}$. We don't observe $\mathbf{y}'\mathbf{y}$ but can we approximate it from the GWAS results. For an individual variant from GWAS, the expression for the squared standard error of the estimated effect can be rearranged to arrive at

$$(\mathbf{y}'\mathbf{y})_j = \widehat{\sigma}^2(\mathrm{b}_j)\mathbf{x}_j'\mathbf{x}_j(n-2) + \mathrm{b}_j^2\mathbf{x}_j'\mathbf{x}_j. \tag{15}$$

Yang *et al.*[21] suggest that the mean across all SNPs is a good estimate of $\mathbf{y}'\mathbf{y}$.

The estimation of $SSE$ highlights the problems with approximating the individual data algorithm using summary data and a sparse $\mathbf{X}'\mathbf{X}$. For example, the $SSE$ equation contains the genetic effect estimates $\mathbf{b}$, which were calculated using the full $\mathbf{X}$ matrix, whereas the

$\mathbf{X}'\mathbf{X}$ is now banded. In the individual data algorithm each marker requires the sampled value of $\sigma_\varepsilon^2$ in each calculation of $l_{jc}$ and the sample of the SNP effect using equation (7). If the full LD matrix is used in the algorithm routine then $SSE$ is the same for each variant. However, as each variant ignores a unique set of LD correlations we propose a marker specific variance $(\sigma_\varepsilon^2)_j$, which attempts to correct for the discrepancy between the fact that a sparse $\mathbf{X}'\mathbf{X}$ has replaced the full $\mathbf{X}'\mathbf{X}$ in (14).

We can attempt to improve on this by estimating a marker specific residual variance that only contains a contribution from non-zero LD correlation matrix elements of $\boldsymbol{\beta}'\mathbf{X}'\mathbf{X}\boldsymbol{\beta}$, which is specific to each variant.

$$(\widehat{\sigma}_\varepsilon^2)_j = \frac{SSE_j}{n_j - 1}. \tag{16}$$

The computation of $\boldsymbol{\beta}'\mathbf{X}'\mathbf{X}\boldsymbol{\beta}$ for each variant is expensive. Therefore, we use the corrected right hand side to efficiently compute the marker specific residual variance. Generally, corrected right hand side $\mathbf{r}^* = \mathbf{X}'\mathbf{y} - \mathbf{X}'\mathbf{X}\boldsymbol{\beta}$ can be combined with (14) for a more efficient update

$$\begin{aligned}
SSE &= \mathbf{y}'\mathbf{y} - 2\boldsymbol{\beta}'\mathbf{X}'\mathbf{y} + \boldsymbol{\beta}'\mathbf{X}'\mathbf{X}\boldsymbol{\beta} \\
&= \mathbf{y}'\mathbf{y} - \boldsymbol{\beta}'\mathbf{X}'\mathbf{y} - \boldsymbol{\beta}'\mathbf{X}'\mathbf{y} + \boldsymbol{\beta}'\mathbf{X}'\mathbf{X}\boldsymbol{\beta} \\
&= \mathbf{y}'\mathbf{y} - \boldsymbol{\beta}'\mathbf{X}'\mathbf{y} - \boldsymbol{\beta}'\mathbf{r}^* && (17) \\
&= \mathbf{y}'\mathbf{y} - \boldsymbol{\beta}'(\mathbf{r}^* + \mathbf{X}'\mathbf{y}) = \mathbf{y}'\mathbf{y} - \sum_{k=1}^{p:\mathrm{LD}\neq 0} \beta_k'[r_k^* + (\mathbf{X}'\mathbf{y})_k] - \sum_{k=1}^{p:\mathrm{LD}=0} \beta_k'[r_k^* + (\mathbf{X}'\mathbf{y})_k]. && (18)
\end{aligned}$$

This allows for an efficient calculation of the per variant marker effects variance

$$SSE_j = (\mathbf{y}'\mathbf{y})_j - \sum_{k=1}^{p:\mathrm{LD}\neq 0} \beta_k'[r_k{}^* + (\mathbf{X}'\mathbf{y})_k],$$

which intuitively is the individual variant total sum of squares subtract the contributions from those other variants in LD with variant $j$. The individual variance is calculated using (16) and used in the per variant calculation of $l_{jc}$ and the sampling of the effect from (7). These values are updated every 100 iterations of the MCMC chain as a compromise between accuracy and computational efficiency. After the completion of the sampling of all marker effects the $SSE$ is calculated using (17) and a global $\sigma_\varepsilon^2$ is sampled from a scaled inverse chi-square distribution with degrees of freedom $\widetilde{\nu}_\varepsilon = n + \nu_\varepsilon$ and scale parameter $\widetilde{S}_\varepsilon^2 = \frac{SSE + \nu_\varepsilon S_\varepsilon^2}{n + \nu_\varepsilon}$. This value is only used for $h^2_{SNP}$ estimation.

##### *Computing genotypic variance*

We are interested in the quantity $\sigma_g^2 = \mathrm{Var}(\mathbf{X}\boldsymbol{\beta})$. This can be approximated by $MSS/n$, where

$$MSS = \boldsymbol{\beta}'\mathbf{X}'\mathbf{X}\boldsymbol{\beta}.$$

Again to efficiently update this we use $\mathbf{r}^* = \mathbf{X}'\mathbf{y} - \mathbf{X}'\mathbf{X}\boldsymbol{\beta} \rightarrow (\mathbf{X}'\mathbf{y} - \mathbf{r}^*) = \mathbf{X}'\mathbf{X}\boldsymbol{\beta}$ and thus

$$\begin{aligned} MSS &= \boldsymbol{\beta}'(\mathbf{X}'\mathbf{y} - \mathbf{r}^*) \\ &= \boldsymbol{\beta}'\mathbf{X}'\mathbf{y} - \boldsymbol{\beta}' r^*. \end{aligned}$$

Therefore, for each iteration of the MCMC chain we calculate the genetic variance using the corrected right hand side, the current sampled $\boldsymbol{\beta}$, $\mathbf{X}'\mathbf{y}$ and

$$\sigma_g^2 = \mathrm{Var}(\mathbf{X}'\boldsymbol{\beta}) = MSS/n.$$

##### *Full-data likelihood equivalence*

If we have access to the individual level data then under the multiple regression model the full-data likelihood for inferring $\boldsymbol{\beta}$ is

$$\mathcal{L}(\boldsymbol{\beta}; \mathbf{y}, \mathbf{X}, \sigma_\varepsilon^2) = (2\pi\sigma_\varepsilon^2)^{-n/2} \exp\left[-\frac{(\mathbf{y}-\mathbf{X}\boldsymbol{\beta})'(\mathbf{y}-\mathbf{X}\boldsymbol{\beta})}{2\sigma_\varepsilon^2}\right]. \tag{19}$$

Zhu and Stephens[3] show that under their likelihood and the assumptions that the LD correlation matrix $\widehat{\mathbf{B}}$ has been computed from the genotypes $\mathbf{X}$, $n > p$ and that $\sigma_\varepsilon^2 = n^{-1}\mathbf{y}'\mathbf{y}$ that the full-data likelihood is equivalent to their likelihood up to a constant that does not depend on $\boldsymbol{\beta}$. We will seek to arrive at the same conclusion under the likelihood proposed in equation (10). Replacing in (10) $\mathbf{B}$ with $\widehat{\mathbf{B}}$ and $\mathbf{D}$ with $\widehat{\mathbf{D}}$ then the summary data likelihood is

$$\mathcal{L}(\boldsymbol{\beta}; \mathbf{b}, \widehat{\mathbf{B}}, \widehat{\mathbf{D}}) = |2\pi\sigma_\varepsilon^2\widehat{\mathbf{D}}^{-1/2}\widehat{\mathbf{B}}\widehat{\mathbf{D}}^{-1/2}|^{-1/2} \times \exp\left[-\frac{(\mathbf{b}-\widehat{\mathbf{D}}^{-1/2}\widehat{\mathbf{B}}\widehat{\mathbf{D}}^{1/2}\boldsymbol{\beta})'(\widehat{\mathbf{D}}^{-1/2}\widehat{\mathbf{B}}\widehat{\mathbf{D}}^{-1/2})^{-1}(\mathbf{b}-\widehat{\mathbf{D}}^{-1/2}\widehat{\mathbf{B}}\widehat{\mathbf{D}}^{1/2}\boldsymbol{\beta})}{2\sigma_\varepsilon^2}\right].$$

When $n > p$ then $\widehat{\mathbf{B}}$ computed from the sample genotypes is non-singular and thus we require this assumption in the statement of the likelihood. Taking the logarithm and expanding we have

$$\log[\mathcal{L}(\boldsymbol{\beta}; \mathbf{b}, \widehat{\mathbf{B}}, \widehat{\mathbf{D}})] = -\frac{p}{2}\log(2\pi\sigma_\varepsilon^2) - \frac{1}{2}\log|\widehat{\mathbf{D}}^{-1/2}\widehat{\mathbf{B}}\widehat{\mathbf{D}}^{-1/2}| - \frac{1}{2\sigma_e^2}\mathbf{b}'(\widehat{\mathbf{D}}^{-1/2}\widehat{\mathbf{B}}\widehat{\mathbf{D}}^{-1/2})^{-1}\mathbf{b} + \frac{1}{\sigma_e^2}(\widehat{\mathbf{D}}^{-1/2}\widehat{\mathbf{B}}\widehat{\mathbf{D}}^{1/2}\boldsymbol{\beta})'(\widehat{\mathbf{D}}^{-1/2}\widehat{\mathbf{B}}\widehat{\mathbf{D}}^{-1/2})^{-1}\mathbf{b} - \frac{1}{2\sigma_\varepsilon^2}(\widehat{\mathbf{D}}^{-1/2}\widehat{\mathbf{B}}\widehat{\mathbf{D}}^{1/2}\boldsymbol{\beta})'(\widehat{\mathbf{D}}^{-1/2}\widehat{\mathbf{B}}\widehat{\mathbf{D}}^{-1/2})^{-1}(\widehat{\mathbf{D}}^{-1/2}\widehat{\mathbf{B}}\widehat{\mathbf{D}}^{1/2}\boldsymbol{\beta}).$$

Similarly for the full data likelihood, we take the logarithm and expand

$$\log[\mathcal{L}(\boldsymbol{\beta};\mathbf{y},\mathbf{X},\sigma_\varepsilon^2)] = -\frac{n}{2}\log(2\pi\sigma_\varepsilon^2) - \frac{1}{2\sigma_\varepsilon^2}\mathbf{y}'\mathbf{y} + \frac{1}{\sigma_\varepsilon^2}\boldsymbol{\beta}'\mathbf{X}'\mathbf{y} - \frac{1}{2\sigma_\varepsilon^2}\boldsymbol{\beta}'\mathbf{X}'\mathbf{X}\boldsymbol{\beta}. \tag{20}$$

Looking at the difference we have

$$\log[\mathcal{L}(\boldsymbol{\beta};\mathbf{y},\mathbf{X},\sigma_\varepsilon^2)] - \log[\mathcal{L}(\boldsymbol{\beta};\mathbf{b},\widehat{\mathbf{B}},\widehat{\mathbf{D}})] = -\frac{n}{2}\log(2\pi\sigma_\varepsilon^2) - \frac{1}{2\sigma_\varepsilon^2}\mathbf{y}'\mathbf{y} + \frac{1}{\sigma_\varepsilon^2}\boldsymbol{\beta}'\mathbf{X}'\mathbf{y} - \frac{1}{2\sigma_\varepsilon^2}\boldsymbol{\beta}'\mathbf{X}'\mathbf{X}\boldsymbol{\beta} -$$
$$\frac{p}{2}\log(2\pi\sigma_\varepsilon^2) + \frac{1}{2}\log|\widehat{\mathbf{D}}^{-1/2}\widehat{\mathbf{B}}\widehat{\mathbf{D}}^{-1/2}| \ + \frac{1}{2\sigma_\varepsilon^2}\mathbf{b}'(\widehat{\mathbf{D}}^{-1/2}\widehat{\mathbf{B}}\widehat{\mathbf{D}}^{-1/2})^{-1}\mathbf{b} \ -$$
$$\frac{1}{\sigma_\varepsilon^2}(\widehat{\mathbf{D}}^{-1/2}\widehat{\mathbf{B}}\widehat{\mathbf{D}}^{1/2}\boldsymbol{\beta})'(\widehat{\mathbf{D}}^{-1/2}\widehat{\mathbf{B}}\widehat{\mathbf{D}}^{-1/2})^{-1}\mathbf{b} \ + \frac{1}{2\sigma_\varepsilon^2}(\widehat{\mathbf{D}}^{-1/2}\widehat{\mathbf{B}}\widehat{\mathbf{D}}^{1/2}\boldsymbol{\beta})'(\widehat{\mathbf{D}}^{-1/2}\widehat{\mathbf{B}}\widehat{\mathbf{D}}^{-1/2})^{-1}(\widehat{\mathbf{D}}^{-1/2}\widehat{\mathbf{B}}\widehat{\mathbf{D}}^{1/2}\boldsymbol{\beta}).$$

Gathering up terms that do not depend on $\boldsymbol{\beta}$ and letting them equal $Q$, and substituting $\widehat{\mathbf{D}}^{-1/2}\mathbf{X}'\mathbf{X}\widehat{\mathbf{D}}^{-1/2} = \widehat{\mathbf{B}}$, $\mathbf{b} = \widehat{\mathbf{D}}^{-1}\mathbf{X}'\mathbf{y}$ and $(\widehat{\mathbf{D}}^{-1}\mathbf{X}'\mathbf{X}\widehat{\mathbf{D}}^{-1})^{-1} = \widehat{\mathbf{D}}(\mathbf{X}'\mathbf{X})^{-1}\widehat{\mathbf{D}}$ then the difference is

$$\begin{aligned}
&= Q + \frac{1}{\sigma_\varepsilon^2}\boldsymbol{\beta}'\mathbf{X}'\mathbf{y} - \frac{1}{\sigma_\varepsilon^2}(\widehat{\mathbf{D}}^{-1}\mathbf{X}'\mathbf{X}\boldsymbol{\beta})'(\widehat{\mathbf{D}}^{-1}\mathbf{X}'\mathbf{X}\widehat{\mathbf{D}}.^{-1})^{-1}\mathbf{b} \\
&\quad - \frac{1}{2\sigma_\varepsilon^2}\boldsymbol{\beta}'\mathbf{X}'\mathbf{X}\boldsymbol{\beta} \ + \frac{1}{2\sigma_\varepsilon^2}(\widehat{\mathbf{D}}^{-1}\mathbf{X}'\mathbf{X}\boldsymbol{\beta})'(\widehat{\mathbf{D}}^{-1}\mathbf{X}'\mathbf{X}\widehat{\mathbf{D}}^{-1})^{-1}(\widehat{\mathbf{D}}^{-1}\mathbf{X}'\mathbf{X}\boldsymbol{\beta}) \\
&= Q + \frac{1}{\sigma_\varepsilon^2}\boldsymbol{\beta}'\mathbf{X}'\mathbf{y} - \frac{1}{\sigma_\varepsilon^2}(\widehat{\mathbf{D}}^{-1}\mathbf{X}'\mathbf{X}\boldsymbol{\beta})'\widehat{\mathbf{D}}(\mathbf{X}'\mathbf{X})^{-1}\widehat{\mathbf{D}}\widehat{\mathbf{D}}^{-1}\mathbf{X}'\mathbf{y} \ + \\
&\quad - \frac{1}{2\sigma_\varepsilon^2}\boldsymbol{\beta}'\mathbf{X}'\mathbf{X}\boldsymbol{\beta} + \frac{1}{2\sigma_\varepsilon^2}(\widehat{\mathbf{D}}^{-1}\mathbf{X}'\mathbf{X}\boldsymbol{\beta})'\widehat{\mathbf{D}}(\mathbf{X}'\mathbf{X})^{-1}\widehat{\mathbf{D}}(\widehat{\mathbf{D}}^{-1}\mathbf{X}'\mathbf{X}\boldsymbol{\beta}) \\
&= Q.
\end{aligned}$$

Given this the summary and individual data models will be equivalent up to a constant $Q = -\frac{n}{2}\log(2\pi\sigma_\varepsilon^2) + \frac{p}{2}\log(2\pi\sigma_\varepsilon^2) + \frac{1}{2}\log|\widehat{\mathbf{D}}^{-1/2}\widehat{\mathbf{B}}\widehat{\mathbf{D}}^{-1/2}|$, as the $-\frac{1}{2\sigma_\varepsilon^2}\mathbf{y}'\mathbf{y}$ and $\frac{1}{2\sigma_\varepsilon^2}\mathbf{b}'(\widehat{\mathbf{D}}^{-1/2}\widehat{\mathbf{B}}\widehat{\mathbf{D}}^{-1/2})^{-1}\mathbf{b}$ terms cancel, that does not depend on $\boldsymbol{\beta}$. This assumes that $\sigma_\varepsilon^2$ is known and the full LD matrix is computed from the individual data genotypes. In reality we do not know $\sigma_\varepsilon^2$ but estimate it using posterior inference and the MCMC algorithm. The difference between the individual and summary likelihoods in dependent on $\sigma_\varepsilon^2$ with the deviation dependent on the difference between $n$ and $p$.

#### Method implementation

These summary based Bayesian multiple regression method has been implemented in a software tool named Genome-wide Complex Trait Bayesian analyses (GCTB). The tool has been written in the C++ programming language and is available and freely distributable under a MIT License. The method requires the following data:

- The univariate regression effects from GWAS $\mathbf{b}$.
- The standard error estimates for each genetic effect from univariate regression $\widehat{\sigma}^2(\mathbf{b})$.
- An LD matrix calculated from the cohort or a population matched reference $\mathbf{B} = \mathbf{D}^{-1/2}\mathbf{X}'\mathbf{X}\mathbf{D}^{-1/2}$, where $\mathbf{X}$ is the genotype matrix from the cohort to analysed or a reference. If we assume that the SNP covariates have been mean adjusted, or the mean has been fitted in the univariate regression analysis, then $\mathbf{D}$ is a diagonal matrix with diagonal elements that should be very well approximated in large samples by equation (13). If the genotypes are assumed to have been centred and scaled then $\mathbf{D}$ is a diagonal matrix with diagonal elements $n_j$. The algorithm requires $\mathbf{X}'\mathbf{X}$ and thus using effects estimated from a PLINK GWAS we have $\mathbf{X}'\mathbf{X} = \mathbf{D}^{1/2}\mathbf{B}\mathbf{D}^{1/2}$.
- The algorithm also requires $\mathbf{X}'\mathbf{y}$, which from the least squares solutions can be recovered $\widehat{\mathbf{b}} = \text{diag}(\mathbf{X}'\mathbf{X})^{-1}\mathbf{X}'\mathbf{y} = \mathbf{D}^{-1}\mathbf{X}'\mathbf{y}$ and thus $\mathbf{X}'\mathbf{y} = \mathbf{D}\widehat{\mathbf{b}}$.

##### Parameters to be estimated

- The joint genetic effects $\boldsymbol{\beta}_0$. Initialised as all zeros.
- The proportion of effects in each class $\boldsymbol{\pi}$. This is by default the original BayesR model with four components and initialised such that a large proportion of variants have no effect, for example, $\boldsymbol{\pi}_0 = (0.95, 0.02, 0.02, 0.01)$.
- The vector $\gamma$ specifies the weights for the mixture normal class variances, which are by default set to $(0, 0.01, 0.1, 1)$. These deviate from the BayesR model $(0, 0.0001, 0.001, 0.01)'$ as they represent the weights for the marker effect variance as opposed to the genetic variance as in Erbe *et al.*[14] and Moser *et al.*[2].
- The variance components include the marker effect variance $\sigma^2_\beta$ and the residual

variance $\sigma_\varepsilon^2$. The initial value of the residual variance is set to $(\sigma_\varepsilon^2)_0 = \frac{SSE}{n-1}$, where $SSE = \mathbf{y}'\mathbf{y} - 2\boldsymbol{\beta}_0'\mathbf{X}'\mathbf{y} + \boldsymbol{\beta}_0'\mathbf{X}'\mathbf{X}\boldsymbol{\beta}_0 = \mathbf{y}'\mathbf{y}$ (given $\boldsymbol{\beta}_0 = \mathbf{0}$), and $\mathbf{y}'\mathbf{y} = \frac{1}{p}\sum_{j=1}^{p}(\mathbf{y}'\mathbf{y})_j$ where $(\mathbf{y}'\mathbf{y})_j = \widehat{\sigma}^2(\widehat{b}_j)\mathbf{x}_j'\mathbf{x}_j(n-2) + \widehat{b}_j^2\mathbf{x}_j'\mathbf{x}_j$, which is reconstructed from the summary statistics from the univariate regression for each variant. The parameter $\sigma_\beta^2$ is initialised as $(\sigma_\beta^2)_0 = (\sigma_g^2)_0/[(1-(\boldsymbol{\pi}_0)_1)\sum_j 2p_j(1-p_j)]$[22], where we use the double notation and $p_j$ is the allele frequency of allele $j$ $(\sigma_g^2)_0$ is the genotypic variance and is set to $(\sigma_g^2)_0 = h_{SNP}^2\frac{\mathbf{y}'\mathbf{y}}{(n-1)}$ and $h_{SNP}^2 = 0.5$ is set by default set a starting $h_{SNP}^2$.

#### Hyperparameters to be set

- The degrees of freedom $\nu$ and scale parameters $S^2$ for the scale inverse chi-squared distribution, which form the priors for the $\sigma_\beta^2$ and $\sigma_\varepsilon^2$ parameters are required to be set. For both the degrees of freedom are set to 4[17] and $S_\alpha^2 = \frac{(\nu_\beta-2)(\sigma_\beta^2)_0}{\nu_\beta} = \frac{(\sigma_\beta^2)_0}{2}$ and $S_\varepsilon^2 = \frac{(\nu_\varepsilon-2)(\sigma_\varepsilon^2)_0}{\nu_\varepsilon} = \frac{(\sigma_\varepsilon^2)_0}{2}$, which comes from a method of moments estimator for the scale parameter.

**Algorithm 2** Summary data algorithm

Initialise parameters and read summary statistics
Reconstruct $\mathbf{X'X}$ and $\mathbf{X'y}$ from summary statistics and LD reference panel
Calculate $\mathbf{r}^* = \mathbf{X'y} - \mathbf{X'X}\boldsymbol{\beta}$
**for** i :=1 **to** number of iterations **do**
  **for** i :=1 **to** $p$ **do**
    Calculate $\mathbf{r}_j = \mathbf{r}_j^* + \mathbf{x}_j'\mathbf{x}_j\beta_j$
    Calculate $\sigma_c^2 = \sigma_\alpha^2\gamma_{\delta_j=c}$ for each fo $C$ classes (e.g., SBayesR C=4 and $\boldsymbol{\gamma} = (0, 0.01, 0.1, 1)'$)
    Calculate the left hand side $l_{jc} = \mathbf{x}_j'\mathbf{x}_j + \frac{\sigma_e^2}{\sigma_c^2}$ for each of the $C$ classes
    Calculate the log densities of given $\delta_j = c$ using $\log(\mathcal{L}_c) = -\frac{1}{2}\left[\log\left(\frac{\sigma_c^2 l_{jc}}{\sigma_e^2}\right) - \frac{r_j^2}{\sigma_e^2 l_{jc}}\right] + \log(\pi_c)$, where $\pi_c$ is the current
    Calculate the full conditional posterior probability for $\delta_j = c$ for $C$ classes with $\mathbb{P}(\delta_j = c|\boldsymbol{\theta}, \mathbf{y}) = \frac{1}{\sum_{l=1}^{C}\exp[\log(\mathcal{L}_l) - \log(\mathcal{L}_c)]}$
    Using full conditional posterior probabilities sample class membership for $\beta_j^{(i)}$ using categorical random variable sampler
    Given class sample SNP effect $\beta_j^{(i)}$ from $N\left(\frac{\mathbf{r}_j}{l_{jc}}, \frac{\sigma_e^2}{l_{jc}}\right)$
    Given SNP effect adjust corrected right hand side $(\mathbf{r}^*)^{(i+1)} = (\mathbf{r}^*)^{(i)} - \mathbf{X}'\mathbf{x}_j\left(\beta_j^{(i+1)} - \beta_j^{(i)}\right)$. $\mathbf{X}'\mathbf{x}_j$ is the $j$th column of $\mathbf{X'X}$.
  **od**
  Sample update from full conditional for $\sigma_\alpha^2$ from scaled inverse chi-squared distribution $\widetilde{\nu}_\alpha = \nu_0 + q$ and $\widetilde{\tau}^2{}_\alpha = \frac{\nu_0\tau_0^2 + \sum_{j=1}^{q}\frac{\beta_j^2}{\gamma_{\delta_j}}}{\nu_0 + q}$, where $q$ is the number of non-zero variants
  Sample update from full conditional for $\sigma_\varepsilon^2$ from scaled inverse chi-squared distribution $\widetilde{\nu}_e = n + \nu_e$ and scale parameter $\widetilde{\tau}_e^2 = \frac{SSE + \nu_e\tau_e^2}{n + \nu_e}$ and $SSE = \mathbf{y'y} - \boldsymbol{\beta}'\mathbf{r}^* - \boldsymbol{\beta}'\mathbf{X'y}$
  Sample update from full conditional for $\boldsymbol{\pi}$, which is Dirichlet($C, \mathbf{c} + \boldsymbol{\alpha}$), where $\mathbf{c}$ is a vector of length $C$ and contains the counts of the number of variants in each variance class.
  Calculate genetic variance for $h_{SNP}^2$ calculation using $\sigma_g^2 = MSS/n$, where $MSS = \widehat{\boldsymbol{\beta}}'\mathbf{X'y} - \widehat{\boldsymbol{\beta}}'r^*$
  Calculate $h_{SNP}^2 = \frac{\sigma_g^2}{\sigma_g^2 + \sigma_e^2}$
**od**
